# Class I Histone Deacetylases (HDAC1–3) are Histone Lysine Delactylases

**DOI:** 10.1101/2021.03.24.436780

**Authors:** Carlos Moreno-Yruela, Di Zhang, Wei Wei, Michael Bæk, Jinjun Gao, Alexander L. Nielsen, Julie E. Bolding, Lu Yang, Samuel T. Jameson, Jiemin Wong, Christian A. Olsen, Yingming Zhao

## Abstract

Lysine l-lactylation [K(l-la)] is a newly discovered histone mark that can be stimulated under conditions of high glycolysis, such as the Warburg effect. K(l-la) is associated with functions that are different from the widely studied histone acetylation. While K(l-la) can be introduced by the acetyltransferase p300, histone delactylase enzymes remain unknown. Here, we report the systematic evaluation of zinc- and NAD^+^-dependent HDACs for their ability to cleave ε-*N*-l-lactyllysine marks. Our screens identified HDACs 1–3 and SIRT1–3 as delactylases in vitro. HDACs 1–3 show robust activity toward not only K(l-la) but also K(d-la) and diverse short-chain acyl modifications. We further confirmed the de-l-lactylase activity of HDACs 1 and 3 in cells. Identification of p300 and HDAC3 as regulatory enzymes suggests that histone lactylation is installed and removed by enzymes as opposed to spontaneous chemical reactivity. Our results therefore represent an important step toward full characterization of this pathway’s regulatory elements.

## Introduction

Emerging lines of evidence suggest that metabolic end products and intermediates can have signaling functions in addition to their cognate functions. A metabolite can exert its function covalently through intrinsic chemical reactivity (*1, 2*) or enzyme-catalyzed reactions (*3, 4*). Classic examples of the latter include acetyl-coenzyme A (CoA) and *S*-adenosylmethionine (SAM), which can be used by acetyltransferases for lysine acetylation and by methyltransferases for lysine methylation, respectively (*3, 5*). Other metabolites, such as nicotinamide adenine dinucleotide (NAD^+^) and α-ketoglutarate, can serve as cofactors and regulate the activities of corresponding deacetylases and demethylases (*6*). l-Lactate, traditionally known as a metabolic waste product, has recently been found to play important roles in metabolism. l-Lactate production regenerates the NAD^+^ consumed by glycolysis within cells. The shuttling of l-lactate between different organs and cells serves as a major circulating carbohydrate source that plays important roles in normal physiology as well as in cancer (*7–9*). l-Lactate is massively induced under hypoxia and during the Warburg effect (*10–12*), which are associated with many cellular processes and are closely linked to diverse diseases including neoplasia, sepsis, and autoimmune diseases (*13*). Nevertheless, the non-metabolic functions of l-lactate in physiology and disease, especially during Warburg effect, remain largely unknown.

We recently reported that l-lactate is a precursor that can label and stimulate histone lysine ε-*N*-l-lactylation [K(l-la)] (*14*). Data suggest that l-lactate is transformed into l-lactyl-CoA (*15*) and transferred onto histones by acetyltransferases such as p300 (Fig. 1A) (*14*). Therefore, like acetyl-CoA and histone lysine acetylation (Kac) (*16*), histone K(l-la) represents another example showing that acyl-CoA species can affect gene expression directly via histone posttranslational modification (PTM). In addition, histone K(l-la) has different kinetics from those of histone (Kac) during glycolysis (*14*). Histone K(l-la) is induced by hypoxia and the Warburg effect, serving as a feedback mechanism to promote homeostatic gene expression in the late phase of macrophage polarization (*14*). Our data therefore indicate that histone K(l-la) is a physiologically relevant histone mark and has unique biological functions.

**Fig. 1.**
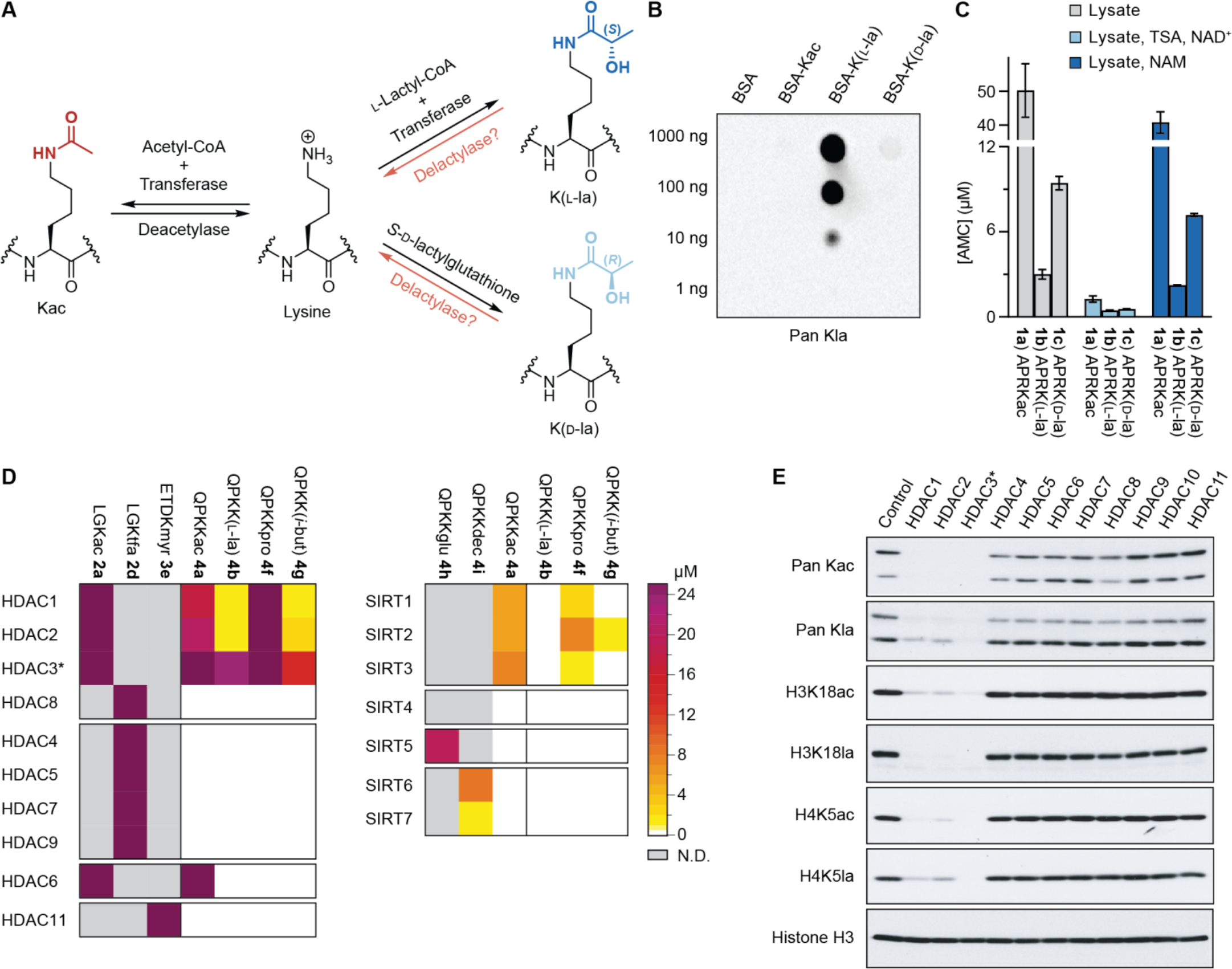
In vitro screening for delactylase activity. (**A**) Reversible post-translational modification of lysine to ε-*N*-acetyllysine (Kac), ε-*N*-l-lactyllysine (K(l-la)) and ε-*N*-d-lactyllysine (K(d-la)). (**B**) Dot blot assay using the pan anti-Kla antibody and bovine serum albumin (BSA) that were unmodified or modified with Kac, K(l-la) or K(d-la) acyl modifications. (**C**) Conversion of 7-amino-4-methylcoumarin (AMC)-conjugated Kac (**1a**), K(l-la) (**1b**) and K(d-la) (**1c**) substrates (see fig. S2 for structures) by HEK293T whole-cell lysates, and inhibition with an HDAC inhibitor (trichostatin A, TSA) or a pan-sirtuin inhibitor (nicotinamide, NAM). Data represent mean ± SD, *n* = 4. (**D**) Deacylase activity screening using short AMC-conjugated peptides, including positive controls for each recombinant enzyme (left part of heat maps), a K(l-la) substrate (**4b**), and substrates bearing short aliphatic modifications (**4f**, **4g**). Heat maps represent mean values of AMC concentration, *n* = 2 (see fig. S2 for substrate structures and fig. S3 for bar graphs). (**E**) Deacylase activity screening using core histones from HeLa cells and antibodies against Kac and K(l-la) modifications, with histone H3 as loading control (4 h reaction, see fig. S1A for data after 1 h incubation). *HDAC3 incubated with the deacetylase activation domain (DAD) of NCoR2.

In addition to l-lactate (with (*S*) configuration), its structural isomer d-lactate (with (*R*) configuration) is also found in cells, although at much lower concentration (nM concentration, compared to typical mM concentration of l-lactate, which can reach up to 40 mM in some cancer cells) (*17, 18*). d-Lactate is formed primarily from methylglyoxal (MGO) through the glyoxalase pathway (*19, 20*) and is overproduced in rare conditions, including certain cases of short bowel syndrome. Under these unusual circumstances, d-lactate can reach 3 mM concentration or higher in plasma (*18, 21*). *S*-d-(*R*)-Lactylglutathione, an intermediate in the glyoxalase pathway, can react non-enzymatically to install K(d-la) PTMs on glycolytic enzymes (Fig. 1A) (*22*). Nevertheless, five lines of evidence suggest that histones are l-lactylated rather than d-lactylated: (1) the huge difference in cellular concentration of the precursor metabolites, l-lactate vs d-lactate, (2) the specificity of the antibodies used in our work toward K(l-la) (~100-fold or higher, Fig. 1B), (3) the fact that histone lactylation can be stimulated and labeled by l-lactate (*14*), (4) the specific genomic localization of histone K(l-la) that excludes the possibility of random, spontaneous chemical installation (*14*), and (5) the fact that K(d-la) is enriched solely in cytosolic proteins that are in close contact with *S*-d-lactylglutathione (*22*), which is not the case for histones. Despite the progress, key regulatory mechanisms of histone K(l-la) remain unknown, including the enzymes that remove this modification in the cell (Fig. 1A).

Mammals express two families of lysine deacylases with 18 enzymes in total. Histone deacetylases (HDAC1–11, grouped into classes I, II and IV) are Zn^2+^-dependent (*23*), while the sirtuins (SIRT1–7, class III HDACs) are dependent on NAD^+^ as co-substrate (*24*). Among these, several isozymes exhibit preferential enzymatic activities against non-Kac acylations. For example, SIRT5 is an efficient lysine demalonylase, desuccinylase, and deglutarylase but not deacetylase (*25–28*), and HDAC8, HDAC11, SIRT2, and SIRT6 harbor substantial activities against long chain acyl modifications (*29–33*). HDACs 1–3, in addition to their efficient deacetylase activity, also remove ε-*N*-crotonyllysine (Kcr) and ε-*N*-d-β-hydroxybutyryllysine [K(d-bhb)] PTMs (*34–37*). Therefore, it is likely that enzymes from the HDAC classes may be able to catalyze the removal of histone lactyl modifications.

Here, we report a screening of all 18 HDACs for potential delactylase activities using fluorophore-coupled peptides, fluorophore-free histone peptides, and extracted histones as substrates. We show that class I HDACs 1–3 are the most efficient lysine delactylases in vitro, and that HDAC1 and HDAC3 have delactylase activity in cells. These findings support that histone K(l-la) modification is a regulatory and dynamic epigenetic mechanism. Interestingly, HDACs 1–3 harbor activities in vitro toward diverse acyllysine groups including aliphatic and hydroxylated modifications of 2–5 carbons. In addition, even though we detect minor delactylase activity by SIRT1–3 in vitro, the pan-sirtuin inhibitor nicotinamide does not affect the delactylase activity of a cell lysate or the histone K(l-la) levels in cells. Our data suggest HDACs 1 and 3 are the main delactylases in the cell.

## Results

### Delactylase activities of Zn^2+^- and NAD^+^-dependent human HDACs

#### Delactylase activities of cell lysates

Lactylation of the side chains of lysine residues has been found on histones and other proteins (*14, 22*), but no efficient hydrolytic enzyme has been identified for this modification so far. To address this question, we first examined whether enzymes in the lysate of a human cell line (HEK293T) were able to hydrolyze ε-*N*-lactyllysine modifications from short fluorophore-coupled peptide sequences derived from histone 3 [H3_15–18_: Ac-APRK(l-la)-AMC (**1b**) and Ac-APRK(d-la)-AMC (**1c**)]. The whole cell lysate was indeed able to remove the lactyl modifications; albeit, to a lower extent than the corresponding Kac modification (**1a**) (Fig. 1C). Treatment with the broad spectrum HDAC inhibitor trichostatin A (TSA) (*38, 39*) or the pan-sirtuin inhibitor nicotinamide (NAM) (*40, 41*) indicated that the Zn^2+^-dependent HDACs are more likely to be responsible for these activities, because inhibition of sirtuin activity with NAM did not substantially change the outcome of the deacylation events (Fig. 1C).

#### Delactylase activities of recombinant zinc-dependent HDACs and sirtuins toward peptide substrates

Encouraged by these initial results, we then performed a screen of the activities of recombinant HDAC enzymes against known fluorogenic control substrates for each isozyme (*31, 39, 42, 43*) together with a set of fluorophore conjugated p53317-320 peptides (Ac-QPKK-AMC) featuring small acyl-lysine modifications (Fig. 1D). Class I HDACs 1–3 and sirtuins SIRT1–3 cleaved the aliphatic ε-*N*-propionyllysine (Kpro, **4f**) and, to some extent, ε-*N*-isobutyryllysine (K(*i*-but), **4g**) modifications (*31, 44*). However, only HDACs 1–3 exhibited measurable activity against the K(l-la) substrate (**4b**) in these assays.

#### Delactylase activities of recombinant zinc-dependent HDACs and sirtuins toward histone substrates

To corroborate the results, we carried out in vitro delactylation assays, using 18 recombinant HDACs, and core histones as substrates, followed by Western blot analysis. We first evaluated specificities of anti-l-lactyllysine antibodies, using bovine serum albumin (BSA) preparations that were either unmodified, acetylated, l-lactylated, or d-lactylated. We showed that the anti-l-lactyllysine antibody has more than 100-fold higher affinity for K(l-la)-modified than K(d-la)-modified BSA and ~1,000-fold selectivity versus Kac-modified BSA (Fig. 1B). Thus, in this paper, unless specified, all the Kla as well as lactylation will suggest the l-isomer instead of the d-isomer or their mixture.

Our enzymatic results indicated that HDACs 1–3 substantially decrease the overall Kac and Kla levels on histones, and also drastically reduce the levels of H3K18 and H4K5 l-lactylation (Fig. 1E and fig. S1A). In addition, we detected robust l-delactylation by SIRT1–3 to overall histone lactylation as well as H3K18(l-la) and H4K5(l-la) (fig. S1B,C).

Together, our preliminary data showed that HDACs 1–3, and to some extent SIRT1–3, have robust in vitro delactylase activity and that HDACs 1–3 are the most efficient enzymes among the 18 human recombinant enzymes in vitro. Thus, HDACs 1–3 were selected for further investigation for their delactylase activity in vitro and in cells.

### HDAC3 is the most efficient eraser of both l- and d-lactyl-lysine in vitro

To take a further look at the delactylase activity of HDACs 1–3, we decided to investigate their delactylase activities compared to deacetylation. To this end, we synthesized and tested additional fluorogenic, histone-based substrates (H4_10-12_: Ac-LGK-AMC and H3_6-9_: Ac-TARK-AMC) with the Kac, K(l-la), and K(d-la) modifications (please consult Supplementary Methods for synthetic details). The activity of HDACs 1 and 2 caused limited conversion of the lactylated substrates compared to the acetylated counterparts and showed a slight preference for K(d-la) over K(l-la). HDAC3, on the other hand, converted l- and d-lactylated substrates to an extent more similar to that of acetylated substrates (Fig. 2A).

**Fig. 2.**
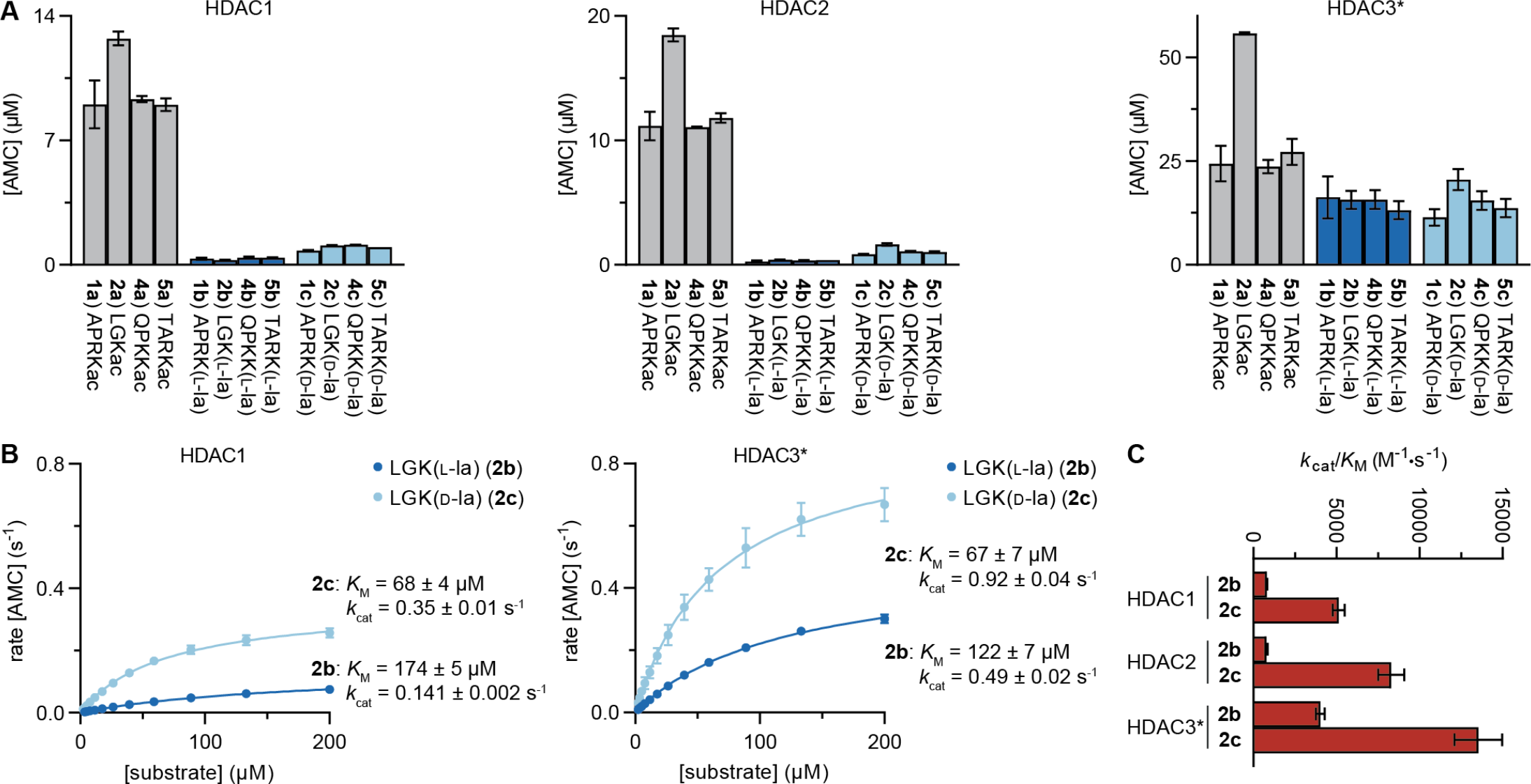
Delactylase activity of class I HDACs 1–3. (**A**) Conversion of Kac, K(l-la) and K(d-la) short AMC-conjugated peptides. Data represent mean ± SD, *n* = 2. (**B**) Michaelis-Menten plots for HDACs 1 and 3 against substrates **2b** and **2c**. Data represent mean ± SEM, *n* = 2 (see fig. S4 for HDAC2 data and sample assay progression curves). (**C**) Catalytic efficiencies of HDACs 1–3 against substrates **2b** and **2c**. Data represent mean ± SEM, *n* = 2 (see fig. S4 for complete *K*_M_ and *k*_cat_ data sets). *HDAC3 incubated with the DAD of NCoR2.

We then determined the delactylase efficiency of each isozyme. Conversion of several K(l-la) and K(d-la) substrates by HDAC3 had exceeded 30% in the previous experiment (initial substrate concentration: 50 μM), which indicates that the steady state was not maintained during the assay. Thus, to gain accurate kinetic insight, conversions of the l- and d-lactylated Ac-LGK-AMC substrates **2b** and **2c** were studied using an enzyme-coupled continuous assay (*45*). This revealed that HDAC1–3 generally remove the K(d-la) modification more efficiently than K(l-la) (Fig. 2B and fig. S4). In agreement with the end-point assay data, HDAC3 was the least sensitive to the configuration of the chiral center in the modification (3.5-fold higher *k*_cat_/*K*_M_ for **2c** than for **2b**, compared to >6-fold for HDACs 1 and 2) and was also the most efficient delactylase enzyme across all HDACs (Fig. 2C).

The delactylase and deacetylase activities of HDACs 1–3 were also studied with longer, non-fluorogenic peptides, in order to account for the potential effect of the fluorophore on enzyme activity and further characterize the site-specific delactylase activity of these enzymes. Six different peptides from histones H2B, H3, and H4 were prepared by solid-phase peptide synthesis (SPPS) and site-selectively modified with Kac, K(l-la) and K(d-la) modifications (Fig. 3A, see Supplementary Methods for synthetic details). The deacetylase and delactylase activities of HDACs 1–3, measured by HPLC after 1 h incubation (Fig. 3B), followed similar patterns for all three modifications and correlated with those observed with fluorogenic substrates. However, larger differences in conversion were obtained depending on the sequence of the peptides. The sequence requirements of HDAC1/2 were different than those of HDAC3: HDAC1/2 showed lower activity at the H3K23 (**8**) and H4K12 (**10**) sites than HDAC3, relative to other peptides, while the H3K9 (**6**), H3K18 (**7**), and H4K8 (**9**) substrates were converted to a larger extent by all three enzymes (Fig. 3C). Interestingly, the N-terminal peptide of H2B with modification at lysine 5 (H2BK5, **5**) was poorly recognized by HDAC1–3. SIRT1–3, which have been shown to remove K(l-la) modifications from a pyruvate kinase M2 (PKM2) peptide to some extent (*46*), and Kbhb modifications from histone H3 peptides (*47*), only exhibited minor activity against the H3K18(l-la) (**7b**) and H3K18(d-la) (**7c**) peptides (Fig. 3D), in agreement with our initial screening data (Fig. 1C). Albeit, SIRT2 and, to a lesser extent, SIRT1 and SIRT3, were able to remove lactyl modifications from purified histones upon 4 h incubation (fig. S1). This could suggest that SIRT2 and SIRT3 serve as lysine delactylases in the cytosol and mitochondria (*46*), respectively, where HDACs 1–3 are not present.

**Fig. 3.**
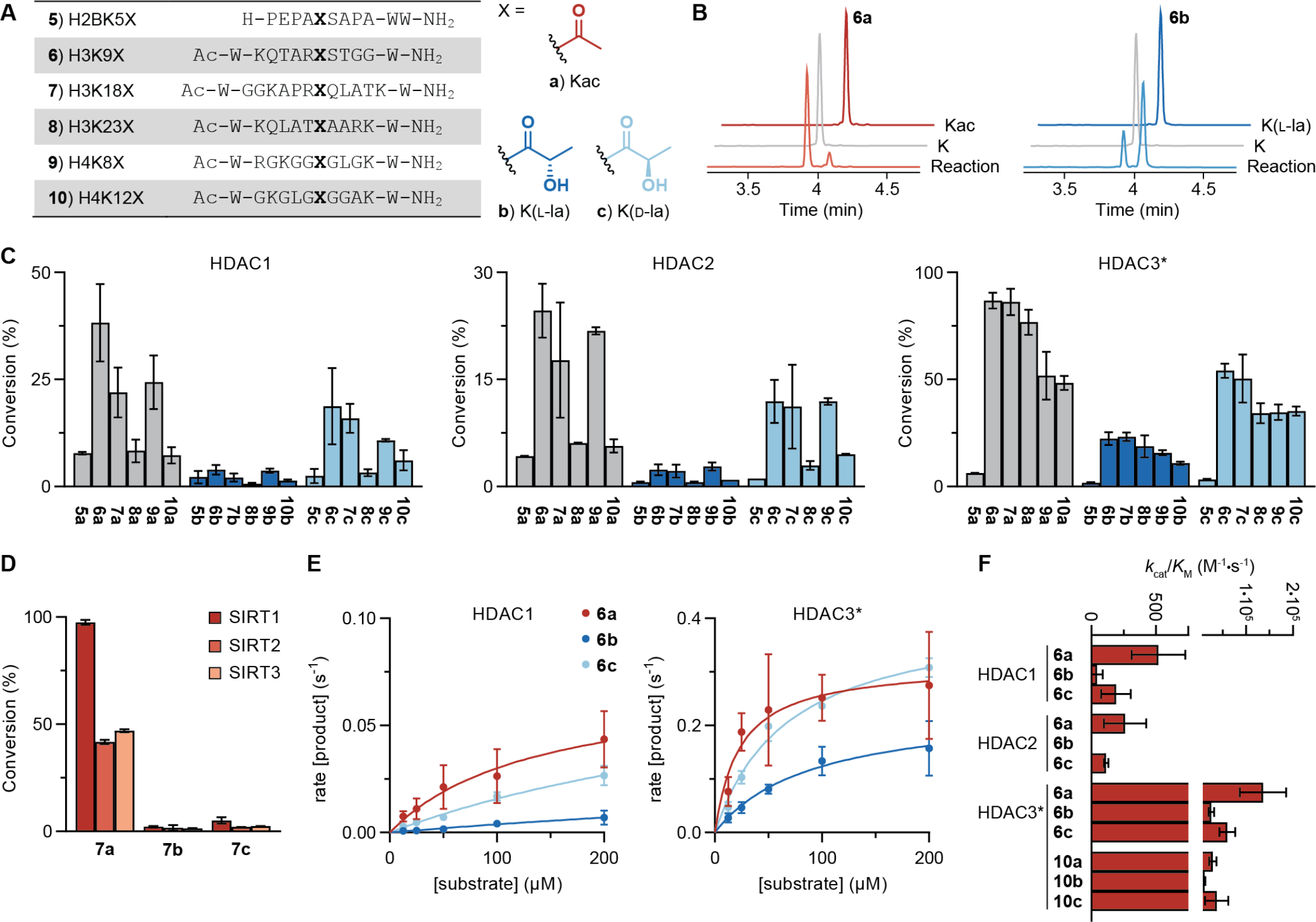
Delactylase activities of HDACs 1–3 at different histone sites. (**A**) Synthesized histone peptide sequences, with X = Kac (**a**), K(l-la) (**b**) or K(d-la) (**c**). (**B**) Sample deacetylation and delactylation HPLC assay traces (60 min reaction of 50 nM HDAC3/NCoR2 with 50 μM peptide **6a** or **6b**). (**C**) Relative conversion of Kac, K(l-la) or K(d-la)-containing histone peptides. Data represent mean ± SD, *n* = 2. (**D**) Relative conversion of Kac, K(l-la) or K(d-la)-containing H3K18 peptide by sirtuins 1–3. Data represent mean ± SD, *n* = 2. (**E**) Michaelis-Menten plots for HDACs 1 and 3 against substrates **6a**, **6b** and **6c**. Data represent mean ± SEM, *n* = 2 (see fig. S5 for HDAC2 data). (**F**) Catalytic efficiencies of HDACs 1–3 against substrates **6a–c** and **10a–c**. Data represent mean ± SEM, *n* = 2 (see fig. S5 for Michaelis-Menten plots of substrates **10a–c** and numerical data). *HDAC3 incubated with the DAD of NCoR2.

Conversion of H3K9 peptides **6a–c** was measured at several substrate concentrations after 10, 15 and 20 min, in order to obtain steady-state conversion rates (Fig. 3E). Michaelis-Menten analysis again revealed a preference for K(d-la) over K(l-la) for HDACs 1 and 2, although slightly less pronounced than that observed with fluorogenic substrates (<5-fold higher *k*_cat_/*K*_M_ for **6c** than for **6b**) (Fig. 3F). In addition, HDAC3 was the most efficient delactylase when compared to its deacetylase activity, both at the H3K9 and H3K18 sites (Fig. 3F), which further supports our observations using short fluorogenic substrates. Furthermore, the efficiencies of HDAC3 for hydrolysis of K(l-la) and K(d-la) on non-fluorogenic peptides were >1,000-fold higher than those reported for SIRT2 (*46*).

### HDAC3 removes various small aliphatic and hydroxylated acyl-lysine PTMs

HDAC3 in association with the NCoR/SMRT nuclear co-repressor complex is a potent lysine deacetylase (*48, 49*), but it also harbors additional catalytic activities. Recent work has revealed that histone ε-*N*-crotonyllysine (Kcr) and ε-*N*-β-hydroxybutyryllysine (Kbhb) modifications are installed by p300 (*37, 50*). HDAC3 hydrolyzes Kcr and Kbhb in vitro and in human cells (*34–37*) and other aliphatic acyl modifications of up to 10 carbons in length can be removed as well (*31*). To elaborate upon this ability and compare its delactylase activity to those reported before, we prepared a collection of H4_10-12_ substrates with small aliphatic and hydroxylated modifications to study the substrate scope of HDAC3 further (Fig. 4A).

**Fig. 4.**
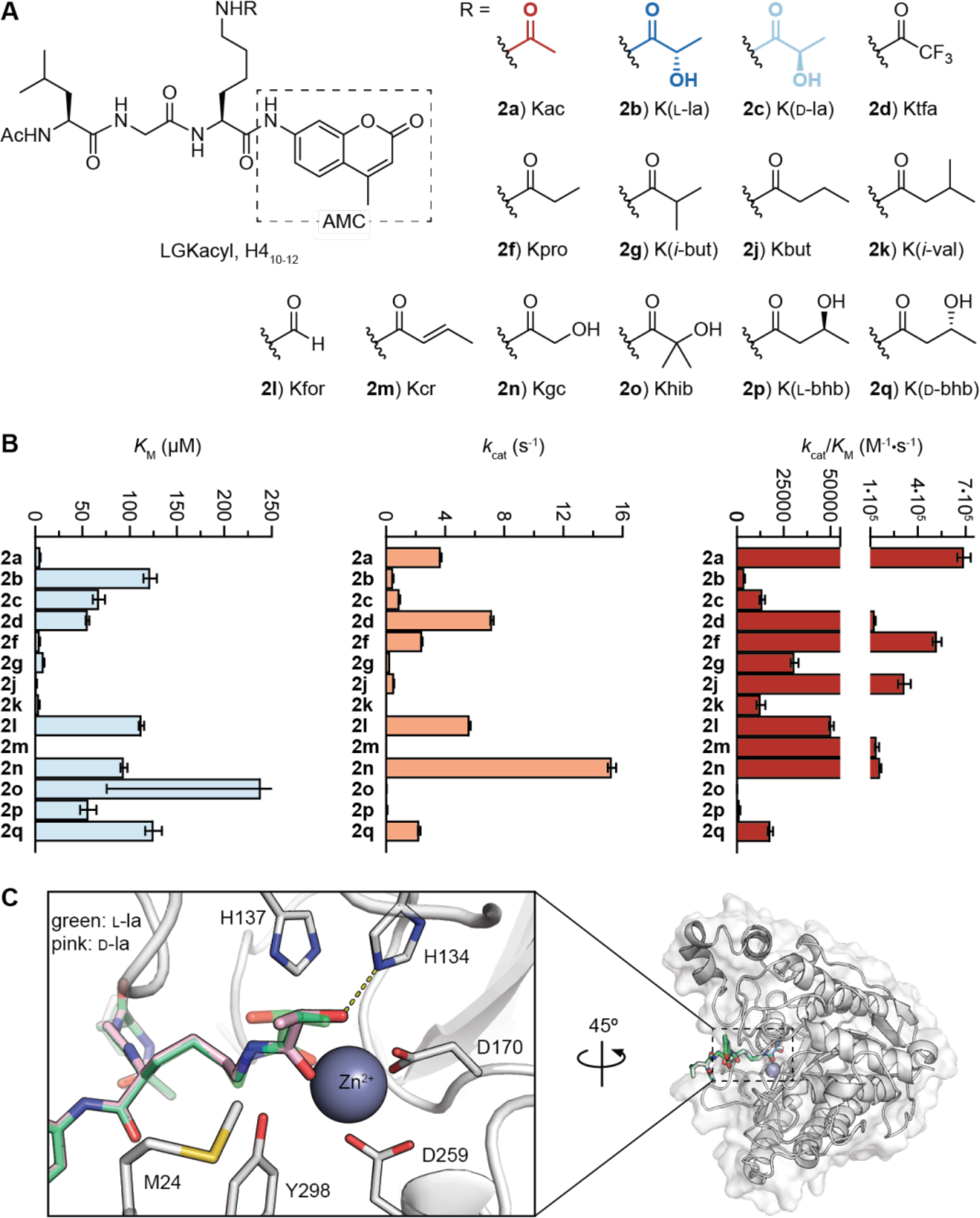
Acyl-lysine substrate scope for HDAC3. (**A**) Structure of H4_10-12_ (Ac-LGK-AMC) substrates. (**B**) Kinetic parameters of HDAC3/NCoR2 against H4_10-12_ substrates with various acyl-lysine modifications. Data represent mean ± SEM, *n* = 2 (see fig. S4 for Michaelis-Menten curves and numerical data). (**C**) Docking of K(l-la) (green) and K(d-la) (pink) substrate mimics into the crystal structure of HDAC3 (pdb 4A69). A hydrogen bond between H134 and the hydroxy group of the K(d-la) modification is predicted.

Michaelis-Menten experiments provided highly diverse steady-state parameters and highlighted critical structural substrate requirements (Fig. 4B). Short linear aliphatic PTMs (Kac, **2a**; Kpro, **2f**; Kbut, **2j**; Kcr, **2m**) exhibited Michaelis-Menten constants (*K*_M_) below 10 μM, in agreement with previous studies for this isozyme (*34, 51*). These four substrates, as well as the ε-*N*-trifluoroacetyllysine modification (Ktfa, **2d**) employed for screening of HDAC inhibitors (*39*), were all cleaved with efficiencies (*k*_cat_/*K*_M_) over 100,000 M^−1^·s^−1^. Addition of a single hydroxy group to Kac to give ε-*N*-glycolyllysine (Kgc, **2n**) dramatically increased the substrate *K*_M_ value but still afforded high efficiency, due to an unusually high catalytic rate constant (*k*_cat_ = 15.3 ± 0.3 s^−1^, Fig. 4B). Branched aliphatic chains [K(*i*-but), **2g**; K(*i*-val), **2k**] provided lower *k*_cat_ values resulting in a ~10-fold lower efficiency (*31*). The efficiency was also lower against branched modifications containing hydroxy groups [K(l-la), **2b**; K(l-bhb), **2p**; Khib, **2o**; etc.], resulting from a combination of lower *k*_cat_ and higher *K*_M_ values. Interestingly, both ε-*N*-d-β-hydroxybutyryllysine [K(d-bhb)] and ε-*N*-2-hydroxyisobutyryllysine (Khib) have been found on histones and connect cell metabolism with gene expression regulation (*52, 53*). Kbhb is also cleaved by SIRT1–3, although with lower efficiency in vitro (*47*). SIRT3 exhibits slight preference for K(l-bhb) over K(d-bhb) (*47*), whereas HDAC3 shows opposite selectivity (Fig. 4B), which could indicate complementary regulatory mechanisms by HDACs and sirtuins (*37*).

Overall, HDAC3 showed efficiencies under 20,000 M^−1^·s^−1^ for all hydroxylated acyl-lysine PTMs except Kgc (Fig. 4B). These data categorized the l- and d-delactylase activities of HDAC3 among the less efficient catalytic conversions for HDACs in vitro (*34, 39*), yet similar to those of HDAC8 or sirtuins recorded for their validated cellular targets in similar assay formats (*54, 55*).

To rationalize the preference of HDAC3 for K(d-la) over K(l-la), small molecule substrate mimics containing both PTMs were docked into the crystal structure of HDAC3 bound to the DAD of NCoR2 (pdb 4A69, Fig. 4C) (*56*). Both modifications were accommodated in the HDAC3 active site in a conformation similar to that of a Kac substrate bound to HDAC8 (*57*). The HDAC8 structure was chosen for comparison because it is the only class I HDAC isozyme co-crystallized with an acylated substrate thus far. Interestingly, the hydroxy group of the K(d-la) modification could be positioned to form a hydrogen bond with histidine 134 (H134) (Fig. 4C, pink molecule), which was not possible with the hydroxy group of K(l-la) as it pointed in the opposite direction (Fig. 4C, green molecule). The H134 residue is conserved in Zn^2+^-dependent HDACs and involved in transition state stabilization (*58*). Thus, this stereoisomer-specific interaction could explain the higher hydrolytic efficiency of HDAC1–3 for K(d-la) over K(l-la).

### Quantification of HDAC3-mediated histone delactylation in vitro

To corroborate the results in peptide-based in vitro assays, we examined the delactylase activity of HDAC3 on histones by incubation of recombinant HDAC3 with acid-extracted histones from HeLa cells (*59*). We found that global histone lactylation levels were decreased by HDAC3, an activity that could be inhibited by both the broad spectrum HDAC inhibitor TSA and the HDAC1–3-specific inhibitor apicidin (Fig. 5A) (*39, 60*). We considered the use of RGFP966 as HDAC3-selective inhibitor as well (*61*). However, in our hands, this compound behaved as a slow-binding inhibitor of HDACs 1–3 (*62*), and thus not more selective than apicidin (fig. S6). To gain further insights into which lactylation sites are regulated by HDAC3, we used a stable isotope labeling by amino acids in cell culture (SILAC)-based quantification method (*63*). Equal number of histones extracted from “light” or “heavy”-labeled cells were incubated with or without recombinant HDAC3 for 4 h. Then, after tryptic digestion, the “light” and “heavy” histone peptides were mixed at a ratio of 1:1 (m/m) and analyzed by HPLC-mass spectrometry (MS). To eliminate potential bias on PTM levels caused by SILAC labeling, we performed both forward and reverse labeling, and in both replicates histone acetylation and lactylation were efficiently removed by HDAC3 (Fig. 5B–D). We detected 39 K(l-la) peptides on histones, among which 31 K(l-la) peptides were decreased by more than 90% in response to HDAC3 in at least one replicate and 38 K(l-la) peptides showed >50% decrease in both replicates (Fig. 5B and table S1). Remarkably, lactylated N-terminal H2B peptides were decreased to a lesser extent than most H3 and H4 peptides (table S1), which was in agreement with the lower conversion of H2BK5 substrates (Fig. 3C). Overall, our results demonstrate that HDAC3 has robust histone delactylase activity in vitro.

**Fig. 5.**
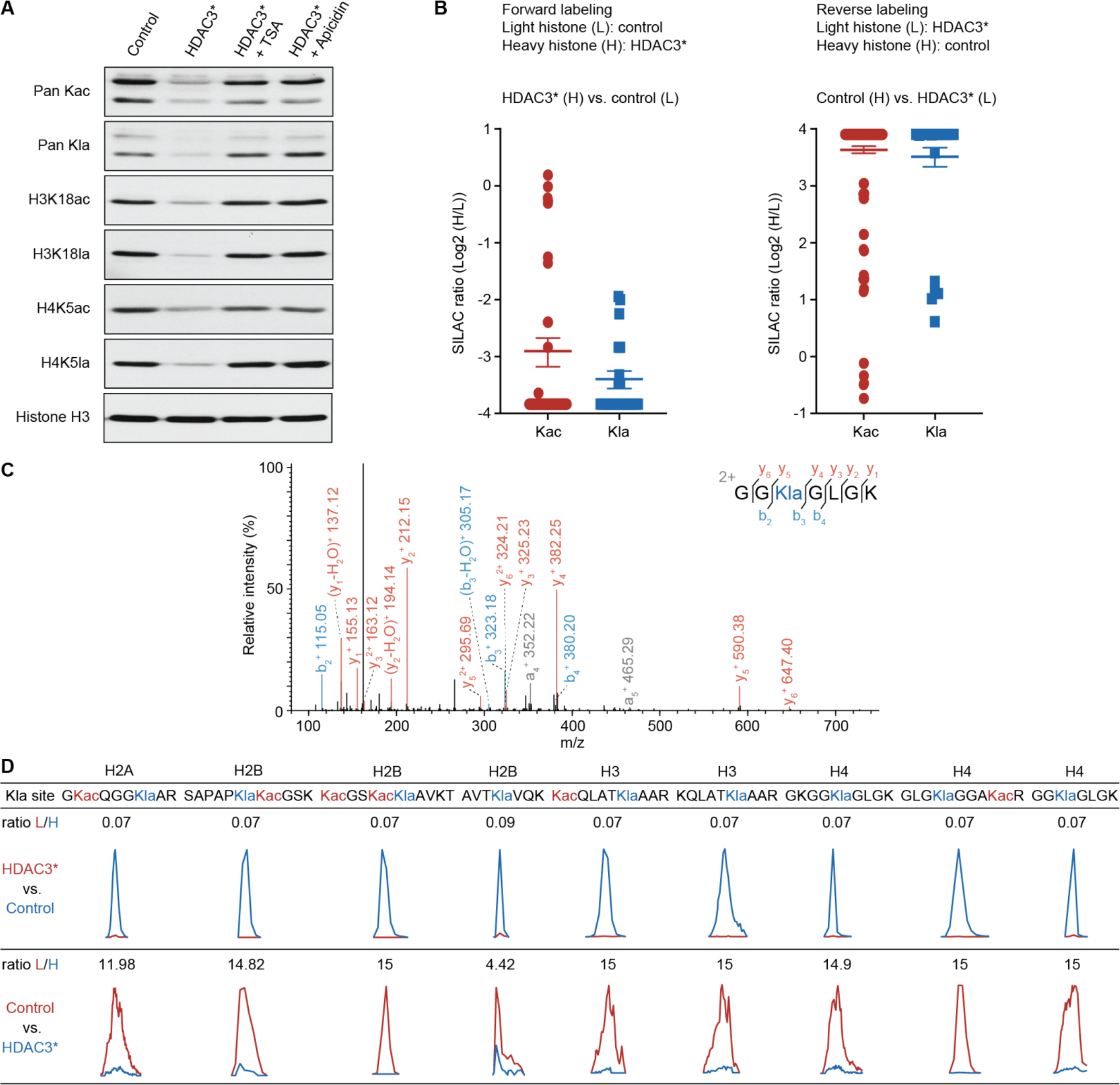
Mass-spectrometric identification and quantification of in vitro HDAC3-catalyzed delactylation targets on core histones. (**A**) Changes in lactylation and acetylation of histones (overall, and at the sites H3K18 and H4K5) upon HDAC3 treatment, and inhibition with TSA and apicidin. (**B**) Forward and backward MS quantification of Kla and Kac sites, with or without HDAC3. (**C**) A representative MS/MS spectrum of the histone H4 peptide GGKlaGLGK. (**D**) Representative MS quantification from in vitro histone deacylation products shown in (**B**). A full list of quantified peptides can be found in table S1.

### HDAC1 and HDAC3 remove ε-*N*-lactyllysine modifications on histones in the cell

We then explored whether the delactylase activity of HDACs 1–3 is maintained in living cells in culture. Treatment of HeLa cells for 5 h with broad spectrum HDAC inhibitors (sodium butyrate and TSA) increased global histone lactylation levels and the HDAC1–3-specific inhibitor apicidin also enhanced lactylation (Fig. 6A and fig. S7). In contrast, the class IIa HDAC inhibitor TMP195 and the sirtuin inhibitor nicotinamide (NAM) failed to affect histone lactylation levels substantially (Fig. 6A). This result indicated that HDACs 1–3 are the major delactylases in cells, which is consistent with our in vitro data.

**Fig. 6.**
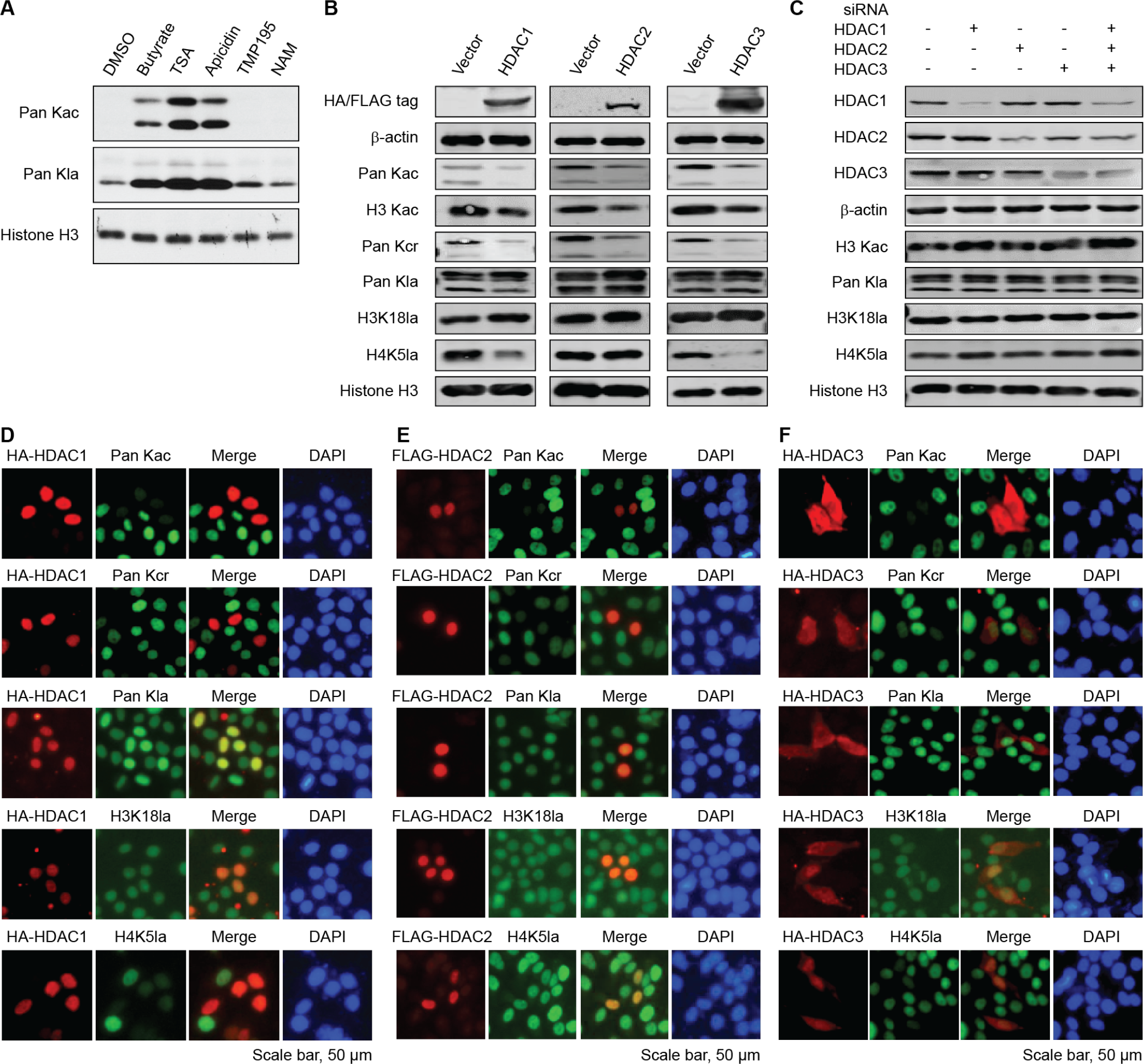
In cellulo deacetylase, decrotonylase and delactylase activities of HDACs 1–3. (**A**) Changes in overall lactylation and acetylation of histones in HeLa cells upon 5 h treatment with pan-HDAC inhibitors (sodium butyrate, 2 mM; TSA, 1 μM), the HDAC1–3-selective inhibitor apicidin (1 μM), the class IIa-selective inhibitor TMP195 (5 μM), or the pan-sirtuin inhibitor nicotinamide (NAM, 10 mM). See fig. S7 for full Western blots. (**B**) Western blot analysis of core histones from HeLa cells with or without transfection of HA-tagged HDAC1, FLAG-tagged HDAC2 or HA-tagged HDAC3. (**C**) Western blot analysis of core histones from HeLa cells with or without siRNA-mediated knockdown of HDAC1, 2, 3, or their combination (72 h treatment). Top blots correspond to whole-cell lysates and show the level of ectopically expressed HDAC (**B**) or HDAC knockdown (**C**) relative to β-actin. See fig. S7 for full Western blots. (**D–F**) Immunostaining images of HeLa cells transfected with HA-tagged HDAC1, FLAG-tagged HDAC2 or HA-tagged HDAC3 and the levels of histone acetylation, crotonylation, and lactylation. DAPI: 4’,6-diamidino-2-phenylindole (nuclear stain).

To further dissect the action of each isozyme, HDACs 1–3 were overexpressed in HeLa cells, as verified by Western blot (Fig. 6B). Overexpression of each isozyme resulted in a reduction in the overall levels of histone acetylation (pan Kac) and crotonylation (pan Kcr) (*35, 64*), but no change was observed for overall levels of histone lactylation (pan Kla) (Fig. 6B). Site-specific lactylation levels at H4K5, on the other hand, were reduced by HDACs 1 and 3, while the H3K18la modification remained unchanged in all three experiments (Fig. 6B). These results were corroborated by immunofluorescent staining and analysis of individual nuclei (Fig. 6D–F). HDAC1 and 2 constructs were expressed in the nuclei, while HDAC3 was also observed in the cytosol, as reported (*65, 66*). In these experiments, cells with detectable levels of transfected HDACs 1 and 3 also showed strong reduction of Kac, Kcr, and H4K5la levels, but not for the H3K18la mark or global Kla modification. Thus, overexpression of a single class I HDAC is not sufficient for altering the overall levels of histone lactylation to an extent detectable by Western blot or immunofluorescence. Moreover, HDACs 1–3 exhibit site-selective activities in the cell, which are different from those observed in vitro (Fig. 3C). A plausible explanation for this effect would be the formation of nuclear multiprotein complexes able to direct the delactylase activity towards specific histone sites (*67*).

To validate our findings from the overexpression experiments, we performed RNA interference experiments to knock down HDACs 1–3. Western blot analysis revealed that knockdown of HDAC1 or HDAC3 but not HDAC2 slightly increased histone H4K5 lactylation levels, and triple knockdown of HDACs 1–3 led to a stronger effect on H4K5 lactylation (Fig. 6C). In sum, our cell-based assays suggest that HDAC1 and HDAC3 have delactylase activity in cells.

## Discussion

Metabolic intermediates are chemical sources for histone PTMs and affect gene expression (*1, 16, 68*). In particular, the growing landscape of histone acyl modifications has revealed a dynamic and diverse network of metabolic modifications that regulate transcription in health and disease (*14, 52, 53*). The metabolic importance of l-lactate and its association with disease have been known for almost a century, but it was only recently discovered as a precursor for stimulating histone K(l-la) marks that in turn modulate gene expression (*14*). K(l-la) modifications, similar to other histone acylations, can be installed by HATs (*14, 37, 50*), but little is known about their reversibility and turnover. Identification of key regulatory enzymes, lactyltransferases and delactylases, as well as their substrate proteins, will be a stepping stone for dissecting the functions of l-lactate signaling pathways in the future.

In this paper, we report the first systematic in vitro screening of delactylase activity using synthetic peptides and core histones as substrates. The nuclear HDACs 1–3 were the most robust and efficient delactylases, while the only activity detected for the sirtuins in vitro was on purified histones. These results were corroborated in cells by the effect of HDAC and sirtuin inhibitors with different specificities. Nevertheless, it remains to be addressed in the future whether sirtuins 1–3 can act as delactylases at protein sites complementary to HDACs or in other cellular compartments, as it has been proposed for SIRT2 in the cytosol (*46*). By overexpression and knockdown experiments, we revealed that HDACs 1 and 3 are efficient delactylases in the cell, but we also observed that individual isozymes cannot alter the overall levels but H4K5 of histone lactylation to a large extent. These data suggest the existence of site-specific K(l-la) regulatory pathways, and potentially other unknown delactylases exist in cells.

HDACs 1–3 perform their catalytic functions as part of large multicomponent protein complexes in the nucleus (*67*). These complexes consist of scaffolding proteins that allow the HDACs to interact with other epigenetic enzymes, such as demethylases (*69*) and transcription factors (*67*). The activity of the HDACs is enhanced when engaged in their cognate complexes (*48*), and standard recombinant preparations of HDAC3 include the directly-interacting DAD domain of NCoR2 for increased activity. This might explain the enhanced delactylase activity observed in vitro for HDAC3 compared to HDACs 1 and 2. It has also been shown that HDACs 1–3 display distinct Kac substrate selectivity upon interaction with different complex partners (*70*), which could be the reason for the discrepancies observed between in vitro and cellular data. Interestingly, HDACs 1 and 3, but not HDAC2, can prominently change the levels of lactylation at the H4K5 site. HDAC1 and HDAC2 are frequently present in the same multiprotein complex in the nucleus and are thought to compensate for each other in the regulation of histone Kac (*67, 71*). Our data show that HDAC1 and HDAC2 may have unique individual deacylation substrates and thus indicate distinct functions of these two enzymes in cells. However, it remains to be studied to what extent the formation of complexes directs HDAC activity toward specific histone K(l-la) sites.

*n-*Butyrate and other short-chain fatty acids modify gene transcription patterns via inhibition of HDACs leading to histone hyperacetylation (*72, 73*). This epigenetic mechanism has also been attributed to Land d-lactate (*74*), and to β-hydroxybutyrate (*75*). However, the concentrations necessary for achieving HDAC inhibition by these hydroxylated species are in the high millimolar range (~100-fold higher than that of *n*-butyrate) and often lead to minor changes in acetylation (*74, 76, 77*). Alternatively, dynamic covalent modification of histones by l-lactyl-CoA and other activated acyl groups may explain their regulatory effects at physiological concentration.

l-Lactate governs lactate metabolism and signaling in humans (*8*), whose concentration is typically 100 times higher than d-lactate in cells. Both l-lactate and d-lactate are known precursors for covalent protein modifications (*14, 22*). l-Lactate labels proteins through enzyme-catalyzed reaction (likely via lactyl-CoA) (*14, 15*), which was demonstrated in nuclei but can most likely happen in the cytosol similar to other lysine acylation events, e.g. lysine acetylation (*78*). HDACs 1–3 show remarkable activities toward diverse ε-*N*-acyllysine groups including K(d-la), and it remains to be confirmed if these activities are physiologically relevant. Here, we demonstrate that HDACs 1–3 are responsible for the reversible and dynamic regulation of histone l-lactylation. This result and the specificity of genomic localization of histone l-lactylation observed by ChIP-seq (*14*) argue loudly that histone l-lactylation is likely the major pathway in nuclear lactylation. Taken together, these findings not only provide insight into the new activities of class I HDACs but also validate the regulation of histone K(l-la) by pharmacologically targetable enzymes, which lays the foundation for future functional studies of this modification and its associated regulatory pathways.

## Materials and Methods

### Chemical synthesis

Please find the synthesis and characterization of peptide substrates in the Supplementary Methods.

### Assay materials

Fluorescence-based deacylase activity assays were performed in black low binding 96-well microtiter plates (Corning half-area wells, Fischer Scientific, cat. # 3686), with duplicate series in each assay and each assay performed at least twice. Control wells without enzyme were included in each plate. HPLC-based deacylase activity assays were performed in microcentrifuge plastic tubes (1.5 mL, Sarstedt AG, cat. #: 72.690.001) and filtered through 0.45 μm PTFE filters prior to HPLC injection (Macherey-Nagel GmbH & Co., cat. #: 729015), with each assay performed at least twice. Experiments were performed in HEPES buffer [50 mM HEPES/Na, 100 mM KCl, 0.001% (v/v) tween-20, 0.2 mM tris(2-carboxyethyl)phosphine (TCEP), pH 7.4] (*39*) or Tris buffer [50 mM Tris/Cl, 137 mM NaCl, 2.7 mM KCl, 1 mM MgCl, pH 8] (Fluor de Lys buffer, Enzo Life Sciences) with bovine serum albumin (BSA, Sigma-Aldrich, cat. #: A7030) as indicated for each assay. β-nicotinamide adenine dinucleotide hydrate (NAD^+^) for sirtuin assays was from commercial source (Sigma-Aldrich, cat. #: N7004), as well as trypsin (Sigma-Aldrich, cat. #: T1426). The following recombinant enzymes were acquired from BPS Bioscience (San Diego, CA): HDAC1 (full length, C-terminal His tag, cat. #: 50051), HDAC2 (full length, C-terminal His tag, cat. #: 50002), HDAC3/NCoR2 (full length, C-terminal His tag, with DAD domain of NCor2, N-terminal GST tag, cat. #: 50003), HDAC4 (aa 627–1084, N-terminal GST tag, cat. #: 50004), HDAC5 (aa 656–1122, C-terminal His tag, cat. #: 50005), HDAC6 (full length, C-terminal FLAG tag, cat. #: 50056), HDAC7 (used only with extracted histones, aa 518-end, N-terminal GST tag, cat. #: 50007), HDAC8 (full length, C-terminal His tag, cat. #: 50008), HDAC9 (aa 604–1066, C-terminal His tag, cat. #: 50009), HDAC10 (used only with extracted histones, aa 2-631, N-terminal FLAG tag, cat. #: 50060), HDAC11 (full length, untagged, cat. #: 50021), SIRT1 (aa 193–741, N-terminal GST tag, cat. #: 50012), SIRT2 (aa 50–356, C-terminal His tag, cat. #: 50013), SIRT3 (aa 102–399, N-terminal GST tag, cat. #: 50014), SIRT4 (aa 25–314, N-terminal GST tag, cat. #: 50015), SIRT5 (used only with extracted histones, full length, N-terminal GST tag, cat. #: 50016), SIRT6 (full length, N-terminal GST tag, cat. #: 50017), SIRT7 (used only with extracted histones, full length, C-terminal FLAG tag, cat. #: 50018); or from Millipore (Temecula, CA): HDAC7 (used only with fluorogenic substrates, aa 383–end, N-terminal His tag, cat. #: 14-832). Recombinant SIRT5 and SIRT7 used with fluorogenic substrates were produced and purified as described below. PTM antibodies anti-Kla (cat. #: PTM-1401), anti-Kac (cat. #: PTM-101), anti-Kcr (cat. #: PTM-501), and sequence-specific anti-H3K18la (cat. #: PTM-1406) and anti-H4K5la (cat. #: PTM-1407) were purchased from PTM Biolabs, Inc. (Chicago, IL). Anti-histone H3 (cat. #: 4499) and anti-actin (cat. #: 4970) were purchased from Cell Signaling Technology, Inc. (Danvers, MA), anti-FLAG (cat. #: F7425/F1804) was purchased from MilliporeSigma (Burlington, MA); anti-HA (cat. #: SC-805) was purchased from Santa Cruz Biotechnology, Inc. (Dallas, TX); and anti-HDAC1 (cat. #: A0238), anti-HDAC2 (cat. #: A2084) and anti-HDAC3 (cat. #: A2139) were purchased from ABclonal, Inc. (Woburn, MA). Inhibitors were of commercial source: apicidin (Sigma-Aldrich, cat. #: A8851), entinostat (MS-275, TargetMol, cat. #: T6233), nicotinamide (MilliporeSigma, cat. #: N0636), sodium butyrate (MilliporeSigma, cat. #: B5887), TMP195 (Cayman Chemical Company, cat. #: 23242), trichostatin A (TSA, Tokyo Chemical Industry, cat. #: T2477; or MilliporeSigma, cat. #: T8552), RGFP-966 (Sigma-Aldrich, cat. #: 16917). Stocks were prepared in DMSO (1–40 mM), and concentration of peptide substrates were determined based on absorbance [ε280(Trp) = 5690 M^−1^·cm^−1^ or ε326(Ac-Lys-AMC) = 17783 M^−1^·cm^−1^] (*79*) using a Thermo Scientific NanoDropC instrument. Assay concentration of substrates and inhibitors were obtained by dilution from DMSO stock solutions in buffer, and appropriate concentration of enzyme obtained by dilution of the stock provided by the supplier. Data analysis was performed using GraphPad Prism 8 or 9.

### Expression and purification of SIRT5 and SIRT7

pET100/D-TOPO vectors encoding *E. coli* codon-optimized SIRT5 (aa 34-310) or SIRT7 (aa 1-400) and an N-terminal cleavage site for Tobacco Etch Virus (TEV) protease were designed in GeneArt™ and purchased from ThermoFischer Scientific. Proteins were expressed in *E. Coli* BL21(DE3), and cells were harvested and disrupted by sonication for 6×1 min in lysis buffer [SIRT5: 12.5 mM Na2HPO4, 1.8 mM KH_2_PO_4_, 2.7 mM KCl, 137 mM NaCl, 1% Tween-20, 15 mM β-mercaptoethanol, pH 7.6; SIRT7: 50 mM Tris, 400 mM NaCl, 20 mM β-mercaptoethanol, 10% glycerol, pH 8]. The supernatants were filtrated and subjected to affinity chromatography using a 5 mL HisTrap HP His-tag purification column (GE healthcare, cat. # GE29-0510-21) operated by a peristaltic pump (Cytiva, cat. # 18111091) at 4 °C. Supernatants were run onto the column followed by 25 mL lysis buffer. SIRT5 was eluted with a gradient of 0-50% elution buffer (SIRT5 lysis buffer with 500 mM imidazole, pH 7.6), and SIRT7 was eluted with 25 mL elution buffer (SIRT7 lysis buffer with 500 mM imidazole, pH 8). Fractions were collected using an ÄKTApurifier system with ÄKTA Frac-950 fractions collector (GE Healthcare Life Sciences). The eluted fractions were analyzed by gel electrophoresis and Coomasie blue-staining, collected, dialyzed against lysis buffer using Spectra/Por 4 dialysis tubing (12-14 kDa MWCO, Spectrum, cat. # 132697), and treated with TEV protease (0.2 mg/mL) overnight at 4 °C. Samples were then subjected to reverse affinity chromatography using a 5 mL HisTrap HP His-tag purification column at 4 °C. Flow through and following washes were collected and concentrated to <1 mL using an Ultra-15 Centrifugal Filter unit with cut-off value 30 kDa (Amicon, #UFC903024). The purified SIRT7 was then stored at −80 °C. The SIRT5 sample was subjected to gel filtration on a Superdex™ 75 HR 10/30 column (Amersham) in buffer (25 mM Tris/HCl, 100 mM NaCl, 15 mM β-mercaptoethanol E, 10% v/v glycerol, pH 7.5) using the ÄKTApurifier system, and fractions containing SIRT5 were pooled, concentrated and stored at −80 °C. These enzymes were used only with fluorogenic substrates.

### Fluorogenic lysate deacylase assay

HEK293T cells (ATCC) were cultured under a humidified 5% CO_2_ atmosphere in Dulbecco’s modified Eagle’s medium (DMEM, Thermo Scientific, cat. #: 11965118) supplemented with 10% (v/v) fetal bovine serum (FBS), 1% penicillin and 1% streptomycin. For lysate preparation, media was aspirated, cells were washed with PBS and treated with lysis buffer (1% Triton X-100, 0.2% SDS and cOmplete EDTA-free protease inhibitor cocktail, COEDTAF-RO Sigma-Aldrich, in PBS). Upon scraping, lysates were collected in microcentrifuge tubes, sonicated with a Bandelin Sonopuls mini20 (2 s on, 2 s off, 80% amplitude, 1 min), centrifuged (14,000 g, 10 min, 4 °C), and protein concentrations of the supernatants were determined by the bicinchoninic acid assay (BCA1, Sigma-Aldrich). Lysates (30 μg per well) were then incubated with substrates **1a–c** (50 μM), and NAD^+^ (500 μM) and TSA (100 μM), or NAM (10 mM) in Tris buffer with 0.5 mg/mL BSA at a final volume of 25 μL per well. The reaction was incubated at 37 °C for 60 min, then a solution of trypsin (25 μL, 5.0 mg/mL; final concentration of 2.5 mg/mL) and NAM (4 mM; final concentration of 2 mM) was added, and the assay development was allowed to proceed for 90 min at room temperature before fluorescence analysis. The data were analyzed to afford [AMC] relative to control wells without enzyme.

### Fluorogenic substrate screening assays

The initial screening for substrate deacylation activity (Fig. 1D) was performed in Tris buffer with 0.5 mg/mL BSA, with end-point AMC fluorophore release by trypsin. For a final volume of 25 μL per well, acyl substrates (50 μM) and NAD^+^ (500 μM, only for sirtuin assays) were added to each well, followed by a solution of the appropriate KDAC enzyme (HDACs: 50 nM, SIRT1, 2, 5: 200 nM, SIRT3, 4, 6, 7: 500 nM). The reaction was incubated at 37 °C for 60 min (HDACs) or 120 min (sirtuins), then a solution of trypsin (25 μL, 5.0 mg/mL; final concentration of 2.5 mg/mL) was added, and the assay development was allowed to proceed for 90 min at room temperature before fluorescence analysis. The data were analyzed to afford [AMC] relative to control wells without enzyme.

Follow-up screening with Kac, K(l-la) and K(d-la) fluorogenic substrates (Fig. 2A) was performed in HEPES buffer with 0.5 mg/mL BSA, with end-point AMC fluorophore release by trypsin. Substrate (50 μM) and enzyme (2 nM and 10 nM) were incubated at 37 °C for 60 min in a final volume of 25 μL. Then, a solution of trypsin (25 μL, 5.0 mg/mL; final concentration of 2.5 mg/mL) was added, and the assay development was allowed to proceed for 90 min at room temperature before fluorescence analysis. The data were analyzed to afford [AMC] relative to control wells without enzyme.

### Fluorogenic Michaelis-Menten assays

Rate experiments for determination of kinetic parameters with AMC-containing substrates were performed in HEPES buffer with 0.5 mg/mL BSA. Trypsin concentration was optimized for HDACs 1–3 at 100 μM substrate concentration, in order to ensure trypsin-mediated AMC release was not the rate-limiting step while minimizing its effect on HDAC stability (*45*). For a final volume of 50 μL, substrate (**2a**, **2f**: 100–1.74 μM; **2b**, **2c**: 200–2.31 μM; **2d**, **2g**, **2j**, **2k**, **2l**, **2n**, **2o**, **2p**, **2q**: 200–3.47 μM; **2m**: 25–43 μM; 1.5-fold dilutions), trypsin (10 μg/mL) and enzyme (HDAC1: 30 nM; HDAC2: 15 nM; HDAC3 with **2a**, **2d**, **2f**, **2n**: 1 nM, with **2b**, **2c**, **2l**: 5 nM, with **2g**, **2j**, **2k**, **2q**: 10 nM, and with **2m**, **2o**, **2p**: 20 nM) were added to a microtiter plate and immediately placed in the plate reader. *In situ* fluorophore release was monitored by fluorescence readings every 30 s for 60 min at 25 °C. Then, initial conversion rates (v_0_) were determined for each concentration and data were fitted to the Michaelis-Menten equation to afford *K*_M_ and *k*_cat_ values (*45*).

### HPLC-based substrate assays

Substrate assays with non-fluorogenic peptide substrates were performed in HEPES buffer (HDACs) or Tris buffer (sirtuins) without BSA, with end-point chromatographic determination of relative conversion. Substrate (50 μM) and enzyme (HDAC1, HDAC2: 250 nM; HDAC3: 50 nM; SIRT1: 50 nM; SIRT2, SIRT3: 100 nM) were incubated at 37 °C for 60 min in a final volume of 30 or 40 μL in microcentrifuge tubes. Then, MeOH/HCOOH [94:6 (v/v)] was added to quench the reaction (15 or 20 μL, for 45 or 60 μL total volume), and samples were agitated on an orbital shaker and centrifuged at 20,000 g for 10 min. Samples were then taken up with a plastic syringe, filtered, and analyzed by analytical HPLC on an Agilent 1260 Infinity II series system equipped with a diode array detector. A gradient with eluent III (H_2_O/MeCN/TFA, 95:5:0.1, v:v) and eluent IV (0.1% TFA in MeCN) rising linearly 0–40% during *t* = 1–5 min was applied at a flow rate of 1.2 mL/min in an HPLC column over at 40 °C. The obtained chromatograms at 280 nm were used to obtain relative substrate conversion data and, additionally, control substrate and product samples were analyzed by UPLC-MS to obtain ionization spectra.

### HPLC-based Michaelis-Menten assays

Michaelis-Menten parameters for long peptide substrates were obtained by measuring conversion rates in a discontinuous manner. Assays were performed in HEPES buffer without BSA, where substrate (200–12.5 μM, 2-fold dilutions) and enzyme (HDAC1 with **6a**: 150 nM, with **6b**, **6c**: 250 nM; HDAC2 with **6a**: 150 nM, with **6b**, **6c**: 250 nM; or HDAC3 with **6a**: 10 nM, with **6b**, **6c**, **10a**, **10c**: 50 nM, with **10b**: 150 nM) were incubated in triplicate at 37 °C in a final volume of 30 μL in microcentrifuge tubes. Then, MeOH/HCOOH [94:6 (v/v)] was added to quench each of the reactions at 10, 15 and 20 min incubation times, respectively (15 μL, for 45 μL total volume). Samples were then agitated on an orbital shaker, centrifuged at 20,000 g for 10 min, taken up with a plastic syringe, filtered, and analyzed by analytical HPLC as before. The obtained chromatograms at 280 nm were used to obtain the relative substrate conversion at each time point. Then, if all three values followed a linear trend, the obtained rates (v_0_) at each substrate concentration were adjusted to the Michaelis-Menten equation to afford *K*_M_ and *k*_cat_ values. The entire experiment was performed at least twice.

### Histone extraction

Histones were extracted using previously published method (*59*). Briefly, HeLa cells were harvested and washed twice with phosphate buffered saline (PBS, 137 mM NaCl, 2.7 mM KCl, 10 mM Na_2_HPO_4_ and 2 mM KH_2_PO_4_) and then suspended in cold extraction buffer (10 mM HEPES pH 8.0, 10 mM KCl, 1.5 mM MgCl_2_, 0.34 M sucrose, and 0.1% Triton X-100) at 4 °C for 30 min to remove cytosolic components. After centrifugation at 1,300 g for 10 min at 4 °C, the pellets were re-suspended in nosalt buffer (3 mM EDTA and 0.2 mM EGTA) and incubated at 4 °C for 30 min to release soluble nuclear components. The pellets were collected after centrifugation at 6,500 g for 10 min at 4 °C and then extracted with 0.2 M H_2_SO_4_ at 4 °C overnight. The supernatants containing histones were collected after centrifugation at 16,000 g for 15 min at 4 °C. The resulting histones were further purified by precipitation using 20% (v/v) trichloroacetic acid, washing with cold acetone, and then dried and stored until further analysis.

### In vitro de-lactylation/acetylation assay using histone as substrates

Histones extracted from HeLa cells (20 μg) were incubated with individual recombinant human HDACs 1–11 or Sirtuins 1–7 (0.2 μg) in reaction buffer for 1 h or 4 h at 37 °C. HDACs 1–11 were tested using the standard Tris buffer, and SIRT1–7 were tested in 50 mM Tris HCl, pH 8.0 with 1 mM Dithiothreitol (DTT), and 1 mM NAD^+^. The reaction was ended by adding sample loading buffer and boiling for 5 min before Western blot analysis.

### Peptide Enrichment

Histones extracted from HeLa cells were dissolved in 50 mM NH4HCO3 buffer and digested with trypsin (Promega) as per the manufacturer’s instructions. The digested histone peptides were incubated with pan anti-Kla beads at 4 °C overnight. The beads were then washed twice with NETN buffer (50 mM Tris HCl, pH 8.0, 100 mM NaCl, 1 mM EDTA, and 0.5% NP-40), once with ETN buffer (50 mM Tris HCl, pH 8.0, 100 mM NaCl, and 1 mM EDTA) and twice with ddH_2_O. The peptides were eluted with 0.1% (v/v) trifluoroacetic acid (TFA) and dried in a SpeedVac system (Thermo Fisher Scientific).

### HPLC/MS/MS analysis

The resulting peptide samples were dissolved in buffer A (0.1% formic acid in water, v/v) and loaded onto a home-made capillary column (10 cm length with 75 μm ID) packed with Jupiter C12 resin (4 μm particle size, 90 Å pore size, Phenomenex) connected to an EASY-nLC 1000 HPLC system (Thermo Fisher Scientific). Peptides were separated and eluted with a gradient of 2% to 90% buffer B (0.1% formic acid in 90% acetonitrile, v/v) in buffer A (0.1% formic acid in water, v/v) at a flow rate of 200 nL min^−1^ over 60 min. The eluted peptides were then ionized and analyzed by a Q-Exactive mass spectrometer (Thermo Fisher Scientific). Full mass spectrometry was acquired in the Orbitrap mass analyzer over the range m/z 300 to 1,400 with a resolution of 70,000 at m/z 200. The 12 most intense ions with charge ≥2 were fragmented with normalized collision energy of 27 and tandem mass spectra were acquired with a mass resolution of 17,500 at m/z 200. The raw data were uploaded into MaxQuant and searched against the UniProt human proteome database.

### Cell culture and transfection

HeLa (ATCC CCL-2) cells were cultured in DMEM high glucose medium (Gibco) containing 10% fetal bovine serum (GeminiBio), 1% GlutaMAX (Gibco) and 5% CO_2_. DNA and siRNA transient transfection were performed using lipofectamine 2000 (Invitrogen) according to the manufacturer’s instructions. siRNAs targeting human HDAC1, HDAC2 and HDAC3 were purchased from GenePharma (Shanghai, China) as a SMART pool.

### Western blot analysis

Protein samples (20 μg whole cell lysate or 2 μg histone) were separated on SDS-PAGE gels (10% for non-histones and 15% for histones) and then transferred to polyvinylidene fluoride (PVDF) membranes. The membranes were blocked in 3% BSA diluted in TBST (20 mM Tris HCl, pH 7.6, 150 mM NaCl, 0.1% Tween-20) for 1 h at room temperature. The membranes were then incubated with primary antibody overnight at 4 °C. The blots were washed with TBST and then incubated for 1 h with rocking in Li-Cor secondary antibodies used at a concentration of 1:10,000. Western blots were developed using a Li-Cor Odyssey.

### Immunofluorescent staining

HeLa cells grown on coverslips were washed with PBS prior to fixation with 4% paraformaldehyde in PBS for 30 min at room temperature. After rinse with PBS twice, the coverslips were incubated with 1% Triton X-100 for 15 min on ice, blocked with 5% BSA for 60 min at 37 °C incubator and incubated with primary antibodies for 2 h at room temperature. The coverslips were washed three times with PBST, followed by incubation with Texas green-conjugated secondary antibodies. Images were acquired with an Olympus microscope system.

### Molecular docking

Molecular modeling was performed with modules from the Schrödinger Small Molecule Drug Discovery Suite (Maestro), release 2018–3, using the OPLS3 force field for parameterization (*80*). The X-ray co-crystal structure of the HDAC3/SMRT-DAD complex (pdb 4A69) was imported from the protein data bank and prepared using the Protein Preparation Wizard using default settings (*81*). Ligands (**2a**, **2b** and **2c**) were prepared using LigPrep (*81*). A receptor-grid box was built in a 25 Å radius around His172 in the catalytic lysine binding pocket and docking was performed using the extra precision (XP) mode in Glide (*82, 83*). Given the flexibility of a 3-mer peptide, numerous high scoring binding conformations were predicted (XP score > 9.5 kcal/mol), so to assess the likely binding conformation, the binding modes were superpositioned with the backbone conformation of the residues of SMRT-DAD (derived from X-ray co-crystal structure) to assess the reasonability of the binding modes and select the most likely binding mode. Finally, the ligand and residues of HDAC3 in 5 Å proximity were refined and re-scored using the MM-GBSA module in Prime (*84*). Figures were generated using PyMol Molecular Graphics System (version 2.3.4, Schrödinger, LLC).

## Supplementary Materials

Fig. S1. Supplementary deacylase activity screening.

Fig. S2. Structure of fluorogenic substrates.

Fig. S3. Bar graphs corresponding to the heat map in Fig. 1D.

Fig. S4. Deacylase efficiencies of HDACs 1–3 against fluorogenic substrates.

Fig. S5. Deacylase efficiencies of HDACs 1–3 against non-fluorogenic substrates.

Fig. S6. Inhibition of HDACs 1–3 by RGFP966.

Fig. S7. Unmodified Western blots corresponding to Fig. 6.

Table S1. List of quantified acetylated and lactylated peptides, relative to Fig. 5.

Supplementary Methods: HDAC rate inhibition assays

Supplementary Methods: Chemical synthesis

## General

we thank Dr. Iacopo Galleano for donation of peptide substrates, Dr. Stephan A. Pless for donation of the HEK293T cell lines applied in the Olsen Lab, and we are grateful to Prof. Phil Cole at Harvard for generous supply of the apicidin compound.

## Author contributions

conceptualization, C.M.-Y., D.Z., M.B., C.A.O., and Y.Z.; methodology, C.M.-Y., D.Z., W.W., M.B., J.G., A.L.N., C.A.O., and Y.Z., formal analysis, C.M.-Y., D.Z., W.W., M.B., and J.G, investigation, C.M.-Y., D.Z., W.W., M.B., J.G., A.L.N., J.E.B., and L.Y. resources, C.M.-Y., D.Z., W.W., A.L.N., J.E.B., and S.T.J., writing – original draft, C.M.-Y., D.Z., C.A.O., and Y.Z., writing – review & editing, all authors, visualization, C.M.-Y., and D.Z., supervision, J.W., C.A.O., and Y.Z., project administration, J.W., C.A.O., and Y.Z., funding acquisition, J.W., C.A.O., and Y.Z.

## Data and materials availability

all data needed to evaluate the conclusions in the paper are present in the paper and/or the Supplementary Materials.

