## Supplementary Material for "Class I Histone Deacetylases (HDAC1–3) are Histone Lysine Delactylases"

Supplementary Figures S2

Supplementary Tables S10

Supplementary Methods S11

Chemical synthesis S11

NMR spectra S28

HPLC purity traces S42

Supplementary References S46

#### Supplementary Figures

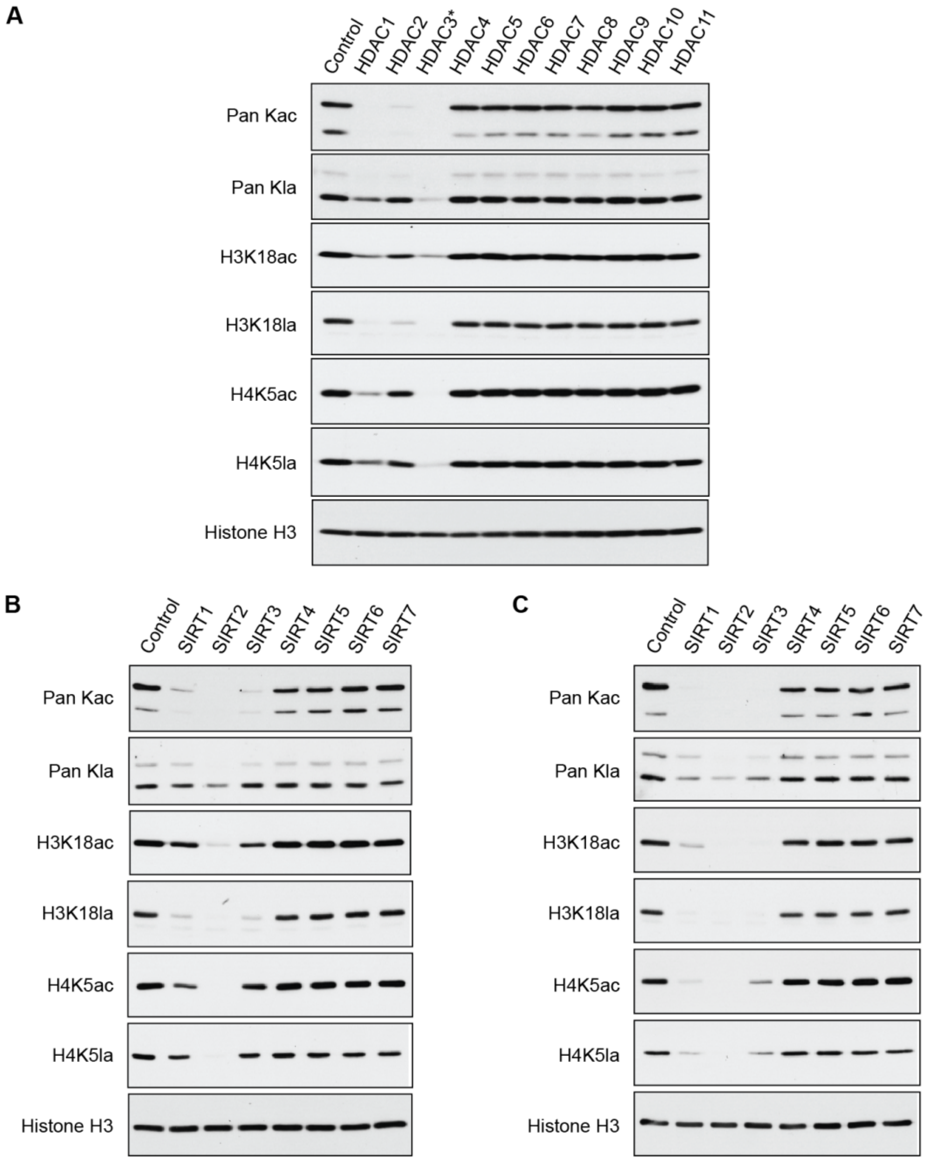


**Fig. S1. Selectivity of the Pan Kla antibody and supplementary deacylase activity screening.** (**A**) Deacylase activity screening using purified histones from HeLa cells and antibodies against Kac and Kla modifications, with histone H3 as loading control (1 h reaction). (**B**) Deacylase activity screening of recombinant sirtuin enzymes using purified histones from HeLa cells and antibodies against Kac and Kla modifications, with histone H3 as loading control (1 h reaction). (**C**) Deacylase activity screening of recombinant sirtuin enzymes using purified histones from HeLa cells (4 h reaction).

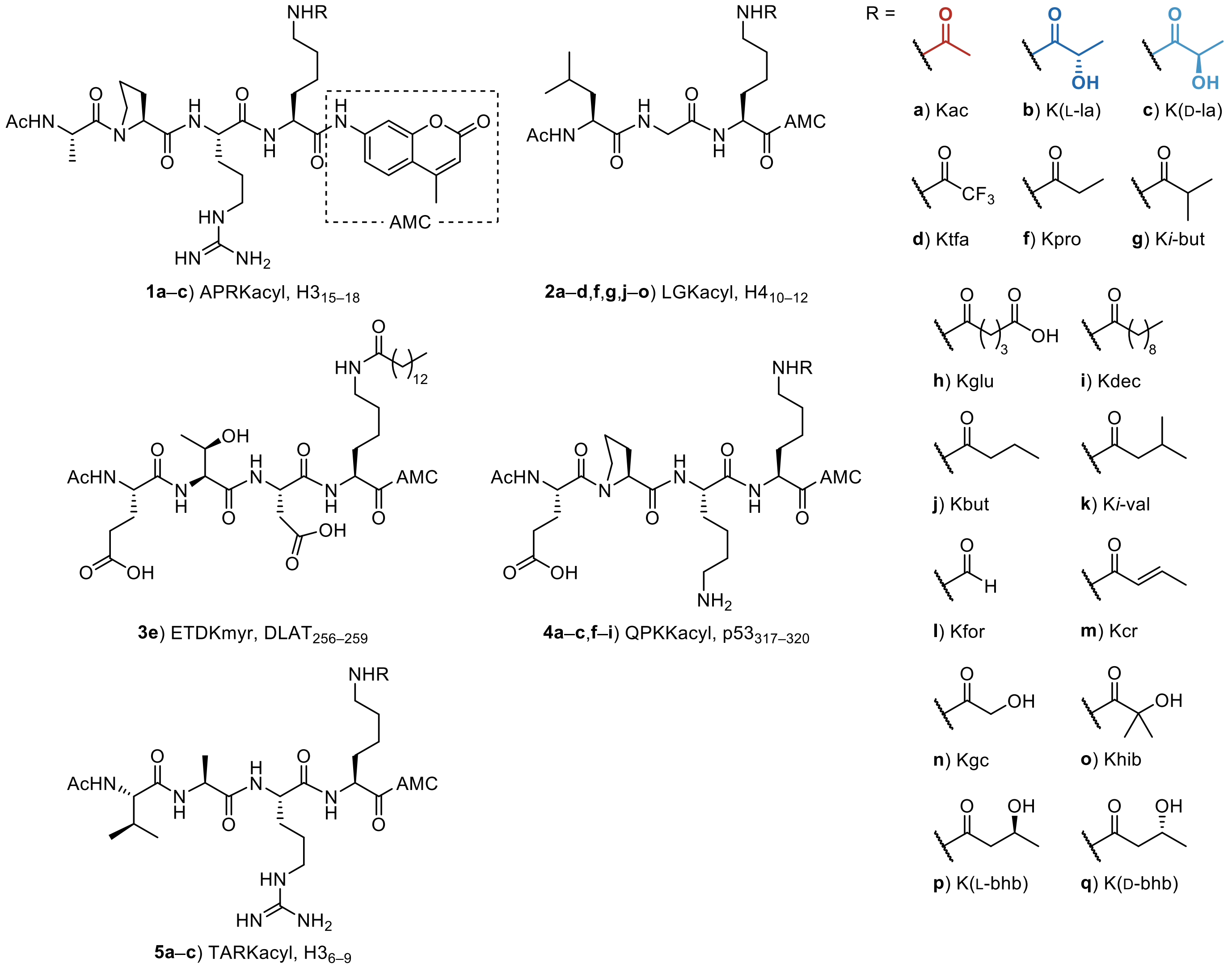


**Fig. S2. Structure of fluorogenic substrates.** AMC: 7-amino-4-methylcoumarin.

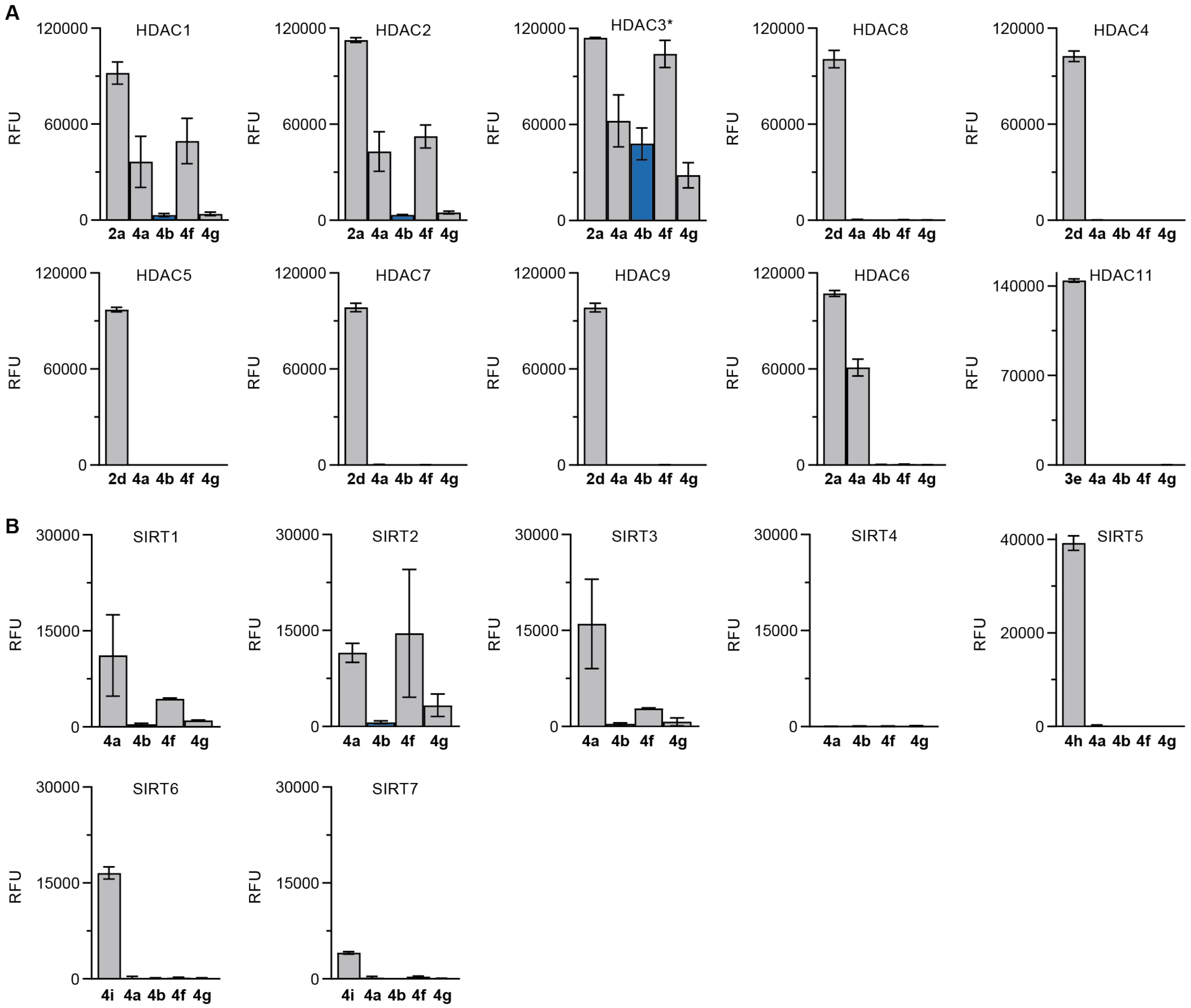


**Fig. S3. Bar graphs corresponding to the heat map in Fig. 1D.**

**A**

| | HDAC1 | | HDAC2 | | HDAC3* | |
|---|---|---|---|---|---|---|
| | LGK(L-la) (**2b**) | LGK(D-la) (**2c**) | LGK(L-la) (**2b**) | LGK(D-la) (**2c**) | LGK(L-la) (**2b**) | LGK(D-la) (**2c**) |
| $K_M$ (µM) | 174 ± 5 | 68 ± 4 | 180 ± 10 | 69 ± 6 | 122 ± 7 | 67 ± 7 |
| $k_{cat}$ ($s^{-1}$) | 0.141 ± 0.002 | 0.35 ± 0.01 | 0.144 ± 0.006 | 0.57 ± 0.02 | 0.49 ± 0.02 | 0.92 ± 0.04 |
| $k_{cat}/K_M$ ($M^{-1}\cdot s^{-1}$) | (0.81 ± 0.02)·$10^3$ | (5.1 ± 0.3)·$10^3$ | (0.80 ± 0.06)·$10^3$ | (8.3 ± 0.7)·$10^3$ | (4.0 ± 0.2)·$10^3$ | (14 ± 1)·$10^3$ |

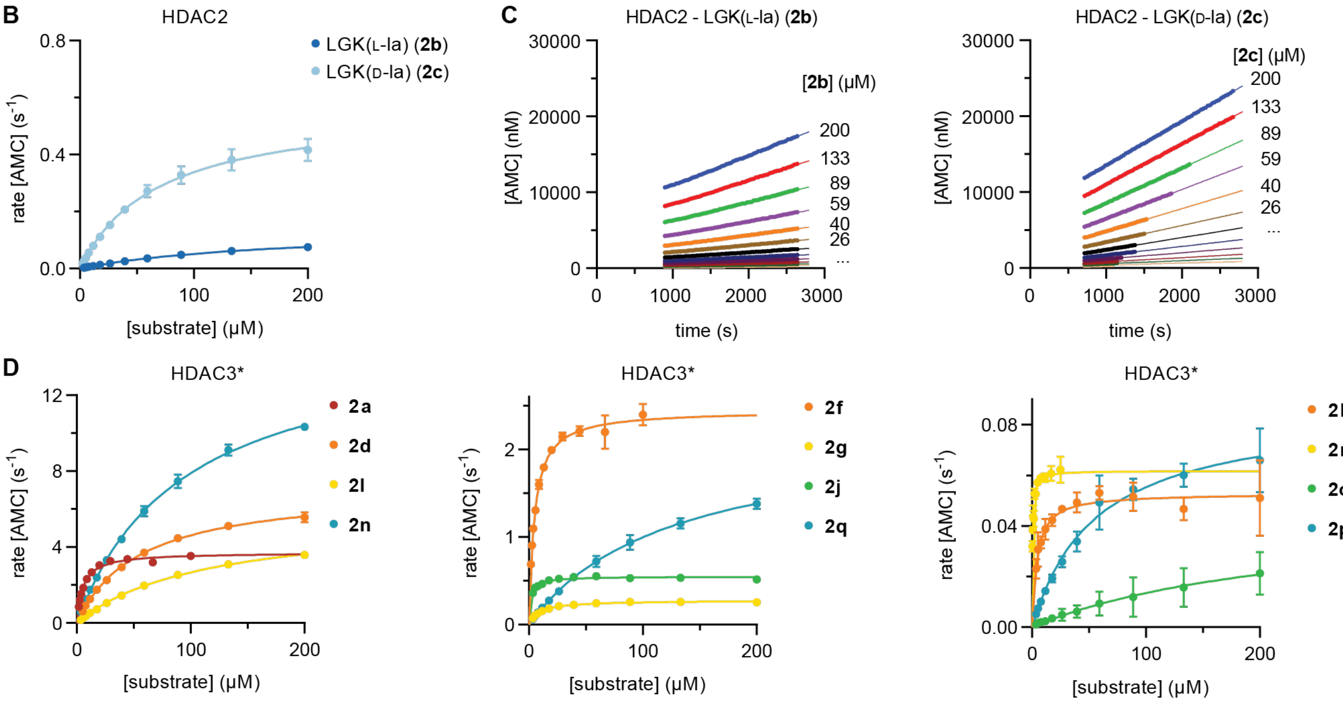


**E**

| HDAC3* | $K_M$ (µM) | $k_{cat}$ ($s^{-1}$) | $k_{cat}/K_M$ ($M^{-1}\cdot s^{-1}$) |
|---|---|---|---|
| LGKac (**2a**) | 5.4 ± 0.3 | 3.72 ± 0.06 | (690 ± 40)·$10^3$ |
| LGK(L-la) (**2b**) | 122 ± 7 | 0.49 ± 0.02 | (4.0 ± 0.2)·$10^3$ |
| LGK(D-la) (**2c**) | 67 ± 7 | 0.92 ± 0.04 | (14 ± 1)·$10^3$ |
| LGKtfa (**2d**) | 56 ± 2 | 7.2 ± 0.1 | (129 ± 5)·$10^3$ |
| LGKpro (**2f**) | 4.7 ± 0.2 | 2.45 ± 0.03 | (520 ± 30)·$10^3$ |
| LGK(*i*-but) (**2g**) | 9.0 ± 0.6 | 0.28 ± 0.01 | (31 ± 2)·$10^3$ |
| LGKbut (**2j**) | 1.7 ± 0.2 | 0.55 ± 0.01 | (310 ± 40)·$10^3$ |
| LGK(*i*-val) (**2k**) | 4.1 ± 0.8 | 0.053 ± 0.002 | (13 ± 2)·$10^3$ |

| HDAC3* | $K_M$ (µM) | $k_{cat}$ ($s^{-1}$) | $k_{cat}/K_M$ ($M^{-1}\cdot s^{-1}$) |
|---|---|---|---|
| LGKfor (**2l**) | 112 ± 3 | 5.7 ± 0.1 | (50 ± 1)·$10^3$ |
| LGKcr (**2m**) | 0.43 ± 0.04 | 0.062 ± 0.001 | (140 ± 10)·$10^3$ |
| LGKgc (**2n**) | 94 ± 4 | 15.3 ± 0.3 | (163 ± 6)·$10^3$ |
| LGKhib (**2o**) | 240 ± 160 | 0.05 ± 0.02 | (0.2 ± 0.1)·$10^3$ |
| LGK(L-bhb) (**2p**) | 56 ± 9 | 0.086 ± 0.006 | (1.5 ± 0.2)·$10^3$ |
| LGK(D-bhb) (**2q**) | 125 ± 9 | 2.25 ± 0.09 | (18 ± 1)·$10^3$ |

**Fig. S4. Deacylase efficiencies of HDACs 1–3 against fluorogenic substrates.** (**A**) Steady-state parameters of HDACs 1–3 against substrates **2b** and **2c** relative to curves presented in Figs. 2B and S4B. (**B**) Michaelis-Menten plots for HDAC2 against substrates **2b** and **2c**. Data represent mean ± SEM, *n* = 2. (**C**) Sample assay progression curves for HDAC2. Only data corresponding to the steady state is included in the analysis. (**D**) Michaelis-Menten plots for HDAC3/NCoR2 against substrates **2a**, **2d**, **2f**, **2g** and **2j–2q**. Data represent mean ± SEM, *n* = 2. (**E**) Steady-state parameters of HDAC3/NCoR2 against fluorogenic substrates relative to bar graphs in Fig. 4B. *HDAC3 incubated with the DAD of NCoR2.

**A**

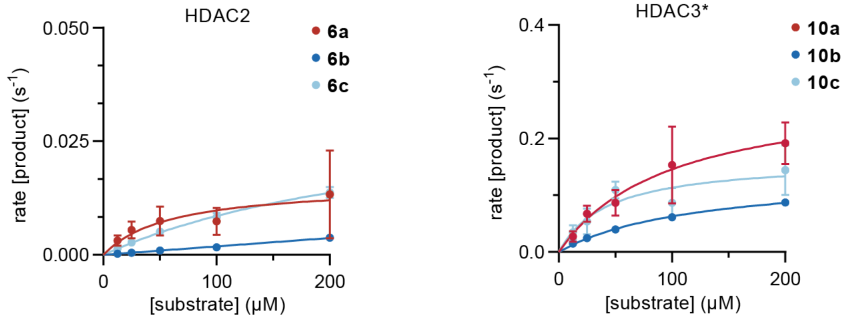


**B**

| | HDAC1 | | | HDAC2 | | |
|---|---|---|---|---|---|---|
| | H3K9ac (**6a**) | H3K9(L-la) (**6b**) | H3K9(D-la) (**6c**) | H3K9ac (**6a**) | H3K9(L-la) (**6b**) | H3K9(D-la) (**6c**) |
| $K_M$ (μM) | 150 ± 50 | 900 ± 700 | 500 ± 200 | 60 ± 30 | > 200 | 300 ± 40 |
| $k_{cat}$ (s$^{-1}$) | 0.07 ± 0.01 | 0.04 ± 0.03 | 0.09 ± 0.03 | 0.015 ± 0.03 | > 0.02 | 0.034 ± 0.003 |
| $k_{cat}/K_M$ (M$^{-1}$·s$^{-1}$) | (0.5 ± 0.2)·10$^3$ | (0.04 ± 0.04)·10$^3$ | (0.20 ± 0.09)·10$^3$ | (0.3 ± 0.1)·10$^3$ | < 0.04·10$^3$ | (0.11 ± 0.01)·10$^3$ |

| | HDAC3* | | | | | |
|---|---|---|---|---|---|---|
| | H3K9ac (**6a**) | H3K9(L-la) (**6b**) | H3K9(D-la) (**6c**) | H4K12ac (**10a**) | H4K12(L-la) (**10b**) | H4K12(D-la) (**10c**) |
| $K_M$ (μM) | 23 ± 8 | 90 ± 20 | 70 ± 20 | 100 ± 20 | 120 ± 10 | 40 ± 30 |
| $k_{cat}$ (s$^{-1}$) | 0.32 ± 0.03 | 0.24 ± 0.02 | 0.42 ± 0.04 | 0.29 ± 0.03 | 0.14 ± 0.01 | 0.16 ± 0.04 |
| $k_{cat}/K_M$ (M$^{-1}$·s$^{-1}$) | (14 ± 5)·10$^3$ | (2.6 ± 0.6)·10$^3$ | (6 ± 1)·10$^3$ | (2.9 ± 0.8)·10$^3$ | (1.1 ± 0.1)·10$^3$ | (4 ± 2)·10$^3$ |

**Fig. S5. Sirtuin delactylase activity and deacylase efficiencies of HDACs 1–3 against non-fluorogenic substrates.** (**A**) Michaelis-Menten plots for HDAC2 against substrates **6a–c** and for HDAC3/NCoR2 against substrates **10a–c**. Data represent mean ± SEM, *n* = 2. (**B**) Steady-state parameters of HDACs 1–3 against non-fluorogenic histone substrates relative to bar graphs in Fig. 3F. *HDAC3 incubated with the DAD of NCoR2.

**A**

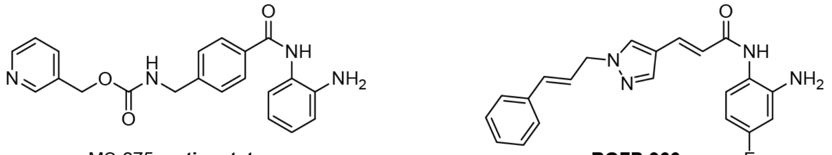


MS-275, **entinostat** **RGFP-966**

**B**

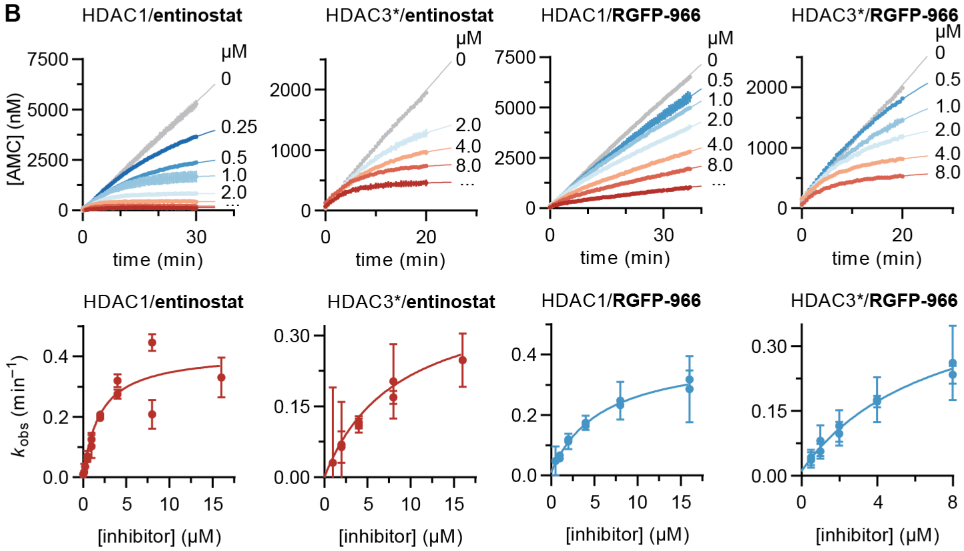


**C**

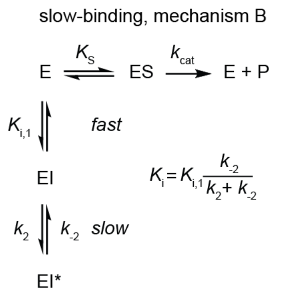


| **Entinostat** | HDAC1 | HDAC2 | HDAC3* |
|---|---|---|---|
| $k_2$ (min$^{-1}$) | 0.41 ± 0.05 | 1.2 ± 0.8 | 0.40 ± 0.07 |
| $k_{-2}$ (min$^{-1}$) | ~1 × 10$^{-8}$ | ~1 × 10$^{-2}$ | ~5 × 10$^{-10}$ |
| $K_{i,1}$ (nM) | 500 ± 200 | 2100 ± 2100 | 2000 ± 1100 |
| $K_i$ (nM) | –† | ~19 | –† |
| Dis. $t_{½}$ (min) | > 10$^5$ | ≥ 62 | > 10$^5$ |
| **RGFP966** | HDAC1 | HDAC2 | HDAC3* |
| $k_2$ (min$^{-1}$) | 0.40 ± 0.02 | 0.55 ± 0.06 | 0.45 ± 0.07 |
| $k_{-2}$ (min$^{-1}$) | (1 ± 1) × 10$^{-2}$ | (1.7 ± 0.9) × 10$^{-2}$ | (1 ± 1) × 10$^{-2}$ |
| $K_{i,1}$ (nM) | 1400 ± 300 | 1700 ± 400 | 1600 ± 600 |
| $K_i$ (nM) | ~35 | ~51 | ~40 |
| Dis. $t_{½}$ (min) | ≥ 66 | ≥ 42 | ≥ 62 |

**Fig. S6. RGFP966 inhibits HDACs 1–3.** (**A**) Structures of the HDAC1–3-selective inhibitor **entinostat** and of **RGFP966**. (**B**) Representative continuous assay progression curves and kinetic data fitting for **entinostat** and **RGFP966** against HDACs 1 and 3. Data fitting represent mean ± SEM, *n* = 2. (**C**) Mechanism B of slow-binding kinetics, and kinetic and equilibrium constants for **entinostat** and **RGFP966** against HDACs 1–3. Data represent mean ± SEM, *n* = 2.

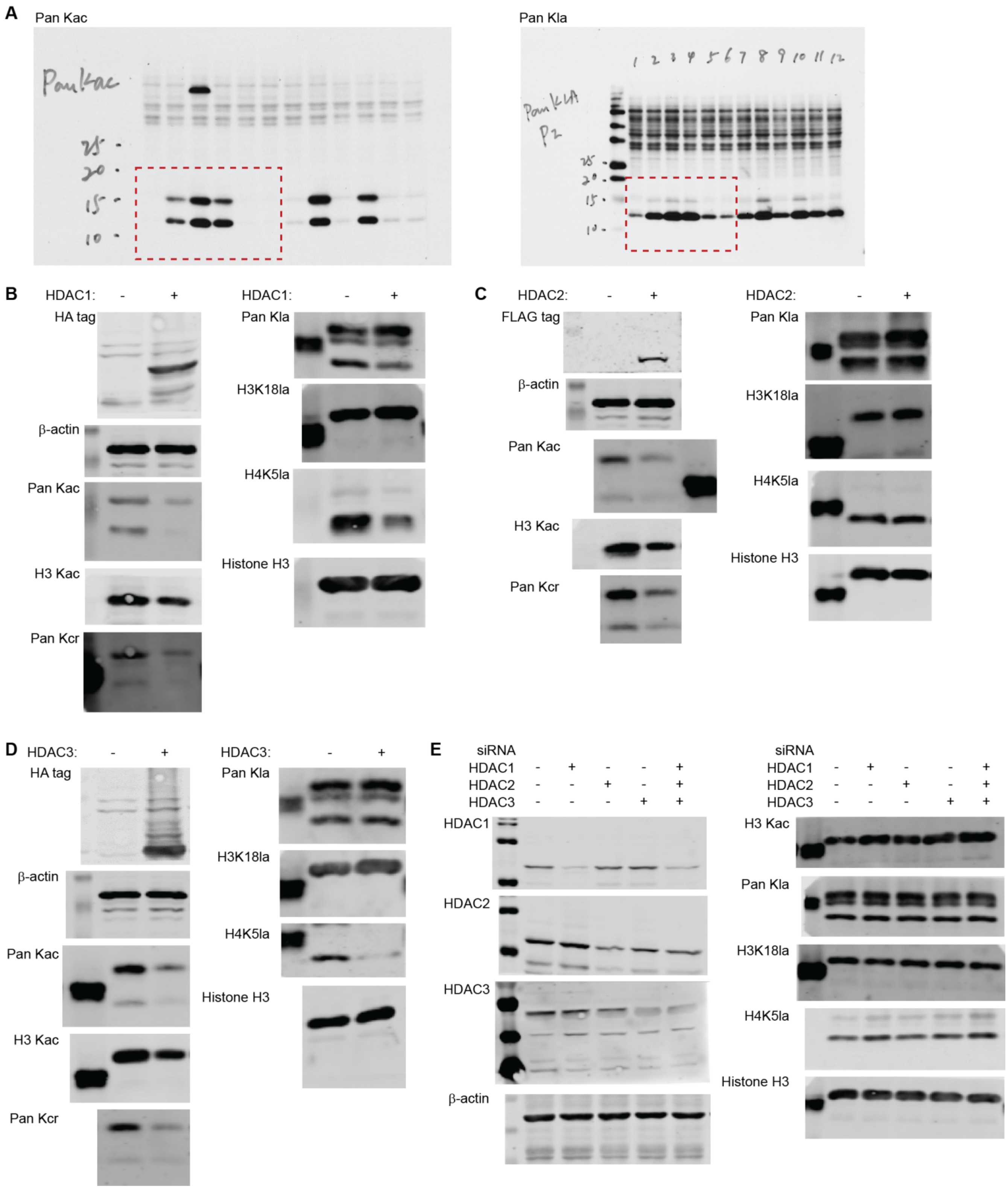


**Fig. S7. Unmodified Western blots corresponding to Fig. 6.** See legend in next page.

**Fig. S7** (contd.) (**A**) Western blots depicted in Fig. 6A. (**B**) Western blots relative to samples transfected with HA-tagged HDAC1. (**C**) Western blots relative to samples transfected with FLAG-tagged HDAC2. (**D**) Western blots relative to samples transfected with HA-tagged HDAC3. (**E**) Western blots relative to samples with or without knockdown of HDAC1, 2, 3, or their combination.

#### Supplementary Tables

**Table S1. List of quantified acetylated and lactylated peptides, relative to Fig. 5.** Symbols represent Kac (K#) and Kla (K*).

| ipi | description | symbol | sequence | mass | charge | Ratio_Forward labeling | Ratio_Reverse labeling |
|---|---|---|---|---|---|---|---|
| P68431 | HIST1H3A Histone H3.1 | HIST1H3A | R.K#QLATK*AAR.K | 1099,635 | 2 | 0,07 | 15 |
| P04908 | HIST1H2AB Histone H2A type 1-B/E | HIST1H2AB | R.GK#QGGK*AR.A | 914,4934 | 2 | 0,07 | 11,99 |
| O60814 | HIST1H2BK Histone H2B type 1-K | HIST1H2BK | K.SAPAPK*K#GSK.K | 1083,592 | 2 | 0,07 | 14,82 |
| P62805 | HIST1H4A Histone H4 | HIST1H4A | R.GKGGK*GLGK.G | 872,508 | 2 | 0,07 | 14,9 |
| O60814 | HIST1H2BK Histone H2B type 1-K | HIST1H2BK | K.K#GSK#K*AVTK.A | 1101,639 | 2 | 0,07 | 15 |
| P62805 | HIST1H4A Histone H4 | HIST1H4A | K.GGK*GLGK#GGAK.R | 1042,577 | 2 | 0,07 | 15 |
| P62805 | HIST1H4A Histone H4 | HIST1H4A | K.GGK*GLGK.G | 687,3915 | 2 | 0,07 | 15 |
| P62805 | HIST1H4A Histone H4 | HIST1H4A | K.GLGK*GGAK#R.H | 956,5403 | 2 | 0,07 | 15 |
| P62805 | HIST1H4A Histone H4 | HIST1H4A | R.GK#GGK*GLGK#GGAK#R.H | 1467,816 | 2 | 0,07 | 15 |
| P62805 | HIST1H4A Histone H4 | HIST1H4A | R.GK#GGK*GLGK#GGAK.R | 1269,704 | 2 | 0,07 | 15 |
| P62805 | HIST1H4A Histone H4 | HIST1H4A | R.GK#GGK*GLGK.G | 914,5185 | 2 | 0,07 | 15 |
| Q5QNW6-2 | HIST2H2BF Isoform 2 of Histone H2B type 2-F | HIST2H2BF | K.K#GSK#K*AVTK.V | 1101,639 | 2 | 0,07 | 15 |
| O60814 | HIST1H2BK Histone H2B type 1-K | HIST1H2BK | -.PEPAK#SAPAPK*K.G | 1333,724 | 2 | 0,07 | NA |
| O60814 | HIST1H2BK Histone H2B type 1-K | HIST1H2BK | K.AVTK*AQK.K | 816,4705 | 2 | 0,07 | NA |
| O60814 | HIST1H2BK Histone H2B type 1-K | HIST1H2BK | K.KAVTK*AQK.K | 944,5655 | 2 | 0,07 | NA |
| P62805 | HIST1H4A Histone H4 | HIST1H4A | K.GGK#GLGK*GGAK#R.H | 1240,689 | 2 | 0,07 | 15 |
| P62805 | HIST1H4A Histone H4 | HIST1H4A | K.GGK#GLGK*GGAK.R | 1042,577 | 2 | 0,07 | 15 |
| P62805 | HIST1H4A Histone H4 | HIST1H4A | K.GGK*GLGK#GGAK#R.H | 1240,689 | 2 | 0,07 | 15 |
| P62805 | HIST1H4A Histone H4 | HIST1H4A | R.GK*GGK#GLGK#GGAK#R.H | 1467,816 | 3 | 0,07 | 15 |
| P62805 | HIST1H4A Histone H4 | HIST1H4A | R.GKGGK#GLGK*GGAK#R.H | 1425,805 | 2 | 0,07 | 15 |
| P68431 | HIST1H3A Histone H3.1 | HIST1H3A | R.K#STGGK*APR.K | 1014,546 | 2 | 0,07 | NA |
| P68431 | HIST1H3A Histone H3.1 | HIST1H3A | R.KQLATK*AAR.K | 1057,624 | 2 | 0,07 | 15 |
| P68431 | HIST1H3A Histone H3.1 | HIST1H3A | R.KQLATK*AAR.K | 1057,624 | 3 | 0,07 | 15 |
| Q5QNW6-2 | HIST2H2BF Isoform 2 of Histone H2B type 2-F | HIST2H2BF | K.K#AVTK*VQK.K | 1014,607 | 2 | 0,07 | NA |
| P68431 | HIST1H3A Histone H3.1 | HIST1H3A | R.K*STGGK#APR.K | 1014,546 | 2 | 0,07 | 15 |
| P62805 | HIST1H4A Histone H4 | HIST1H4A | R.GK#GGKGLGK*GGAK#R.H | 1425,805 | 3 | 0,08 | 15 |
| Q99880 | HIST1H2BL Histone H2B type 1-L | HIST1H2BL | -.PELAK*SAPAPK.K | 1179,65 | 2 | 0,09 | 2,5 |
| Q5QNW6-2 | HIST2H2BF Isoform 2 of Histone H2B type 2-F | HIST2H2BF | K.AVTK*VQK.K | 844,5018 | 2 | 0,09 | 4,42 |
| P0C0S5 | H2AFZ Histone H2A.Z | H2AFZ | -.AGGK#AGK#DSGK*AK.T | 1329,689 | 2 | 0,1 | 4,74 |
| O60814 | HIST1H2BK Histone H2B type 1-K | HIST1H2BK | K.SAPAPK*K.G | 769,4334 | 2 | 0,14 | 2,02 |
| Q5QNW6-2 | HIST2H2BF Isoform 2 of Histone H2B type 2-F | HIST2H2BF | K.KAVTK*VQK.K | 972,5968 | 2 | 0,14 | 2,15 |
| Q99879 | HIST1H2BM Histone H2B type 1-M | HIST1H2BM | -.PEPVK*SAPVPK.K | 1219,681 | 2 | 0,21 | 2,11 |
| O60814 | HIST1H2BK Histone H2B type 1-K | HIST1H2BK | -.PEPAK*SAPAPK.K | 1163,619 | 3 | 0,25 | NA |
| P23527 | HIST1H2BO Histone H2B type 1-O | HIST1H2BO | -.PDPAK*SAPAPK.K | 1149,603 | 2 | 0,25 | NA |
| O60814 | HIST1H2BK Histone H2B type 1-K | HIST1H2BK | -.PEPAK*SAPAPK.K | 1163,619 | 2 | 0,26 | 1,53 |
| O60814 | HIST1H2BK Histone H2B type 1-K | HIST1H2BK | K.K#GSK*K#AVTK#AQK.K | 1470,841 | 3 | 0,28 | NA |
| P68431 | HIST1H3A Histone H3.1 | HIST1H3A | K.QLATK*AAR.K | 929,5294 | 2 | NA | 15 |
| O60814 | HIST1H2BK Histone H2B type 1-K | HIST1H2BK | K.K#AVTK*AQK.K | 986,576 | 2 | NA | 15 |
| P58876 | HIST1H2BD Histone H2B type 1-D | HIST1H2BD | -.PEPTK*SAPAPK.K | 1193,629 | 2 | NA | 2,2 |

### Supplementary Methods

#### HDAC rate inhibition assays

Kinetic HDAC inhibition assays were performed in Tris buffer [50 mM Tris/Cl, 137 mM NaCl, 2.7 mM KCl, 1 mM MgCl, pH 8] with 0.5 mg/mL BSA. For a final volume of 50 µL, substrate (**2a**, 20 µM), trypsin (10 µg/mL), inhibitor (16000–31.2 nM, 2-fold dilutions) and enzyme (HDAC1: 3.5 nM, HDAC2: 1.8 nM, HDAC3: 0.8 nM) were added to a microtiter plate and immediately placed in the plate reader. *In situ* fluorophore release was monitored by fluorescence readings every 30 s for 60 min at 25 °C. Then, apparent first-order rate constants for establishment of the enzyme-inhibitor equilibrium ($k_{obs}$) and inhibitor concentration ([I]) were fitted to mechanism B of slow kinetics (hyperbolic relationship, **Eq. S1**). $K_i$ and dissociative half-live (Dis. $t_{½}$) values were estimated when possible using **Eq. S2** and **Eq. S3**, respectively (*1, 2*).

$$k_{\mathrm{obs}} = \frac{k_2}{[\mathrm{I}] + K_{\mathrm{i},1}\left(1 + \frac{[\mathrm{S}]}{K_{\mathrm{M}}}\right)}[\mathrm{I}] + k_{-2} \quad (\mathbf{Eq.\,S1})$$

$$K_{\mathrm{i}} = K_{\mathrm{i},1}\frac{k_2}{k_2 + k_{-2}} \quad (\mathbf{Eq.\,S2})$$

$$\mathrm{Dis.\,t_{½}} = \frac{\ln(2)}{\frac{k_{-1} \cdot k_{-2}}{(k_{-1} + k_2 + k_{-2})}} \quad (\mathbf{Eq.\,S3})$$

#### Chemical synthesis

##### *General methods*

All commercial reagents and solvents were of analytical grade and used without further purification. Anhydrous solvents were obtained from a PureSolv system. Reactions were conducted under an atmosphere of nitrogen whenever anhydrous solvents were used. Reactions were monitored by thin-layer chromatography (TLC) using silica gel-coated plates (analytical $SiO_2$-60, F-254) and by HPLC-MS analysis. TLC plates were visualized under UV light and/or by staining with (a) a solution of potassium permanganate (10 g/L), potassium carbonate (67 g/L) and sodium hydroxide (0.83 g/L) in water, (b) a solution of ninhydrin (3 g/L) in 3% acetic acid in water (v/v), or (c) a solution of molybdate-phosphoric acid (12.5 g/L) and cerium(IV)sulfate (5 g/L) in 3% conc. sulfuric acid in water (v/v). Evaporation of solvents was carried out under reduced pressure at a temperature below 40 °C. HPLC−MS analyses were performed on a Phenomenex Kinetex column (1.7 µm, 50×2.10 mm) using a Waters Acquity ultra high-performance liquid chromatography (UPLC) system. Gradient A with eluent I (0.1% HCOOH in $H_2O$) and eluent II (0.1% HCOOH in MeCN) rising linearly from 0% to 95% of II during *t* = 0.00-5.20 min was applied at a flow rate of 0.6 mL/min. Preparative reversed-phase HPLC purification was performed on a C18 Phenomenex® Luna column (5 µm, 250×20 mm) or a C8(2) Phenomenex® Luna column (5 µm, 250×21.2 mm) using an Agilent 1260 LC system equipped with a diode array UV detector and an evaporative light scattering detector (ELSD). Gradient B with eluent III ($H_2O$/MeCN/TFA, 95:5:0.1, v:v) and eluent IV (0.1% TFA in MeCN) rising linearly from 0-30% to 95% of IV during *t* = 5−45 min was applied at a flow rate of 20 mL/min. Analytical HPLC was performed on a C18 Infinity Poroshell 120 column (2.7 µm, 100×3.0 mm) using an Agilent 1260 Infinity II series system equipped with a diode array UV detector using eluent III and eluent IV, rising linearly from 0% to 50% or 95% of IV during $t = 1{-}11$ was applied at a flow rate of 1.2 mL/min at 40 °C. Nuclear magnetic resonance (NMR) spectra were recorded on a Bruker Avance III HD equipped with a cryogenically cooled probe ($^{1}$H NMR and $^{13}$C NMR recorded at 600 and 151 MHz, respectively). All spectra were recorded at 298 K. Chemical shifts are reported in ppm relative to deuterated solvent as internal standard ($\delta_H$ DMSO-$d_6$ 2.50 ppm; $\delta_C$ DMSO-$d_6$ 39.52 ppm). Assignments of NMR spectra are based on 2D correlation spectroscopy (COSY, HSQC, TOCSY and HMBC spectra). High-resolution mass spectrometry (HRMS) was recorded either on a QExactive Orbitrap mass spectrometer (Thermo Scientific, Bremen, Germany) equipped with a SMALDI5 ion source (TransMIT GmbH, Giessen, Germany), or on a Bruker Solarix WR by either matrix assisted laser desorption/ionization (MALDI) or electrospray ionization (ESI).

*Synthesis of AMC-coupled fluorogenic substrates*

Multiple fluorogenic substrate building blocks were synthesized following procedures reported in the literature (Scheme S1), and the Ac-Ala-Pro-Arg(Pbf)-Lys-AMC·TFA building block (**S5**) was prepared following similar procedures (Scheme S2). Then, substrates were obtained by acylation in solution (Schemes S3–S6).

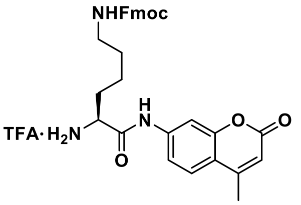

H-Lys(Fmoc)-AMC·TFA
**S1**

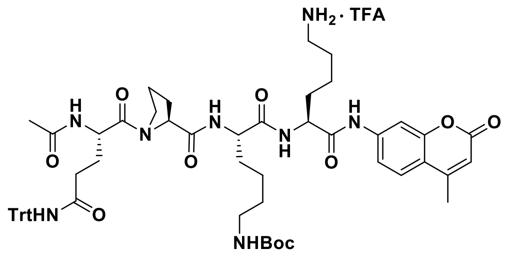

Ac-Gln(Trt)-Pro-Lys(Boc)-Lys-AMC·TFA
**S2**

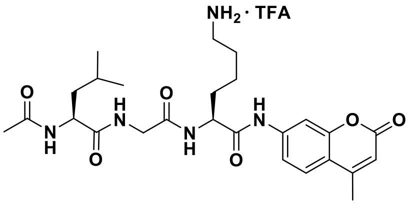

Ac-Leu-Gly-Lys-AMC·TFA
**S3**

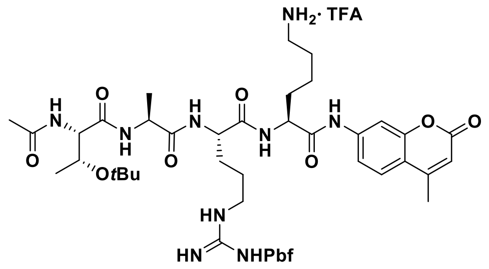

Ac-Thr(*t*Bu)-Ala-Arg(Pbf)-Lys-AMC·TFA
**S4**

**Scheme S1. Building blocks used for the synthesis of AMC-coupled substrates.** Compounds **S1** (*3*), **S2** (*4*), **S3** (*5*), **S4** (*6*) were synthesized as published.

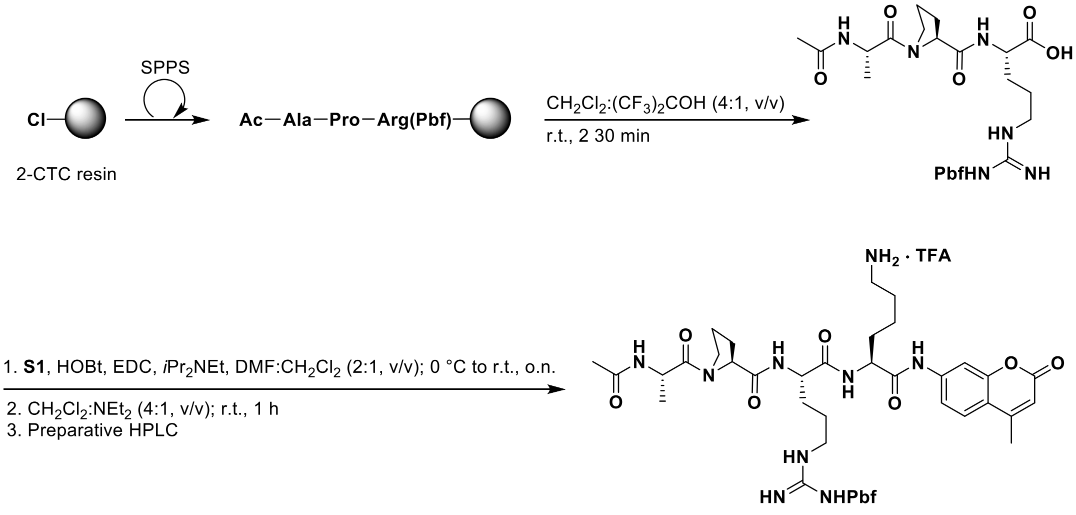


Ac-Ala-Pro-Arg(Pbf)-Lys-AMC · TFA
**S5**
(37% from resin loading)

**Scheme S2. Synthesis of building block S5.**

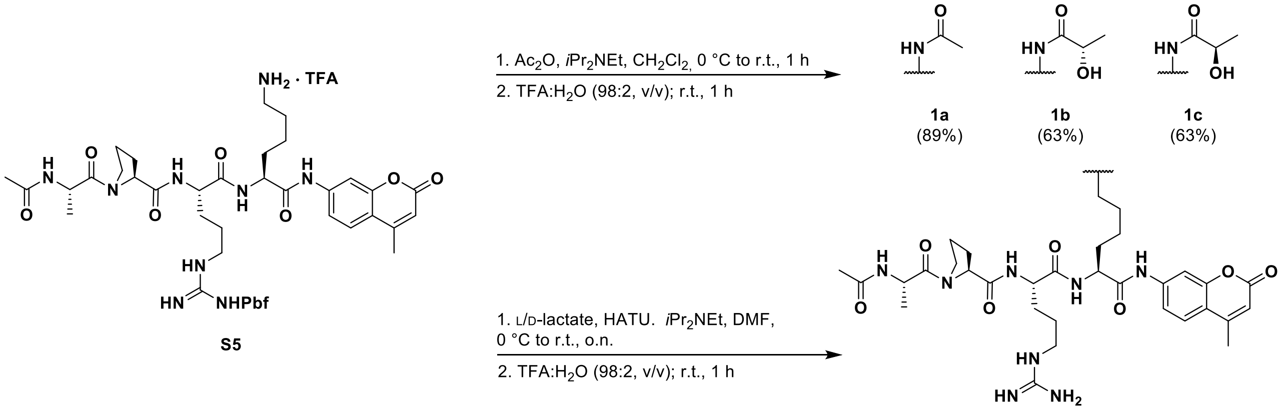


**Scheme S3. Synthesis of substrates 1a–1c.**

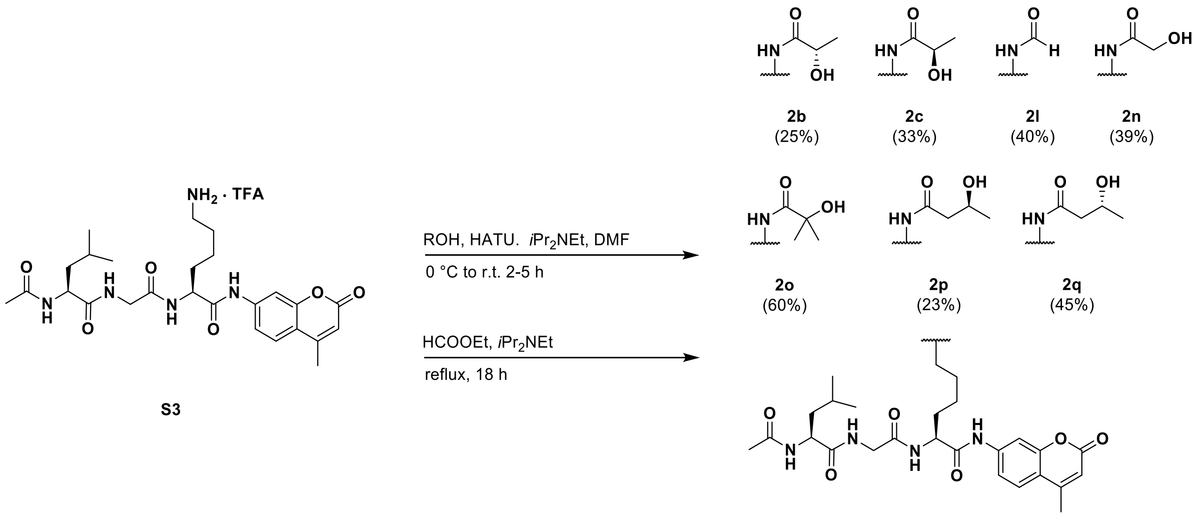


**Scheme S4. Synthesis of substrates 2b, 2c, 2l, 2n, 2o, 2p and 2q.**

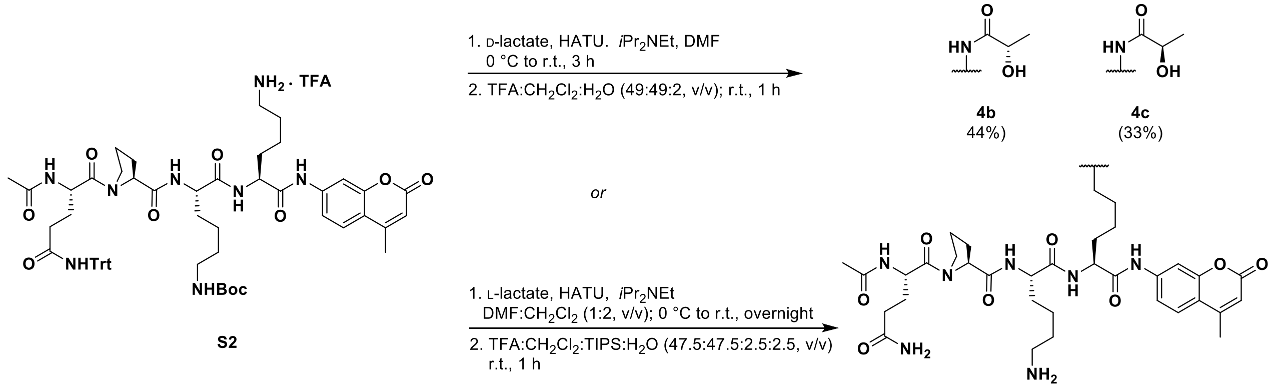


**Scheme S5. Synthesis of substrates 4b and 4c.**

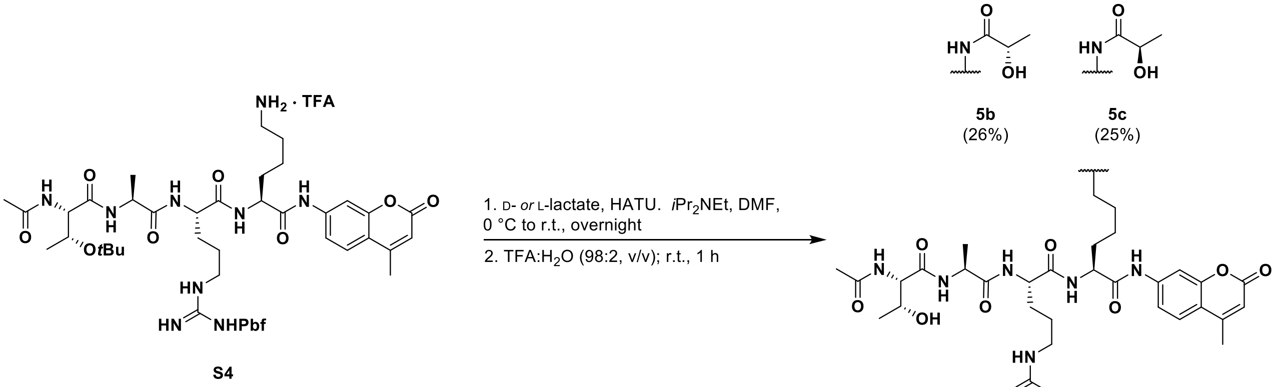


**Scheme S6. Synthesis of substrates 5b and 5c.**

**Ac-Ala-Pro-Arg(Pbf)-Lys-AMC (S5).** Ac-Ala-Pro-Arg(Pbf)-resin (232 µmol resin loading) was synthesized on 2-chlorotrityl chloride (2-CTC) resin by using standard SPPS procedures as previously described.[1] The peptide was cleaved off the resin with $CH_2Cl_2$/hexafluoroisopropanol (4:1, v/v, 2×4 mL, 2×30 min) and concentrated under reduced pressure. Excess hexafluoroisopropanol was removed by co-evaporation with $CH_2Cl_2$:toluene (1:1, v/v, 15 mL) affording the crude 3-mer tentatively assigned as Ac-Ala-Pro-Arg(Pbf)-OH (HPLC-MS $t_R$ 1.38 min, *m*/*z* 637.3; $[M+H]^+$, $C_{29}H_{45}N_6O_8S^+$, Calcd 637.3) as a white solid (166 mg, quant.), which was used without further purification. The crude was redissolved in anh. $CH_2Cl_2$/DMF (2:1, v/v, 6 mL) and cooled to 0 °C. Compound **S1** (144 mg, 0.23 mmol), HOBt (38 mg, 0.28 mmol), *i*$Pr_2$NEt (121 µL, 0.70 mmol) and then EDC (53 mg, 0.28 mmol) was added to the reaction mixture, which was stirred for 10 min at 0 °C and then overnight going towards ambient temperature. The reaction mixture was diluted with $CH_2Cl_2$ (20 mL) and washed with brine (20 mL), 5% $KHSO_4$ (3×20 mL), saturated aq. $NaHCO_3$ (2×20 mL), and brine (20 mL). The organic phase was dried over $Na_2SO_4$ and concentrated under reduced pressure, affording a crude intermediate tentatively assigned as Ac-Ala-Pro-Arg(Pbf)-Lys(Fmoc)-AMC (HPLC-MS $t_R$ 2.10 min, *m*/*z* 1144.6; $[M+H]^+$, $C_{60}H_{74}N_9O_{12}S^+$, Calcd 1144.5) as a white solid (297 mg), which was used without further purification. The crude was redissolved in $CH_2Cl_2$ (4.0 mL) and diethyl amine (1.0 mL) was added dropwise to the reaction mixture, which was stirred for 1 h at ambient temperature. Solvent was removed under reduced pressure, and preparative reversed-phase HPLC purification afforded the desired amine **S5** (87 mg, 37% based on resin loading), as a white fluffy TFA-salt after lyophilization. HPLC-MS $t_R$ 1.31 min, *m*/*z* 922.5 ($[M+H]^+$, $C_{45}H_{64}N_9O_{10}S^+$, Calcd 922.4). AMC=7-amino-4-methylcoumarin.

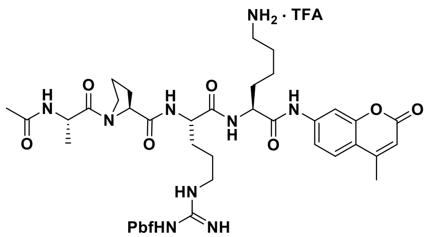


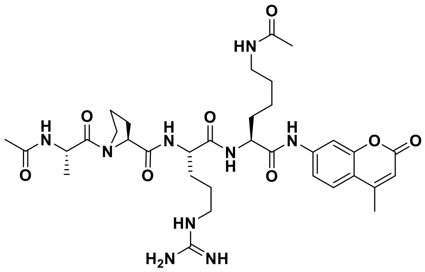

**Ac-Ala-Pro-Arg-Lys(Ac)-AMC (1a).** Compound **S5** (13 mg, 0.013 mmol) was dissolved in anh. $CH_2Cl_2$ (2.0 mL) and cooled to 0 °C. *i*$Pr_2$NEt (7 µL, 0.038 mmol) and acetic anhydride (0.019 mmol) was added to the reaction mixture, which was stirred for 1 h going towards ambient temperature. Solvent was removed under reduced pressure to afford the acetylated intermediate, tentatively assigned as Ac-Ala-Pro-Arg(Pbf)-Lys(Ac)-AMC (HPLC-MS $t_R$ 1.64 min, *m*/*z* 964.5 ($[M+H]^+$, $C_{47}H_{65}N_9O_{11}S^+$, Calcd 964.5), which was used without further purification. TFA:$H_2O$ (98:2, v/v, 1.0 mL) was added to the intermediate and stirred for 1 h at ambient temperature. Solvent was removed under a stream of nitrogen, and preparative reversed-phase HPLC purification afforded the title compound (8 mg, 89% from **S5**) as a white fluffy material after lyophilization. $^1$H NMR (600 MHz, DMSO-$d_6$) δ 10.32 (s, 1H, $NH_{AMC}$), 8.10 (d, *J* = 7.3 Hz, 1H, $NH_{Ala}$), 8.04 (d, *J* = 7.4 Hz, 1H, $NH_{\alpha,Lys}$), 8.01 (d, *J* = 7.5 Hz, 1H, $NH_{\alpha,Arg}$), 7.86–7.78 (m, 2H, $NH_{\varepsilon,Lys}$, $H8_{AMC}$), 7.72 (d, *J* = 8.7 Hz, 1H, $H5_{AMC}$), 7.55 (t, *J* = 5.9 Hz, 1H, $NH_{\delta,Arg}$), 7.50 (dd, J = 8.7, 2.1 Hz, 1H, $H6_{AMC}$), 6.27 (d, *J* = 1.3 Hz, 1H, $H3_{AMC}$), 4.52 (p, *J* = 7.1 Hz, 1H, $H_{\alpha,Ala}$), 4.41–4.29 (m, 2H, $H_{\alpha,Pro}$, $H_{\alpha,Lys}$), 4.29–4.23 (m, 1H, $H_{\alpha,Arg}$), 3.71–3.52 (m, 2H, $H_{\delta,Pro}$), 3.11 (q, *J* = 6.4 Hz, 2H, $H_{\delta,Arg}$), 3.07–2.95 (m, 2H, $H_{\varepsilon,Lys}$), 2.40 (s, 3H, $CH_{3,AMC}$), 2.10–2.00 (m, 1H, $H_{\beta,Pro,A}$), 1.95–1.20 (m, 19H, $H_{\beta,Pro,A}$, $H_{\gamma,Pro}$, $H_{\beta,Lys}$, $H_{\gamma,Lys}$, $H_{\delta,Lys}$, $H_{\beta,Arg}$, $H_{\gamma,Arg}$, $COCH_{3,acetyl}$, $COCH_{3,Lys}$), 1.17 (d, *J* =

6.8 Hz, 3H, $H_{\beta,Ala}$). $^{13}C$ NMR (151 MHz, DMSO) δ 171.9 ($CO_{\alpha,Pro}$), 171.5 ($CO_{Arg}$), 171.3 ($CO_{Ala}$), 171.2 ($CO_{\alpha,Lys}$), 169.0 ($\underline{C}ONH_{\varepsilon,Lys}$), 168.8 ($\underline{C}OCH_{3,acetyl}$), 160.0 ($C2_{AMC}$), 158.4 (q, $J$ = 34.6 Hz, residual $CO_{TFA}$), 156.7 ($NHC(=NH)NH_2$), 153.6 ($C8a_{AMC}$), 153.1 ($C4_{AMC}$), 142.1 ($C7_{AMC}$), 126.0 ($C5_{AMC}$), 116.5 (q, $J$ = 294.6 Hz, residual $CF_{3,TFA}$), 115.2 ($C6_{AMC}$), 115.1 ($C4a_{AMC}$), 112.4 ($C3_{AMC}$), 105.7 ($C8_{AMC}$), 59.6 ($C_{\alpha,Pro}$), 53.7 ($C_{\alpha,Lys}$), 52.2 ($C_{\alpha,Arg}$), 46.8 ($C_{\delta,Pro}$), 46.2 ($C_{\alpha,Ala}$), 40.4 ($C_{\delta,Arg}$), 38.3 ($C_{\varepsilon,Lys}$), 31.3 ($C_{\beta,Lys}$), 29.0 ($C_{\beta,Pro}$), 28.8 ($C_{\beta,Arg}$, $C_{\delta,Lys}$), 25.0 ($C_{\gamma,Arg}$), 24.5 ($C_{\gamma,Pro}$), 22.9 ($C_{\gamma,Lys}$), 22.6 ($CO\underline{C}H_{3,Lys}$), 22.2 ($CO\underline{C}H_{3,acetyl}$), 18.0 ($CH_{3,AMC}$), 16.9 ($C_{\beta,Ala}$). Two sets of signals (approximately 4:1) were detectable due to rotamers. Only peaks for the major rotamer is given. Analytical HPLC gradient 0–95% eluent II in eluent I (11 min total runtime), $t_R$ 4.20 min (>98%, $UV_{230}$). HRMS calcd for $C_{34}H_{50}N_9O_8^+$ $[M+H]^+$, 712.3777; found 712.3776. AMC=7-amino-4-methylcoumarin.

**Ac-Ala-Pro-Arg-Lys(L-La)-AMC (1b).** By the method described for **1c**, the title compound was synthesized using **S5** (18 mg, 0.018 mmol), L-lactate (12 mg, 0.135 mmol), $iPr_2NEt$ (45 µL, 0.270 mmol), and HATU (13 mg, 0.034 mmol). Preparative reversed-phase HPLC purification afforded the title compound (12 mg, 89% from **S5**) as a white fluffy material after lyophilization. $^1H$ NMR (600 MHz, DMSO-$d_6$) δ 10.32 (s, 1H, $NH_{AMC}$), 8.10 (d, $J$ = 7.5 Hz, 1H, $NH_{Ala}$), 8.05 (d, $J$ = 7.3 Hz, 1H, $NH_{\alpha,Lys}$), 8.01 (d, $J$ = 7.6 Hz, 1H, $NH_{\alpha,Arg}$), 7.80 (d, $J$ = 2.0 Hz, 1H, $H8_{AMC}$), 7.72 (d, $J$ = 8.7 Hz, 1H, $H5_{AMC}$), 7.66 (t, $J$ = 5.5 Hz, 1H, $NH_{\varepsilon,Lys}$), 7.55 (t, $J$ = 5.9 Hz, 1H, $NH_{\delta,Arg}$), 7.50 (dd, $J$ = 8.7, 2.1 Hz, 1H, $H6_{AMC}$), 6.27 (d, $J$ = 1.3 Hz, 1H, $H3_{AMC}$), 4.52 (p, $J$ = 7.0 Hz, 1H, $H_{\alpha,Ala}$), 4.40–4.29 (m, 2H, $H_{\alpha,Pro}$, $H_{\alpha,Lys}$), 4.26 (td, $J$ = 7.9, 5.3 Hz, 1H, $H_{\alpha,Arg}$), 3.92 (qd, $J$ = 6.8, 1.6 Hz, 1H, $C(OH)\underline{H}CH_3$), 3.67–3.58 (m, 1H, $H_{\delta,Pro,A}$), 3.59–3.51 (m, 1H, $H_{\delta,Pro,B}$), 3.11 (q, $J$ = 6.7 Hz, 2H, $H_{\delta,Arg}$), 3.08–3.00 (m, 2H, $H_{\varepsilon,Lys}$), 2.40 (d, $J$ = 1.3 Hz, 3H, $CH_{3,AMC}$), 2.10–2.00 (m, 1H, $H_{\beta,Pro,A}$), 1.95–1.21 (m, 16H, $H_{\beta,Pro,A}$, $H_{\gamma,Pro}$, $H_{\beta,Lys}$, $H_{\gamma,Lys}$, $H_{\delta,Lys}$, $H_{\beta,Arg}$, $H_{\gamma,Arg}$, $CH_{3,acetyl}$), 1.20–1.10 (m, 6H, $C(OH)HC\underline{H}_3$, $H_{\beta,Ala}$). $^{13}C$ NMR (151 MHz, DMSO) δ 174.4 ($\underline{C}ONH_{\varepsilon,Lys}$), 171.8 ($CO_{Pro}$), 171.5 ($CO_{Arg}$), 171.3 ($CO_{Ala}$), 171.2 ($CO_{\alpha,Lys}$), 168.8 ($\underline{C}OCH_{3,acetyl}$), 160.0 ($C2_{AMC}$), 158.4 (q, $J$ = 35.1 Hz, residual $CO_{TFA}$), 156.7 ($NHC(=NH)NH_2$), 153.6 ($C8a_{AMC}$), 153.1 ($C4_{AMC}$), 142.1 ($C7_{AMC}$), 125.9 ($C5_{AMC}$), 116.0 (q, $J$ = 293.8 Hz, residual $CF_{3,TFA}$), 115.2 ($C6_{AMC}$), 115.1 ($C4a_{AMC}$), 112.4 ($C3_{AMC}$), 105.7 ($C8_{AMC}$), 67.2 ($\underline{C}(OH)HCH_3$), 59.6 ($C_{\alpha,Pro}$), 53.7 ($C_{\alpha,Lys}$), 52.2 ($C_{\alpha,Arg}$), 46.8 ($C_{\delta,Pro}$), 46.2 ($C_{\alpha,Ala}$), 40.4 ($C_{\delta,Arg}$), 37.9 ($C_{\varepsilon,Lys}$), 31.3 ($C_{\beta,Lys}$), 29.0 ($C_{\beta,Pro}$), 28.93 ($C_{\delta,Lys}$), 28.85 ($C_{\beta,Arg}$), 25.0 ($C_{\gamma,Arg}$), 24.5 ($C_{\gamma,Pro}$), 22.8 ($C_{\gamma,Lys}$), 22.2 ($CO\underline{C}H_{3,acetyl}$), 21.1 ($C(OH)H\underline{C}H_3$), 18.0 ($CH_{3,AMC}$), 16.9 ($C_{\beta,Ala}$). Two sets of signals (approximately 6:1) were detectable due to rotamers. Only peaks for the major rotamer is given. Analytical HPLC gradient 0–95% eluent II in eluent I (11 min total runtime), $t_R$ 4.17 min (>98%, $UV_{230}$). HRMS calcd for $C_{35}H_{52}N_9O_9^+$ $[M+H]^+$, 742.3882; found 742.3884. AMC=7-amino-4-methylcoumarin.

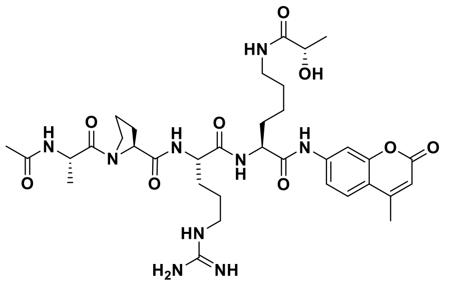


**Ac-Ala-Pro-Arg-Lys(D-La)-AMC (1c).** Compound **S5** (19 mg, 0.019 mmol) and sodium D-lactate (5 mg, 0.041 mmol) were dissolved in anh. DMF (1.0 mL) and cooled to 0 °C. $iPr_2NEt$ (14 µL, 0.082 mmol) and HATU (8 mg, 0.020 mmol) were added to the reaction mixture, which was stirred overnight going towards ambient temperature. Solvent was removed under reduced pressure to afford the crude intermediate tentatively assigned as Ac-Ala-Pro-Arg(Pbf)-Lys(D-la)-AMC (HPLC-MS $t_R$

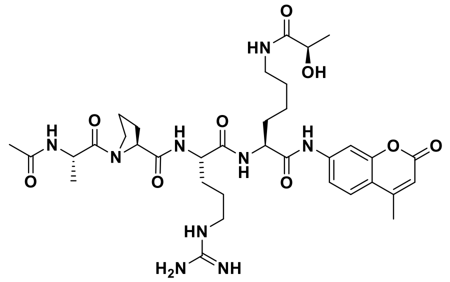

1.61 min, *m/z* 994.5 ([M+H]$^+$, $C_{48}H_{68}N_9O_{12}S^+$, Calcd 994.5), which was used without further purification. TFA/$H_2O$ (98:2, v/v, 1.0 mL) was added to the intermediate, which was stirred for 1 h at ambient temperature. Solvent was removed under a stream of nitrogen, and preparative reversed-phase HPLC purification afforded the title compound (9 mg, 63% from **S5**) as a white fluffy material after lyophilization. $^1$H NMR (600 MHz, DMSO-$d_6$) δ 10.32 (s, 1H, NH$_{AMC}$), 8.10 (d, $J$ = 7.3 Hz, 1H, NH$_{Ala}$), 8.05 (d, $J$ = 7.3 Hz, 1H, NH$_{\alpha,Lys}$), 8.00 (d, $J$ = 7.5 Hz, 1H, NH$_{\alpha,Arg}$), 7.80 (d, $J$ = 2.0 Hz, 1H, H8$_{AMC}$), 7.72 (d, $J$ = 8.7 Hz, 1H, H5$_{AMC}$), 7.67 (t, $J$ = 6.0 Hz, 1H, NH$_{\varepsilon,Lys}$), 7.53 (t, $J$ = 5.7 Hz, 1H, NH$_{\delta,Arg}$), 7.50 (dd, $J$ = 8.7, 2.0 Hz, 1H, H6$_{AMC}$), 6.27 (d, $J$ = 1.3 Hz, 1H, H3$_{AMC}$), 4.52 (p, $J$ = 7.0 Hz, 1H, H$_{\alpha,Ala}$), 4.40–4.29 (m, 2H, H$_{\alpha,Pro}$, H$_{\alpha,Lys}$), 4.26 (td, $J$ = 7.9, 5.4 Hz, 1H, H$_{\alpha,Arg}$), 3.92 (qd, $J$ = 6.8, 2.0 Hz, 1H, C(OH)$\underline{H}$CH$_3$), 3.66–3.60 (m, 1H, H$_{\delta,Pro,A}$), 3.58–3.53 (m, 1H, H$_{\delta,Pro,B}$), 3.11 (q, $J$ = 6.6 Hz, 2H, H$_{\delta,Arg}$), 3.08–2.99 (m, 2H, H$_{\varepsilon,Lys}$), 2.40 (d, $J$ = 1.3 Hz, 3H, CH$_{3,AMC}$), 2.10–1.98 (m, 1H, H$_{\beta,Pro,A}$), 1.95–1.21 (m, 16H, H$_{\beta,Pro,A}$, H$_{\gamma,Pro}$, H$_{\beta,Lys}$, H$_{\gamma,Lys}$, H$_{\delta,Lys}$, H$_{\beta,Arg}$, H$_{\gamma,Arg}$, CH$_{3,acetyl}$), 1.19–1.14 (m, 6H, C(OH)HC$\underline{H}_3$, H$_{\beta,Ala}$). $^{13}$C NMR (151 MHz, DMSO) δ 174.4 ($\underline{C}$ONH$_{\varepsilon,Lys}$), 171.8 (CO$_{\alpha,Pro}$), 171.5 (CO$_{Arg}$), 171.3 (CO$_{Ala}$), 171.2 (CO$_{\alpha,Lys}$), 168.8 ($\underline{C}$OCH$_{3,acetyl}$), 160.0 (C2$_{AMC}$), 158.2 (q, $J$ = 33.3 Hz, residual CO$_{TFA}$), 156.7 (NHC(=NH)NH$_2$), 153.6 (C8a$_{AMC}$), 153.1 (C4$_{AMC}$), 142.1 (C7$_{AMC}$), 125.9 (C5$_{AMC}$), 115.2 (C6$_{AMC}$), 115.1 (C4a$_{AMC}$), 112.4 (C3$_{AMC}$), 105.7 (C8$_{AMC}$), 67.2 ($\underline{C}$(OH)HCH$_3$), 59.6 (C$_{\alpha,Pro}$), 53.7 (C$_{\alpha,Lys}$), 52.2 (C$_{\alpha,Arg}$), 46.8 (C$_{\delta,Pro}$), 46.2 (C$_{\alpha,Ala}$), 40.4 (C$_{\delta,Arg}$), 37.9 (C$_{\varepsilon,Lys}$), 31.3 (C$_{\beta,Lys}$), 29.0 (C$_{\beta,Pro}$), 28.91 (C$_{\delta,Lys}$), 28.85 (C$_{\beta,Arg}$), 24.9 (C$_{\gamma,Arg}$), 24.5 (C$_{\gamma,Pro}$), 22.8 (C$_{\gamma,Lys}$), 22.2 (CO$\underline{C}$H$_{3,acetyl}$), 21.1 (C(OH)H$\underline{C}$H$_3$), 18.0 (CH$_{3,AMC}$), 16.9 (C$_{\beta,Ala}$). Two sets of signals (approximately 6:1) were detectable due to rotamers. Only peaks for the major rotamer is given. Analytical HPLC gradient 0–95% eluent II in eluent I (11 min total runtime), $t_R$ 4.19 min (>98%, UV$_{230}$). HRMS calcd for $C_{35}H_{52}N_9O_9^+$ [M+H]$^+$, 742.3882; found 742.3878. AMC=7-amino-4-methylcoumarin.

**Ac-Leu-Gly-Lys(L-La)-AMC (2b).** By the method described for **2c**, the title compound was synthesized using **S3** (27 mg, 0.032 mmol), L-lactate (11 mg, 0.127 mmol), *i*Pr$_2$NEt (44 µL, 0.253 mmol), and HATU (45 mg, 0.118 mmol). Preparative reversed-phase HPLC purification afforded the title compound (6 mg, 25% from **S3**) as a white fluffy material after lyophilization. $^1$H NMR (600 MHz, DMSO-$d_6$) δ 10.35 (s, 1H, NH$_{AMC}$), 8.30 (t, $J$ = 5.8 Hz, 1H, NH$_{Gly}$), 8.08 (d, $J$ = 7.4 Hz, 1H, NH$_{Leu}$), 8.01 (d, $J$ = 7.6 Hz, 1H, NH$_{\alpha,Lys}$), 7.79 (d, $J$ = 2.1 Hz, 1H, H8$_{AMC}$), 7.72 (d, $J$ = 8.7 Hz, 1H, H5$_{AMC}$), 7.64 (t, $J$ = 5.9 Hz, 1H, NH$_{\varepsilon,Lys}$), 7.52 (dd, $J$ = 8.7, 2.1 Hz, 1H, H6$_{AMC}$), 6.26 (d, $J$ = 1.3 Hz, 1H, H3$_{AMC}$), 4.37 (ddd, $J$ = 9.0, 7.6, 5.1 Hz, 1H, H$_{\alpha,Lys}$), 4.22 (ddd, $J$ = 9.1, 7.4, 6.0 Hz, 1H, H$_{\alpha,Leu}$), 3.92 (q, $J$ = 6.7 Hz, 1H, C(OH)$\underline{H}$CH$_3$), 3.79–3.67 (m, 2H, H$_{\alpha,Gly}$), 3.05 (q, $J$ = 6.8 Hz, 2H, H$_{\varepsilon,Lys}$), 2.40 (d, $J$ = 1.2 Hz, 3H, CH$_{3,AMC}$), 1.85 (s, 3H, CH$_{3,acetyl}$), 1.78–1.70 (m, 1H, H$_{\beta,Lys,A}$), 1.70–1.55 (m, 2H, H$_{\beta,Lys,A}$, H$_{\gamma,Leu}$), 1.50–1.37 (m, 4H, H$_{\beta,Leu}$, H$_{\delta,Lys}$), 1.37–1.29 (m, 1H, H$_{\gamma,Lys,A}$), 1.29–1.20 (m, 1H, H$_{\gamma,Lys,B}$), 1.17 (d, $J$ = 6.8 Hz, 3H, C(OH)HC$\underline{H}_3$), 0.88 (d, $J$ = 6.6 Hz, 3H, H$_{\delta,Leu,1}$), 0.84 (d, $J$ = 6.6 Hz, 3H, H$_{\delta,Leu,2}$). $^{13}$C NMR (151 MHz, DMSO) δ 174.3 ($\underline{C}$ONH$_{\varepsilon,Lys}$), 172.9 (CO$_{Leu}$), 171.4 (CO$_{\alpha,Lys}$), 169.7 ($\underline{C}$OCH$_{3,acetyl}$), 169.0 (CO$_{Gly}$), 160.0 (C2$_{AMC}$), 153.6 (C8a$_{AMC}$), 153.1 (C4$_{AMC}$), 142.1 (C7$_{AMC}$), 125.9 (C5$_{AMC}$), 115.3 (C6$_{AMC}$),

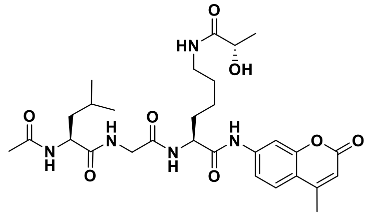

115.1 (C4a$_{AMC}$), 112.3 (C3$_{AMC}$), 105.7 (C8$_{AMC}$), 67.2 ($\underline{C}$(OH)HCH$_3$), 53.6 ($C_{\alpha,Lys}$), 51.4 ($C_{\alpha,Leu}$), 42.0 ($C_{\alpha,Gly}$), 40.5 ($C_{\beta,Leu}$), 37.9 ($C_{\varepsilon,Lys}$), 31.4 ($C_{\beta,Lys}$), 28.9 ($C_{\delta,Lys}$), 24.2 ($C_{\gamma,Leu}$), 22.9 ($C_{\delta,Leu,1}$), 22.8 ($C_{\gamma,Lys}$), 22.5 (CO$\underline{C}$H$_{3,acetyl}$), 21.6 ($C_{\delta,Leu,2}$), 21.1 (C(OH)H$\underline{C}$H$_3$), 18.0 (CH$_{3,AMC}$). Analytical HPLC gradient 0–95% eluent II in eluent I (11 min total runtime), $t_R$ 4.97 min (>98%, UV$_{230}$). HRMS calcd for $C_{29}H_{41}N_5NaO_8^+$ [M+Na]$^+$, 610.2847; found 610.2842. AMC=7-amino-4-methylcoumarin.

**Ac-Leu-Gly-Lys(D-La)-AMC (2c).** Compound **S3** (26 mg, 0.041 mmol) and D-lactate (11 mg, 0.124 mmol) were dissolved in anh. DMF (1.5 mL) and cooled to 0 °C. *i*Pr$_2$NEt (43 µL, 0.249 mmol) and HATU (44 mg, 0.116 mmol) were added to the reaction mixture, which was stirred for 3 h going towards ambient temperature. Solvent was removed under reduced pressure, diluted with $CH_2Cl_2$ (20 mL) and washed with aq. HCl (0.1 M, 4×10 mL), sat. $NaHCO_3$ (10 mL) and dried over $MgSO_4$. Solvent was removed under reduced pressure, and preparative reversed-phase HPLC purification afforded the title compound (8 mg, 33% from **S3**) as a white fluffy material after lyophilization. $^1$H NMR (600 MHz, DMSO-$d_6$) δ 10.35 (s, 1H, NH$_{AMC}$), 8.31 (t, *J* = 5.9 Hz, 1H, NH$_{Gly}$), 8.08 (d, *J* = 7.3 Hz, 1H, NH$_{Leu}$), 8.01 (d, *J* = 7.6 Hz, 1H, NH$_{\alpha,Lys}$), 7.79 (d, *J* = 2.0 Hz, 1H, H8$_{AMC}$), 7.72 (d, *J* = 8.7 Hz, 1H, H5$_{AMC}$), 7.64 (t, *J* = 5.9 Hz, 1H, NH$_{\varepsilon,Lys}$), 7.53 (dd, *J* = 8.7, 2.1 Hz, 1H, H6$_{AMC}$), 6.26 (d, *J* = 1.3 Hz, 1H, H3$_{AMC}$), 4.37 (ddd, *J* = 9.1, 7.6, 5.1 Hz, 1H, $H_{\alpha,Lys}$), 4.22 (ddd, *J* = 9.0, 7.3, 6.0 Hz, 1H, $H_{\alpha,Leu}$), 3.92 (q, *J* = 6.7 Hz, 1H, C(OH)$\underline{H}$CH$_3$), 3.79–3.67 (m, 2H, $H_{\alpha,Gly}$), 3.05 (q, *J* = 6.8 Hz, 2H, $H_{\varepsilon,Lys}$), 2.40 (d, *J* = 1.3 Hz, 3H, CH$_{3,AMC}$), 1.85 (s, 3H, CH$_{3,acetyl}$), 1.78–1.70 (m, 1H, $H_{\beta,Lys,A}$), 1.69–1.55 (m, 2H, $H_{\beta,Lys,A}$, $H_{\gamma,Leu}$), 1.50–1.38 (m, 4H, $H_{\beta,Leu}$, $H_{\delta,Lys}$), 1.37–1.29 (m, 1H, $H_{\gamma,Lys,A}$), 1.29–1.20 (m, 1H, $H_{\gamma,Lys,B}$), 1.17 (d, *J* = 6.8 Hz, 3H, C(OH)HC$\underline{H}_3$), 0.88 (d, *J* = 6.6 Hz, 3H, $H_{\delta,Leu,1}$), 0.84 (d, *J* = 6.6 Hz, 3H, $H_{\delta,Leu,2}$). $^{13}$C NMR (151 MHz, DMSO) δ 174.3 ($\underline{C}$ONH$_{\varepsilon,Lys}$), 172.9 (CO$_{Leu}$), 171.4 (CO$_{\alpha,Lys}$), 169.7 ($\underline{C}$OCH$_{3,acetyl}$), 169.0 (CO$_{Gly}$), 160.0 (C2$_{AMC}$), 153.6 (C8a$_{AMC}$), 153.1 (C4$_{AMC}$), 142.1 (C7$_{AMC}$), 125.9 (C5$_{AMC}$), 115.3 (C6$_{AMC}$), 115.1 (C4a$_{AMC}$), 112.3 (C3$_{AMC}$), 105.7 (C8$_{AMC}$), 67.2 ($\underline{C}$(OH)HCH$_3$), 53.6 ($C_{\alpha,Lys}$), 51.4 ($C_{\alpha,Leu}$), 42.0 ($C_{\alpha,Gly}$), 40.5 ($C_{\beta,Leu}$), 37.9 ($C_{\varepsilon,Lys}$), 31.4 ($C_{\beta,Lys}$), 28.9 ($C_{\delta,Lys}$), 24.2 ($C_{\gamma,Leu}$), 22.9 ($C_{\delta,Leu,1}$), 22.8 ($C_{\gamma,Lys}$), 22.5 (CO$\underline{C}$H$_{3,acetyl}$), 21.6 ($C_{\delta,Leu,2}$), 21.1 (C(OH)H$\underline{C}$H$_3$), 18.0 (CH$_{3,AMC}$). Analytical HPLC gradient 0–95% eluent II in eluent I (11 min total runtime), $t_R$ 4.98 min (>98%, UV$_{230}$). HRMS calcd for $C_{29}H_{41}N_5NaO_8^+$ [M+Na]$^+$, 610.2847; found 610.2842. AMC=7-amino-4-methylcoumarin.

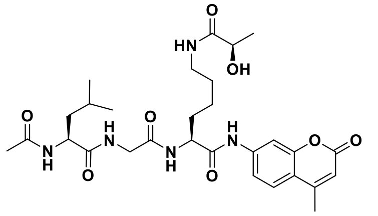

**Ac-Leu-Gly-Lys(For)-AMC (2l).** Compound **S3** (12 mg, 0.019 mmol) was dissolved in ethyl formate (0.77 mL, 9.5 mmol), and *i*Pr$_2$NEt (20 µL, 0.114 mmol) was added to the reaction mixture, which was stirred overnight at reflux. Solvent was removed under reduced pressure, and preparative reversed-phase HPLC purification afforded the title compound (4 mg, 40% from **S3**) as a white fluffy material after lyophilization. $^1$H NMR (600 MHz, DMSO-$d_6$) δ 10.36 (s, 1H, NH$_{AMC}$), 8.32 (t, *J* = 5.9 Hz, 1H, NH$_{Gly}$), 8.09 (d, *J* = 7.3 Hz, 1H, NH$_{Leu}$), 8.00 (d, *J* = 7.7 Hz, 1H, NH$_{\alpha,Lys}$), 7.98 (s, 1H, C$\underline{H}$O), 7.98–7.94 (m, 1H, NH$_{\varepsilon,Lys}$), 7.79 (d, *J* = 2.1 Hz, 1H, H8$_{AMC}$), 7.72 (d, *J* = 8.6 Hz, 1H, H5$_{AMC}$), 7.52 (dd, *J* = 8.7, 2.1 Hz, 1H, H6$_{AMC}$), 6.26 (d, *J* = 1.3 Hz, 1H, H3$_{AMC}$), 4.38 (td, *J* = 8.1, 5.0 Hz, 1H, $H_{\alpha,Lys}$), 4.22 (ddd, *J* = 9.1, 7.3, 6.0 Hz, 1H, $H_{\alpha,Leu}$), 3.78–3.68 (m, 2H, $H_{\alpha,Gly}$), 3.06 (q, *J* = 6.7 Hz, 2H, $H_{\varepsilon,Lys}$), 2.40 (d, *J* = 1.3 Hz, 3H, CH$_{3,AMC}$),

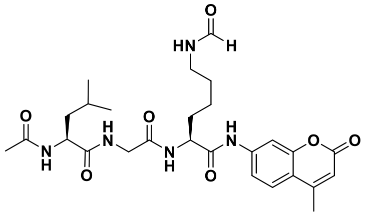

1.85 (s, 3H, $CH_{3,acetyl}$), 1.74 (ddt, $J$ = 14.5, 10.4, 5.3 Hz, 1H, $H_{β,Lys,A}$), 1.69–1.56 (m, 2H, $H_{β,Lys,A}$, $H_{γ,Leu}$), 1.50–1.39 (m, 4H, $H_{β,Leu}$, $H_{δ,Lys}$), 1.39–1.31 (m, 1H, $H_{γ,Lys,A}$), 1.31–1.21 (m, 1H, $H_{γ,Lys,B}$), 0.88 (d, $J$ = 6.6 Hz, 3H, $H_{δ,Leu,1}$), 0.84 (d, $J$ = 6.5 Hz, 3H, $H_{δ,Leu,2}$). $^{13}$C NMR (151 MHz, DMSO) δ 172.9 ($CO_{Leu}$), 171.3 ($CO_{α,Lys}$), 169.7 ($\underline{C}OCH_{3,acetyl}$), 169.0 ($CO_{Gly}$), 160.9 ($\underline{C}HO$), 160.0 ($C2_{AMC}$), 153.6 ($C8a_{AMC}$), 153.1 ($C4_{AMC}$), 142.1 ($C7_{AMC}$), 125.9 ($C5_{AMC}$), 115.3 ($C6_{AMC}$), 115.1 ($C4a_{AMC}$), 112.4 ($C3_{AMC}$), 105.8 ($C8_{AMC}$), 53.5 ($C_{α,Lys}$), 51.5 ($C_{α,Leu}$), 42.1 ($C_{α,Gly}$), 40.5 ($C_{β,Leu}$), 36.9 ($C_{ε,Lys}$), 31.3 ($C_{β,Lys}$), 28.7 ($C_{δ,Lys}$), 24.2 ($C_{γ,Leu}$), 22.9 ($C_{δ,Leu,1}$), 22.8 ($C_{γ,Lys}$), 22.5 ($CO\underline{C}H_{3,acetyl}$), 21.6 ($C_{δ,Leu,2}$), 18.0 ($CH_{3,AMC}$). Analytical HPLC gradient 0–95% eluent II in eluent I (11 min total runtime), $t_R$ 4.96 min (>98%, $UV_{215}$). HRMS calcd for $C_{27}H_{37}N_5NaO_7^+$ [M+Na]$^+$, 566.2585; found 566.2580. AMC=7-amino-4-methylcoumarin.

**Ac-Leu-Gly-Lys(Gc)-AMC (2n).** Compound **S3** (12 mg, 0.019 mmol) and glycolic acid (6 mg, 0.078 mmol) were dissolved in anh. DMF (1.0 mL) and cooled to 0 °C. $i$Pr$_2$NEt (20 µL, 0.114 mmol) and HATU (21 mg, 0.055 mmol) were added to the reaction mixture, which was stirred for 3.5 h going towards ambient temperature. Solvent was removed under reduced pressure, and preparative reversed-phase HPLC purification afforded the title compound (4 mg, 39% from **S3**) as a white fluffy material after lyophilization. $^1$H NMR (600 MHz, DMSO-$d_6$) δ 10.35 (s, 1H, $NH_{AMC}$), 8.31 (t, $J$ = 5.9 Hz, 1H, $NH_{Gly}$), 8.08 (d, $J$ = 7.3 Hz, 1H, $NH_{Leu}$), 8.01 (d, $J$ = 7.6 Hz, 1H, $NH_{α,Lys}$), 7.79 (d, $J$ = 2.0 Hz, 1H, $H8_{AMC}$), 7.72 (d, $J$ = 8.7 Hz, 1H, $H5_{AMC}$), 7.69 (t, $J$ = 6.0 Hz, 1H, $NH_{ε,Lys}$), 7.52 (dd, $J$ = 8.7, 2.0 Hz, 1H, $H6_{AMC}$), 6.26 (d, $J$ = 1.3 Hz, 1H, $H3_{AMC}$), 4.37 (ddd, $J$ = 9.1, 7.5, 5.0 Hz, 1H, $H_{α,Lys}$), 4.22 (ddd, $J$ = 8.9, 7.4, 6.0 Hz, 1H, $H_{α,Leu}$), 3.77 (s, 2H, $C\underline{H}_2OH$), 3.76–3.68 (m, 2H, $H_{α,Gly}$), 3.08 (q, $J$ = 6.8 Hz, 2H, $H_{ε,Lys}$), 2.40 (d, $J$ = 1.3 Hz, 3H, $CH_{3,AMC}$), 1.85 (s, 3H, $CH_{3,acetyl}$), 1.78–1.70 (m, 1H, $H_{β,Lys,A}$), 1.70–1.56 (m, 2H, $H_{β,Lys,A}$, $H_{γ,Leu}$), 1.50–1.39 (m, 4H, $H_{β,Leu}$, $H_{δ,Lys}$), 1.38–1.30 (m, 1H, $H_{γ,Lys,A}$), 1.30–1.21 (m, 1H, $H_{γ,Lys,B}$), 0.88 (d, $J$ = 6.6 Hz, 3H, $H_{δ,Leu,1}$), 0.84 (d, $J$ = 6.6 Hz, 3H, $H_{δ,Leu,2}$). $^{13}$C NMR (151 MHz, DMSO) δ 172.9 ($CO_{Leu}$), 171.6 ($\underline{C}ONH_{ε,Lys}$), 171.4 ($CO_{α,Lys}$), 169.7 ($\underline{C}OCH_{3,acetyl}$), 169.0 ($CO_{Gly}$), 160.0 ($C2_{AMC}$), 158.2 (q, $J$ = 36.8 Hz, residual $CO_{TFA}$), 153.6 ($C8a_{AMC}$), 153.1 ($C4_{AMC}$), 142.1 ($C7_{AMC}$), 125.9 ($C5_{AMC}$), 115.3 ($C6_{AMC}$), 115.1 ($C4a_{AMC}$), 112.3 ($C3_{AMC}$), 105.8 ($C8_{AMC}$), 61.4 ($\underline{C}H_2OH$), 53.6 ($C_{α,Lys}$), 51.5 ($C_{α,Leu}$), 42.0 ($C_{α,Gly}$), 40.5 ($C_{β,Leu}$), 37.8 ($C_{ε,Lys}$), 31.4 ($C_{β,Lys}$), 29.0 ($C_{δ,Lys}$), 24.2 ($C_{γ,Leu}$), 22.9 ($C_{δ,Leu,1}$), 22.8 ($C_{γ,Lys}$), 22.5 ($CO\underline{C}H_{3,acetyl}$), 21.6 ($C_{δ,Leu,2}$), 18.0 ($CH_{3,AMC}$). Analytical HPLC gradient 0–95% eluent II in eluent I (11 min total runtime), $t_R$ 4.87 min (>99%, $UV_{215}$). HRMS calcd for $C_{28}H_{39}N_5NaO_8^+$ [M+Na]$^+$, 596.2690; found 596.2682. AMC=7-amino-4-methylcoumarin.

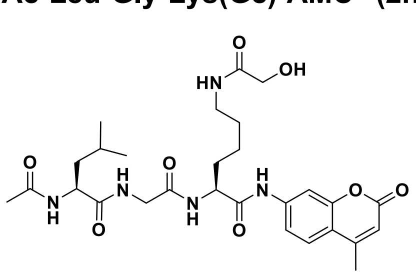

**Ac-Leu-Gly-Lys(Hib)-AMC (2o).** By the method described for **2n**, the title compound was synthesized using **S3** (12 mg, 0.032 mmol), 2-hydroxyisobutyric acid (7 mg, 0.068 mmol), $i$Pr$_2$NEt (20 µL, 0.114 mmol), and HATU (20 mg, 0.053 mmol). Preparative reversed-phase HPLC purification afforded the title compound (7 mg, 60% from **S3**) as a white fluffy material after lyophilization. $^1$H NMR (600 MHz, DMSO-$d_6$) δ 10.36 (s, 1H, $NH_{AMC}$), 8.29 (t, $J$ = 5.9 Hz, 1H, $NH_{Gly}$), 8.07 (d, $J$ = 7.4 Hz, 1H, $NH_{Leu}$), 8.02 (d, $J$ = 7.5 Hz, 1H, $NH_{α,Lys}$), 7.79 (d, $J$ = 2.0 Hz, 1H, $H8_{AMC}$), 7.72 (d, $J$ =

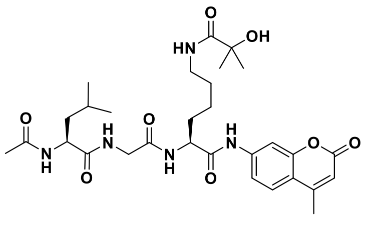

8.7 Hz, 1H, H5$_{AMC}$), 7.60 (t, $J$ = 6.0 Hz, 1H, NH$_{\varepsilon,Lys}$), 7.52 (dd, $J$ = 8.7, 2.0 Hz, 1H, H6$_{AMC}$), 6.26 (d, $J$ = 1.4 Hz, 1H, H3$_{AMC}$), 4.37 (td, $J$ = 8.2, 5.1 Hz, 1H, H$_{\alpha,Lys}$), 4.22 (ddd, $J$ = 9.2, 7.4, 5.9 Hz, 1H, H$_{\alpha,Leu}$), 3.78–3.67 (m, 2H, H$_{\alpha,Gly}$), 3.04 (q, $J$ = 6.8 Hz, 2H, H$_{\varepsilon,Lys}$), 2.40 (d, $J$ = 1.4 Hz, 3H, CH$_{3,AMC}$), 1.85 (s, 3H, CH$_{3,acetyl}$), 1.74 (ddt, $J$ = 15.3, 10.7, 5.4 Hz, 1H, H$_{\beta,Lys,A}$), 1.69–1.56 (m, 2H, H$_{\beta,Lys,A}$, H$_{\gamma,Leu}$), 1.49–1.39 (m, 4H, H$_{\beta,Leu}$, H$_{\delta,Lys}$), 1.36–1.29 (m, 1H, H$_{\gamma,Lys,A}$), 1.29–1.22 (m, 1H, H$_{\gamma,Lys,B}$), 1.21 (s, 3H, CH$_{3,Hib,A}$), 1.20 (s, 3H, CH$_{3,Hib,B}$), 0.88 (d, $J$ = 6.6 Hz, 3H, H$_{\delta,Leu,1}$), 0.84 (d, $J$ = 6.6 Hz, 3H, H$_{\delta,Leu,2}$). $^{13}$C NMR (151 MHz, DMSO) δ 176.2 ($\underline{C}$ONH$_{\varepsilon,Lys}$), 172.9 (CO$_{Leu}$), 171.4 (CO$_{\alpha,Lys}$), 169.6 ($\underline{C}$OCH$_{3,acetyl}$), 169.0 (CO$_{Gly}$), 160.0 (C2$_{AMC}$), 153.6 (C8a$_{AMC}$), 153.1 (C4$_{AMC}$), 142.1 (C7$_{AMC}$), 125.9 (C5$_{AMC}$), 115.3 (C6$_{AMC}$), 115.1 (C4a$_{AMC}$), 112.3 (C3$_{AMC}$), 105.7 (C8$_{AMC}$), 71.8 ($\underline{C}$OH), 53.6 (C$_{\alpha,Lys}$), 51.4 (C$_{\alpha,Leu}$), 42.0 (C$_{\alpha,Gly}$), 40.5 (C$_{\beta,Leu}$), 38.0 (C$_{\varepsilon,Lys}$), 31.4 (C$_{\beta,Lys}$), 28.9 (C$_{\delta,Lys}$), 27.8 ($\underline{C}$H$_{3,Hib,A}$, $\underline{C}$H$_{3,Hib,B}$), 24.2 (C$_{\gamma,Leu}$), 22.9 (C$_{\delta,Leu,1}$), 22.7 (C$_{\gamma,Lys}$), 22.5 (CO$\underline{C}$H$_{3,acetyl}$), 21.6 (C$_{\delta,Leu,2}$), 18.0 (CH$_{3,AMC}$). Analytical HPLC gradient 0–95% eluent II in eluent I (11 min total runtime), $t_R$ 5.13 min (>97%, UV$_{215}$). HRMS calcd for $C_{30}H_{43}N_5NaO_8^+$ [M+Na]$^+$, 624.3004; found 624.2999. AMC=7-amino-4-methylcoumarin.

**Ac-Leu-Gly-Lys(L-Bhb)-AMC (2p).** By the method described for **2n**, the title compound was synthesized using **S3** (12 mg, 0.032 mmol), L-β-hydroxybutyric acid (6 mg, 0.061 mmol), *i*Pr$_2$NEt (20 µL, 0.114 mmol), and HATU (20 mg, 0.053 mmol). Preparative reversed-phase HPLC purification afforded the title compound (3 mg, 23% from **S3**) as a white fluffy material after lyophilization. $^1$H NMR (600 MHz, DMSO-$d_6$) δ 10.36 (s, 1H, NH$_{AMC}$), 8.31 (t, $J$ = 6.0 Hz, 1H, NH$_{Gly}$), 8.08 (d, $J$ = 7.2 Hz, 1H, NH$_{Leu}$), 8.00 (d, $J$ = 7.7 Hz, 1H, NH$_{\alpha,Lys}$), 7.80 (d, $J$ = 2.3 Hz, 1H, H8$_{AMC}$), 7.76 (t, $J$ = 5.6 Hz, 1H, NH$_{\varepsilon,Lys}$), 7.72 (d, $J$ = 8.7 Hz, 1H, H5$_{AMC}$), 7.53 (dd, $J$ = 8.8, 2.4 Hz, 1H, H6$_{AMC}$), 6.26 (s, 1H, H3$_{AMC}$), 4.41–4.33 (m, 1H, H$_{\alpha,Lys}$), 4.22 (dt, $J$ = 7.3, 6.6 Hz, 1H, H$_{\alpha,Leu}$), 3.95–3.89 (m, 1H, C$\underline{H}$OH), 3.78–3.66 (m, 2H, H$_{\alpha,Gly}$), 3.08–2.93 (m, 2H, H$_{\varepsilon,Lys}$), 2.40 (d, $J$ = 2.6 Hz, 3H, CH$_{3,AMC}$), 2.17 (dd, $J$ = 14.0, 7.2 Hz, 1H, C$\underline{H}_{2,A}$CHOH), 2.06 (dd, $J$ = 13.8, 6.0 Hz, 1H, C$\underline{H}_{2,B}$CHOH), 1.85 (s, 3H, CH$_{3,acetyl}$), 1.78–1.70 (m, 1H, H$_{\beta,Lys,A}$), 1.69–1.54 (m, 2H, H$_{\beta,Lys,A}$, H$_{\gamma,Leu}$), 1.50–1.43 (m, 2H, H$_{\beta,Leu}$), 1.43–1.35 (m, 2H, H$_{\delta,Lys}$), 1.35–1.20 (m, 2H, H$_{\gamma,Lys}$), 1.02 (d, $J$ = 6.2 Hz, 3H, CH(OH)C$\underline{H}_3$), 0.88 (d, $J$ = 6.6 Hz, 3H, H$_{\delta,Leu,1}$), 0.84 (d, $J$ = 6.6 Hz, 3H, H$_{\delta,Leu,2}$). $^{13}$C NMR (151 MHz, DMSO) δ 172.9 (CO$_{Leu}$), 171.4 (CO$_{\alpha,Lys}$), 170.5 ($\underline{C}$ONH$_{\varepsilon,Lys}$), 169.7 ($\underline{C}$OCH$_{3,acetyl}$), 169.0 (CO$_{Gly}$), 160.0 (C2$_{AMC}$), 153.6 (C8a$_{AMC}$), 153.1 (C4$_{AMC}$), 142.1 (C7$_{AMC}$), 125.9 (C5$_{AMC}$), 115.3 (C6$_{AMC}$), 115.1 (C4a$_{AMC}$), 112.3 (C3$_{AMC}$), 105.8 (C8$_{AMC}$), 63.8 ($\underline{C}$HOH), 53.6 (C$_{\alpha,Lys}$), 51.5 (C$_{\alpha,Leu}$), 45.3 ($\underline{C}$H$_2$CH), 42.0 (C$_{\alpha,Gly}$), 40.5 (C$_{\beta,Leu}$), 38.1 (C$_{\varepsilon,Lys}$), 31.4 (C$_{\beta,Lys}$), 28.8 (C$_{\delta,Lys}$), 24.2 (C$_{\gamma,Leu}$), 23.3 (CH(OH)$\underline{C}$H$_3$), 22.9 (C$_{\delta,Leu,1}$), 22.8 (C$_{\gamma,Lys}$), 22.5 (CO$\underline{C}$H$_{3,acetyl}$), 21.6 (C$_{\delta,Leu,2}$), 18.0 (CH$_{3,AMC}$). Analytical HPLC gradient 0–95% eluent II in eluent I (11 min total runtime), $t_R$ 4.97 min (>95%, UV$_{215}$). HRMS calcd for $C_{30}H_{43}N_5NaO_8^+$ [M+Na]$^+$, 624.3004; found 624.2999. AMC=7-amino-4-methylcoumarin.

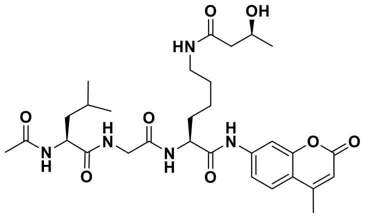

**Ac-Leu-Gly-Lys(D-Bhb)-AMC (2q).** By the method described for **2n**, the title compound was synthesized using **S3** (12 mg, 0.032 mmol), D-β-hydroxybutyric acid (7 mg, 0.066 mmol), *i*Pr$_2$NEt (20 µL, 0.114 mmol), and HATU (20 mg, 0.053 mmol). Preparative reversed-phase HPLC purification afforded the title compound (5 mg, 45% from **S3**) as a white fluffy material after lyophilization. $^1$H NMR (600 MHz, DMSO-$d_6$) δ 10.36 (s, 1H, NH$_{AMC}$), 8.32 (t, $J$ = 5.9 Hz, 1H, NH$_{Gly}$), 8.08 (d, $J$ = 7.3 Hz, 1H,

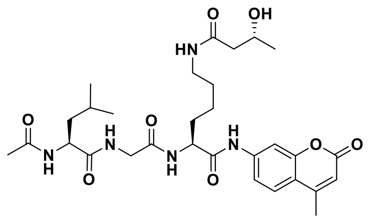

$NH_{Leu}$), 7.99 (d, $J$ = 7.7 Hz, 1H, $NH_{\alpha,Lys}$), 7.79 (d, $J$ = 2.0 Hz, 1H, $H8_{AMC}$), 7.75 (t, $J$ = 5.6 Hz, 1H, $NH_{\varepsilon,Lys}$), 7.72 (d, $J$ = 8.7 Hz, 1H, $H5_{AMC}$), 7.53 (dd, $J$ = 8.7, 2.1 Hz, 1H, $H6_{AMC}$), 6.26 (d, $J$ = 1.3 Hz, 1H, $H3_{AMC}$), 4.37 (td, $J$ = 8.2, 5.1 Hz, 1H, $H_{\alpha,Lys}$), 4.24–4.19 (m, 1H, $H_{\alpha,Leu}$), 3.93 (dt, $J$ = 7.1, 6.0 Hz, 1H, C<u>H</u>OH), 3.78–3.66 (m, 2H, $H_{\alpha,Gly}$), 3.08–2.95 (m, 2H, $H_{\varepsilon,Lys}$), 2.40 (d, $J$ = 1.3 Hz, 3H, $CH_{3,AMC}$), 2.17 (dd, $J$ = 13.8, 7.2 Hz, 1H, $C\underline{H}_{2,A}$CHOH), 2.06 (dd, $J$ = 13.8, 5.9 Hz, 1H, $C\underline{H}_{2,B}$CHOH), 1.85 (s, 3H, $CH_{3,acetyl}$), 1.78–1.70 (m, 1H, $H_{\beta,Lys,A}$), 1.69–1.56 (m, 2H, $H_{\beta,Lys,A}$, $H_{\gamma,Leu}$), 1.49–1.43 (m, 2H, $H_{\beta,Leu}$), 1.43–1.36 (m, 2H, $H_{\delta,Lys}$), 1.36–1.29 (m, 1H, $H_{\gamma,Lys,A}$), 1.29–1.22 (m, 1H, $H_{\gamma,Lys,B}$), 1.03 (d, $J$ = 6.2 Hz, 3H, $CH(OH)C\underline{H}_3$), 0.88 (d, $J$ = 6.6 Hz, 3H, $H_{\delta,Leu,1}$), 0.84 (d, $J$ = 6.6 Hz, 3H, $H_{\delta,Leu,2}$). $^{13}$C NMR (151 MHz, DMSO) δ 172.9 ($CO_{Leu}$), 171.4 ($CO_{\alpha,Lys}$), 170.5 ($\underline{C}ONH_{\varepsilon,Lys}$), 169.7 ($\underline{C}OCH_{3,acetyl}$), 169.0 ($CO_{Gly}$), 160.0 ($C2_{AMC}$), 158.2 (q, $J$ = 36.5 Hz, residual $CO_{TFA}$), 153.6 ($C8a_{AMC}$), 153.1 ($C4_{AMC}$), 142.1 ($C7_{AMC}$), 125.9 ($C5_{AMC}$), 115.3 ($C6_{AMC}$), 115.1 ($C4a_{AMC}$), 112.3 ($C3_{AMC}$), 105.8 ($C8_{AMC}$), 63.8 (<u>C</u>HOH), 53.6 ($C_{\alpha,Lys}$), 51.5 ($C_{\alpha,Leu}$), 45.3 ($\underline{C}H_2CH$), 42.0 ($C_{\alpha,Gly}$), 40.5 ($C_{\beta,Leu}$), 38.1 ($C_{\varepsilon,Lys}$), 31.4 ($C_{\beta,Lys}$), 28.8 ($C_{\delta,Lys}$), 24.2 ($C_{\gamma,Leu}$), 23.3 ($CH(OH)\underline{C}H_3$), 22.9 ($C_{\delta,Leu,1}$), 22.8 ($C_{\gamma,Lys}$), 22.5 ($CO\underline{C}H_{3,acetyl}$), 21.6 ($C_{\delta,Leu,2}$), 18.0 ($CH_{3,AMC}$). Analytical HPLC gradient 0–95% eluent II in eluent I (11 min total runtime), $t_R$ 4.98 min (>97%, $UV_{215}$). HRMS calcd for $C_{30}H_{43}N_5NaO_8^+$ $[M+Na]^+$, 624.3004; found 624.3004. AMC=7-amino-4-methylcoumarin.

**Ac-Gln-Pro-Lys-Lys(L-La)-AMC (4b).** L-lactate (5 mg, 0.053 mmol) was dissolved in anh. DMF/$CH_2Cl_2$ (1:2, v/v, 3.0 mL) and cooled to 0 °C. HATU (20 mg, 0.053 mmol), lutidine (12 µL, 0.106 mmol) and compound **S2** (50 mg, 0.048 mmol) were added to the reaction mixture, which was stirred overnight going towards ambient temperature. The solution was diluted with $CH_2Cl_2$ (25 mL), and washed with brine (30 mL), aq. HCl (0.5 M, 3×30 mL), sat. $NaHCO_3$ (3×30 mL), brine (30 mL) and dried over $MgSO_4$, filtered and concentrated under reduced pressure to afford the crude intermediate tentatively assigned as Ac-Gln(Trt)-Pro-Lys(Boc)-Lys(L-la)-AMC (HPLC-MS $t_R$ 1.94 min, $m/z$ 1111.9 ($[M-H]^-$, $C_{61}H_{75}N_8O_{12}^-$, Calcd 1111.6), which was used without further purification. TFA/$CH_2Cl_2$/$H_2O$/TIPS (47.5:47.5:2.5:2.5, v/v, 4.0 mL) was added to the intermediate and stirred for 1 h at ambient temperature. Solvent was removed under a stream of nitrogen, after which the crude was triturated in ice-cold ether followed by preparative reversed-phase HPLC purification to afford the title compound (19 mg, 44% from **S2**) as a white fluffy material after lyophilization.

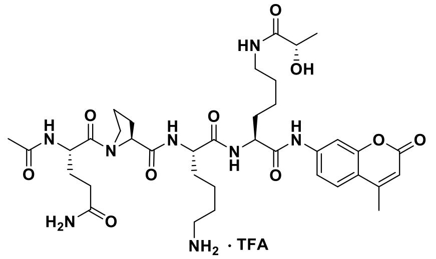

$^1$H NMR (600 MHz, DMSO-$d_6$) δ10.36 (s, 1H, $NH_{AMC}$), 8.11–8.02 (m, 3H, $NH_{\alpha,Gln}$, $NH_{\alpha,Lys}$ $NH_{\alpha,Lys(lactoyl)}$), 7.80 (d, $J$ = 2.0 Hz, 1H, $H8_{AMC}$), 7.76–7.70 (m, 4H, $H5_{AMC}$, $NH_{\varepsilon,Lys}$), 7.67 (t, $J$ = 6.0 Hz, 1H, $NH_{\varepsilon,Lys(lactoyl)}$), 7.48 (dd, $J$ = 8.7, 2.0 Hz, 1H, $H6_{AMC}$), 7.34 (s, 1H, $CONH_{2,Gln,A}$), 6.82 (s, 1H, $CONH_{2,Gln,B}$), 6.27 (d, $J$ = 1.2 Hz, 1H, $H3_{AMC}$), 4.46 (q, $J$ = 8.0 Hz, 1H, $H_{\alpha,Gln}$), 4.38–4.30 (m, 2H, $H_{\alpha,Pro}$, $H_{\alpha,Lys(lactoyl)}$), 4.30–4.23 (m, 1H, $H_{\alpha,Lys}$), 3.92 (q, $J$ = 6.8 Hz, 1H, $C(OH)\underline{H}CH_3$), 3.71–3.59 (m, 2H, $H_{\delta,Pro}$), 3.11–3.00 (m, 2H, $H_{\varepsilon,Lys(lactoyl)}$), 2.83–2.72 (m, 2H, $H_{\varepsilon,Lys}$), 2.40 (d, $J$ = 1.1 Hz, 3H, $CH_{3,AMC}$), 2.19–2.09 (m, 2H, $H_{\delta,Gln}$), 2.09–2.00 (m, 1H, $H_{\beta,Pro,A}$), 1.94–1.21 (m, XH, $H_{\beta,Lys(lactoyl)}$, $H_{\gamma,Lys(lactoyl)}$, $H_{\delta,Lys(lactoyl)}$, $H_{\beta,Lys}$, $H_{\gamma,Lys}$, $H_{\delta,Lys}$, $H_{\beta,Pro,B}$, $H_{\gamma,Pro}$, $H_{\beta,Gln}$, $CH_{3,acetyl}$), 1.16 (d, $J$ = 6.8 Hz, 3H, $C(OH)HC\underline{H}_3$). $^{13}$C NMR (151 MHz, DMSO) δ 174.4 ($\underline{C}ONH_{\varepsilon,Lys(lactoyl)}$), 173.9 ($CO_{\delta,Gln}$), 171.9 ($CO_{\alpha,Lys}$), 171.7 ($CO_{Pro}$), 171.4 ($CO_{\alpha,Lys(lactoyl)}$), 170.5 ($CO_{\alpha,Gln}$),

169.2 ($\underline{C}OCH_{3,acetyl}$), 160.0 (C2$_{AMC}$), 158.3 (q, $J$ = 34.4 Hz, residual CO$_{TFA}$), 153.7 (C8a$_{AMC}$), 153.1 (C4$_{AMC}$), 142.1 (C7$_{AMC}$), 126.0 (C5$_{AMC}$), 116.2 (q, $J$ = 295.4 Hz, CF$_{3,TFA}$), 115.2 (C6$_{AMC}$), 115.1 (C4a$_{AMC}$), 112.3 (C3$_{AMC}$), 105.7 (C8$_{AMC}$), 67.3 ($\underline{C}$(OH)HCH$_3$), 59.5 (C$_{\alpha,Pro}$), 53.7 (C$_{\alpha,Lys(lactoyl)}$), 52.3 (C$_{\alpha,Lys}$), 49.9 (C$_{\alpha,Gln}$), 46.9 (C$_{\delta,Pro}$), 38.8 (C$_{\varepsilon,Lys}$), 37.9 (C$_{\varepsilon,Lys(lactoyl)}$), 31.4 (C$_{\gamma,Gln}$), 31.1 (C$_{\beta,Lys}$), 31.0 (C$_{\beta,Lys(lactoyl)}$), 29.1 (C$_{\beta,Pro}$), 29.0 (C$_{\delta,Lys(lactoyl)}$), 27.2 (C$_{\beta,Gln}$), 26.5 (C$_{\delta,Lys}$), 24.5 (C$_{\gamma,Pro}$), 22.9 (C$_{\gamma,Lys(lactoyl)}$), 22.3 (C$_{\gamma,Lys}$), 22.1 (CH$_{3,acetyl}$), 21.1 (C(OH)HCH$_3$), 18.0 (CH$_{3,AMC}$). Two sets of signals (approximately 9:1) were detectable due to rotamers. Only peaks for the major rotamer is given. Analytical HPLC gradient 0–95% eluent II in eluent I (11 min total runtime), $t_R$ 3.93 min (>98%, UV$_{230}$). HRMS calcd for $C_{37}H_{55}N_8O_{10}^+$ [M+H]$^+$, 771.4036; found 771.4024. AMC=7-amino-4-methylcoumarin.

**Ac-Gln-Pro-Lys-Lys(D-La)-AMC (4c).** Compound **S2** (35 mg, 0.031 mmol) and D-lactate (8 mg, 0.092 mmol) were dissolved in anh. DMF (1.0 mL) and cooled to 0 °C. *i*Pr$_2$NEt (32 µL, 0.184 mmol) and HATU (33 mg, 0.086 mmol) were added to the reaction mixture, which was stirred for 3 h going towards ambient temperature. Solvent was removed under reduced pressure, to afford the crude intermediate tentatively assigned as Ac-Gln(Trt)-Pro-Lys(Boc)-Lys(D-la)-AMC (HPLC-MS $t_R$ 1.94 min, $m/z$ 1113.6 ([M+H]$^+$, $C_{61}H_{77}N_8O_{12}^*$, Calcd 1113.6), which was used without further purification. TFA/CH$_2$Cl$_2$/H$_2$O (49:49:2, v/v, 1.6 mL) was added to the intermediate and stirred for 1 h at ambient temperature. Solvent was removed under a stream of nitrogen, and preparative reversed-phase HPLC purification afforded the title compound (8 mg, 33% from **S2**) as a white fluffy material.

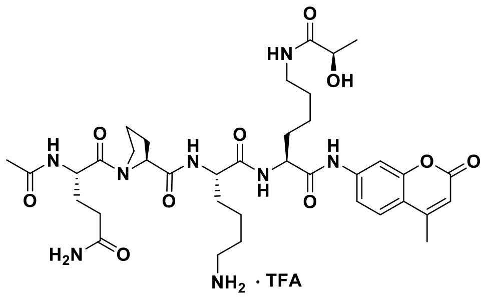


$^1$H NMR (600 MHz, DMSO-$d_6$) δ 10.35 (s, 1H, NH$_{AMC}$), 8.11–8.04 (m, 3H, NH$_{\alpha,Gln}$, NH$_{\alpha,Lys}$ NH$_{\alpha,Lys(lactoyl)}$), 7.80 (d, $J$ = 2.0 Hz, 1H, H8$_{AMC}$), 7.79–7.69 (m, 4H, H5$_{AMC}$, NH$_{\varepsilon,Lys}$), 7.67 (t, $J$ = 6.0 Hz, 1H, NH$_{\varepsilon,Lys(lactoyl)}$), 7.48 (dd, $J$ = 8.7, 2.1 Hz, 1H, H6$_{AMC}$), 7.33 (s, 1H, CONH$_{2,Gln,A}$), 6.81 (s, 1H, CONH$_{2,Gln,B}$), 6.27 (d, $J$ = 1.4 Hz, 1H, H3$_{AMC}$), 4.47 (td, $J$ = 8.2, 5.9 Hz, 1H, H$_{\alpha,Gln}$), 4.39–4.30 (m, 2H, H$_{\alpha,Pro}$, H$_{\alpha,Lys(lactoyl)}$), 4.29–4.22 (m, 1H, H$_{\alpha,Lys}$), 3.92 (q, $J$ = 6.8 Hz, 1H, C(OH)$\underline{H}$CH$_3$), 3.72–3.59 (m, 2H, H$_{\delta,Pro}$), 3.11–3.02 (m, 2H, H$_{\varepsilon,Lys(lactoyl)}$), 2.82–2.73 (m, 2H, H$_{\varepsilon,Lys}$), 2.40 (d, $J$ = 1.3 Hz, 3H, CH$_{3,AMC}$), 2.18–2.10 (m, 2H, H$_{\delta,Gln}$), 2.09–2.00 (m, 1H, H$_{\beta,Pro,A}$), 1.96–1.21 (m, 20H, H$_{\beta,Lys}$, H$_{\gamma,Lys}$, H$_{\delta,Lys}$, H$_{\beta,Lys(lactoyl)}$, H$_{\gamma,Lys(lactoyl)}$, H$_{\delta,Lys(lactoyl)}$, H$_{\beta,Pro,B}$, H$_{\gamma,Pro}$, H$_{\beta,Gln}$, CH$_{3,acetyl}$), 1.17 (d, $J$ = 6.7 Hz, 3H, C(OH)HC$\underline{H}_3$). $^{13}$C NMR (151 MHz, DMSO) δ 174.3 ($\underline{C}$ONH$_{\varepsilon,Lys(lactoyl)}$), 173.9 (CO$_{\delta,Gln}$), 171.9 (CO$_{\alpha,Lys}$), 171.7 (CO$_{Pro}$), 171.4 (CO$_{\alpha,Lys(lactoyl)}$), 170.5 (CO$_{\alpha,Gln}$), 169.2 ($\underline{C}$OCH$_{3,acetyl}$), 160.0 (C2$_{AMC}$), 158.1 (q, $J$ = 32.4 Hz, residual CO$_{TFA}$), 153.6 (C8a$_{AMC}$), 153.1 (C4$_{AMC}$), 142.1 (C7$_{AMC}$), 125.9 (C5$_{AMC}$), 115.2 (C6$_{AMC}$), 115.1 (C4a$_{AMC}$), 112.3 (C3$_{AMC}$), 105.7 (C8$_{AMC}$), 67.2 ($\underline{C}$(OH)HCH$_3$), 59.5 (C$_{\alpha,Pro}$), 53.7 (C$_{\alpha,Lys(lactoyl)}$), 52.3 (C$_{\alpha,Lys}$), 49.8 (C$_{\alpha,Gln}$), 46.9 (C$_{\delta,Pro}$), 38.8 (C$_{\varepsilon,Lys}$), 37.9 (C$_{\varepsilon,Lys(lactoyl)}$), 31.3 (C$_{\gamma,Gln}$), 31.1 (C$_{\beta,Lys}$), 31.0 (C$_{\beta,Lys(lactoyl)}$), 29.1 (C$_{\beta,Pro}$), 28.9 (C$_{\delta,Lys(lactoyl)}$), 27.1 (C$_{\beta,Gln}$), 26.5 (C$_{\delta,Lys}$), 24.5 (C$_{\gamma,Pro}$), 22.8 (C$_{\gamma,Lys(lactoyl)}$), 22.3 (C$_{\gamma,Lys}$), 22.1 (CH$_{3,acetyl}$), 21.1 (C(OH)HCH$_3$), 18.0 (CH$_{3,AMC}$). Two sets of signals (approximately 10:1) were detectable due to rotamers. Only peaks for the major rotamer is given. Analytical HPLC gradient 0–50% eluent II in eluent I (11 min total runtime), $t_R$ 5.20 min (>98%, UV$_{230}$). HRMS calcd for $C_{37}H_{55}N_8O_{10}^+$ [M+H]$^+$, 771.4036; found 771.4027. AMC=7-amino-4-methylcoumarin.

**Ac-Thr-Ala-Arg-Lys(L-La)-AMC (5b).** By the method described for **5c**, the title compound was synthesized using **S4** (8 mg, 0.008 mmol), L-lactate (5 mg, 0.054 mmol), *i*$Pr_2$NEt (19 µL, 0.109 mmol), and HATU (5 mg, 0.012 mmol). Preparative reversed-phase HPLC purification afforded the title compound (2 mg, 26% from **S4**) as a white fluffy material after lyophilization. $^1$H NMR (600 MHz, DMSO-$d_6$) 10.41 (s, 1H, $NH_{AMC}$), 8.09 (d, $J$ = 7.12Hz, 1H, $NH_{\alpha,Lys}$), 8.02–7.93 (m, 2H, $NH_{\alpha,Arg}$, $NH_{Ala}$), 7.86–7.78 (m, 2H, $NH_{Thr}$, $H8_{AMC}$), 7.72 (d, $J$ = 8.7 Hz, 1H, $H5_{AMC}$), 7.67 (t, $J$ = 5.9 Hz, 1H, $NH_{\varepsilon,Lys}$), 7.51–7.43 (m, 2H, $NH_{\delta,Arg}$, $H6_{AMC}$), 6.27 (s, 1H, $H3_{AMC}$), 4.37–4.25 (m, 3H, $H_{\alpha,Ala}$, $H_{\alpha,Arg}$, $H_{\alpha,Lys}$), 4.18 (dd, $J$ = 8.3, 4.4 Hz, 1H, $H_{\alpha,Thr}$), 4.00–3.89 (m, 2H, $H_{\beta,Thr}$, C(OH)$\underline{H}$$CH_3$), 3.14–3.00 (m, 4H, $H_{\delta,Arg}$, $H_{\varepsilon,Lys}$), 2.40 (s, 3H, $CH_{3,AMC}$), 1.91 (s, 3H, $CH_{3,acetyl}$), 1.78–1.20 (m, 13H, $H_{\beta,Lys}$, $H_{\gamma,Lys}$, $H_{\delta,Lys}$, $H_{\beta,Arg}$, $H_{\gamma,Arg}$, $H_{\beta,Ala}$), 1.16 (d, $J$ = 6.7 Hz, 3H, C(OH)HC$\underline{H}_3$), 1.05 (d, $J$ = 6.3 Hz, 3H, $H_{\gamma,Thr}$). $^{13}$C NMR (151 MHz, DMSO) δ 174.3 ($\underline{C}$ONH$_{\varepsilon,Lys}$), 172.3 ($CO_{Ala}$), 171.4 ($CO_{Arg}$), 171.3 ($CO_{\alpha,Lys}$), 170.1 ($CO_{Thr}$), 169.8 ($\underline{C}$OCH$_{3,acetyl}$), 160.0 ($C2_{AMC}$), 156.6 (NHC(=NH)$NH_2$), 153.6 ($C8a_{AMC}$), 153.1 ($C4_{AMC}$), 142.1 ($C7_{AMC}$), 125.9 ($C5_{AMC}$), 115.2 ($C6_{AMC}$), 115.1 ($C4a_{AMC}$), 112.3 ($C3_{AMC}$), 105.7 ($C8_{AMC}$), 67.2 ($\underline{C}$(OH)H$CH_3$), 66.5 ($C_{\beta,Thr}$), 58.4 ($C_{\alpha,Thr}$), 53.7 ($C_{\alpha,Lys}$), 52.1 ($C_{\alpha,Arg}$), 48.3 ($C_{\alpha,Ala}$), 40.4 ($C_{\delta,Arg}$), 37.9 ($C_{\varepsilon,Lys}$), 31.4 ($C_{\beta,Lys}$), 28.97 ($C_{\beta,Arg}$ / $C_{\delta,Lys}$), 28.95 ($C_{\beta,Arg}$ / $C_{\delta,Lys}$), 24.9 ($C_{\gamma,Arg}$), 22.8 ($C_{\gamma,Lys}$), 22.6 ($CH_{3,acetyl}$), 21.1 (C(OH)H$\underline{C}$H$_3$), 19.7 ($C_{\gamma,Thr}$), 18.0 ($CH_{3,AMC}$), 17.9 ($C_{\beta,Ala}$). Analytical HPLC gradient 0–95% eluent II in eluent I (11 min total runtime), $t_R$ 4.04 min (>98%, $UV_{230}$). HRMS calcd for $C_{34}H_{52}N_9O_{10}^+$ $[M+H]^+$, 746.3832; found 746.3826. AMC=7-amino-4-methylcoumarin.

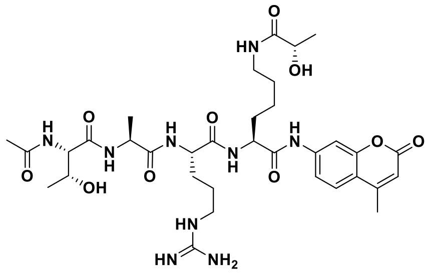


**Ac-Thr-Ala-Arg-Lys(D-La)-AMC (5c).** Compound **S4** (7 mg, 0.007 mmol) and sodium D-lactate (5 mg, 0.041 mmol) were dissolved in anh. DMF (1.0 mL) and cooled to 0 °C. *i*$Pr_2$NEt (14 µL, 0.082 mmol) and HATU (4 mg, 0.010 mmol) were added to the reaction mixture, which was stirred overnight going towards ambient temperature. Additional HATU (4 mg, 0.010 mmol) was added, and the reaction mixture was stirred 1 h until all starting material was consumed. Solvent was removed under reduced pressure to afford the crude intermediate tentatively assigned as Ac-Thr(tBu)-Ala-Arg(Pbf)-Lys(D-la)-AMC (HPLC-MS $t_R$ 1.77 min, $m/z$ 1054.6 ($[M+H]^+$, $C_{51}H_{76}N_9O_{13}S^+$, Calcd 1054.5), which was used without further purification. TFA/TIPS/$H_2O$ (95:2.5:2.5, v/v, 1.0 mL) was added to the intermediate, which was stirred for 1 h at ambient temperature. Solvent was removed under a stream of nitrogen, and preparative reversed-phase HPLC purification afforded the title compound (1 mg, 25% from **S4**), as a white fluffy material after lyophilization. $^1$H NMR (600 MHz, DMSO-$d_6$) δ 10.41 (s, 1H, $NH_{AMC}$), 8.09 (d, $J$ = 7.1 Hz, 1H, $NH_{\alpha,Lys}$), 8.04–7.94 (m, 2H, $NH_{\alpha,Arg}$, $NH_{Ala}$), 7.86–7.78 (m, 2H, $NH_{Thr}$, $H8_{AMC}$), 7.72 (d, $J$ = 8.5 Hz, 1H, $H5_{AMC}$), 7.67 (t, $J$ = 6.0 Hz, 1H, $NH_{\varepsilon,Lys}$), 7.52–7.44 (m, 2H, $NH_{\delta,Arg}$, $H6_{AMC}$), 6.30–6.25 (m, 1H, $H3_{AMC}$), 4.37–4.26 (m, 3H, $H_{\alpha,Ala}$, $H_{\alpha,Arg}$, $H_{\alpha,Lys}$), 4.18 (dd, $J$ = 8.2, 4.3 Hz, 1H, $H_{\alpha,Thr}$), 4.00–3.89 (m, 2H, $H_{\beta,Thr}$, C(OH)$\underline{H}$$CH_3$), 3.10 (q, $J$ = 6.6 Hz, 2H, $H_{\delta,Arg}$), 3.06 (q, $J$ = 6.4 Hz, 3H, $H_{\varepsilon,Lys}$), 2.40 (s, 3H, $CH_{3,AMC}$), 1.91 (s, 3H, $CH_{3,acetyl}$), 1.79–1.21 (m, 13H, $H_{\beta,Lys}$, $H_{\gamma,Lys}$, $H_{\delta,Lys}$, $H_{\beta,Arg}$, $H_{\gamma,Arg}$, $H_{\beta,Ala}$), 1.17 (d, $J$ = 6.6 Hz, 3H, C(OH)HC$\underline{H}_3$), 1.05 (d, $J$ = 6.2 Hz, 3H, $H_{\gamma,Thr}$). $^{13}$C NMR (151 MHz, DMSO) δ 174.3

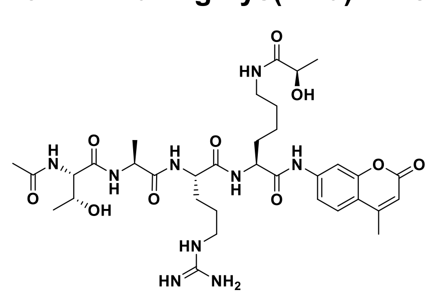

($\underline{C}$ONH$_{\varepsilon,\mathrm{Lys}}$), 172.3 (CO$_{\mathrm{Ala}}$), 171.4 (CO$_{\mathrm{Arg}}$), 171.3 (CO$_{\alpha,\mathrm{Lys}}$), 170.1 (CO$_{\mathrm{Thr}}$), 169.8 ($\underline{C}$OCH$_{3,\mathrm{acetyl}}$), 160.0 (C2$_{\mathrm{AMC}}$), 156.6 (NHC(=NH)NH$_2$), 153.6 (C8a$_{\mathrm{AMC}}$), 153.1 (C4$_{\mathrm{AMC}}$), 142.1 (C7$_{\mathrm{AMC}}$), 125.9 (C5$_{\mathrm{AMC}}$), 115.2 (C6$_{\mathrm{AMC}}$), 115.1 (C4a$_{\mathrm{AMC}}$), 112.3 (C3$_{\mathrm{AMC}}$), 105.7 (C8$_{\mathrm{AMC}}$), 67.2 ($\underline{C}$(OH)HCH$_3$), 66.5 (C$_{\beta,\mathrm{Thr}}$), 58.4 (C$_{\alpha,\mathrm{Thr}}$), 53.7 (C$_{\alpha,\mathrm{Lys}}$), 52.1 (C$_{\alpha,\mathrm{Arg}}$), 48.3 (C$_{\alpha,\mathrm{Ala}}$), 40.4 (C$_{\delta,\mathrm{Arg}}$), 37.9 (C$_{\varepsilon,\mathrm{Lys}}$), 31.4 (C$_{\beta,\mathrm{Lys}}$), 29.0 (C$_{\beta,\mathrm{Arg}}$), 28.9 (C$_{\delta,\mathrm{Lys}}$), 24.9 (C$_{\gamma,\mathrm{Arg}}$), 22.8 (C$_{\gamma,\mathrm{Lys}}$), 22.6 (CH$_{3,\mathrm{acetyl}}$), 21.1 (C(OH)$\underline{C}$H$_3$), 19.7 (C$_{\gamma,\mathrm{Thr}}$), 18.0 (CH$_{3,\mathrm{AMC}}$), 17.9 (C$_{\beta,\mathrm{Ala}}$). Analytical HPLC gradient 0–95% eluent II in eluent I (11 min total runtime), $t_R$ 4.05 min (>98%, UV$_{230}$). HRMS calcd for $C_{34}H_{52}N_9O_{10}^+$ [M+H]$^+$, 746.3832; found 746.3826. AMC=7-amino-4-methylcoumarin.

*Additional substrate sources*

| Code | Sequence | Source |
|---|---|---|
| **2a** | Ac-Leu-Gly-Lys(Ac)-AMC | (*7, 8*) |
| **2d** | Ac-Leu-Gly-Lys(Tfa)-AMC | (*8*) |
| **2f** | Ac-Leu-Gly-Lys(Pro)-AMC | (*9*) |
| **2g** | Ac-Leu-Gly-Lys(*i*-But)-AMC | (*10*) |
| **2j** | Ac-Leu-Gly-Lys(But)-AMC | (*9*) |
| **2k** | Ac-Leu-Gly-Lys(*i*-Val)-AMC | (*10*) |
| **2m** | Ac-Leu-Gly-Lys(Cr)-AMC | (*5*) |
| **3e** | Ac-Glu-Thr-Asp-Lys(Myr)-AMC | (*10, 11*) |
| **4a** | Ac-Gln-Pro-Lys-Lys(Ac)-AMC | Enzo Life Sciences and (*12*) |
| **4f** | Ac-Gln-Pro-Lys-Lys(Pro)-AMC | (*12*) |
| **4g** | Ac-Gln-Pro-Lys-Lys(*i*-But)-AMC | (*12*) |
| **4h** | Ac-Gln-Pro-Lys-Lys(Glu)-AMC | (*13*) |
| **4i** | Ac-Gln-Pro-Lys-Lys(Dec)-AMC | (*11*) |

*Synthesis of non-fluorogenic histone peptide substrates*

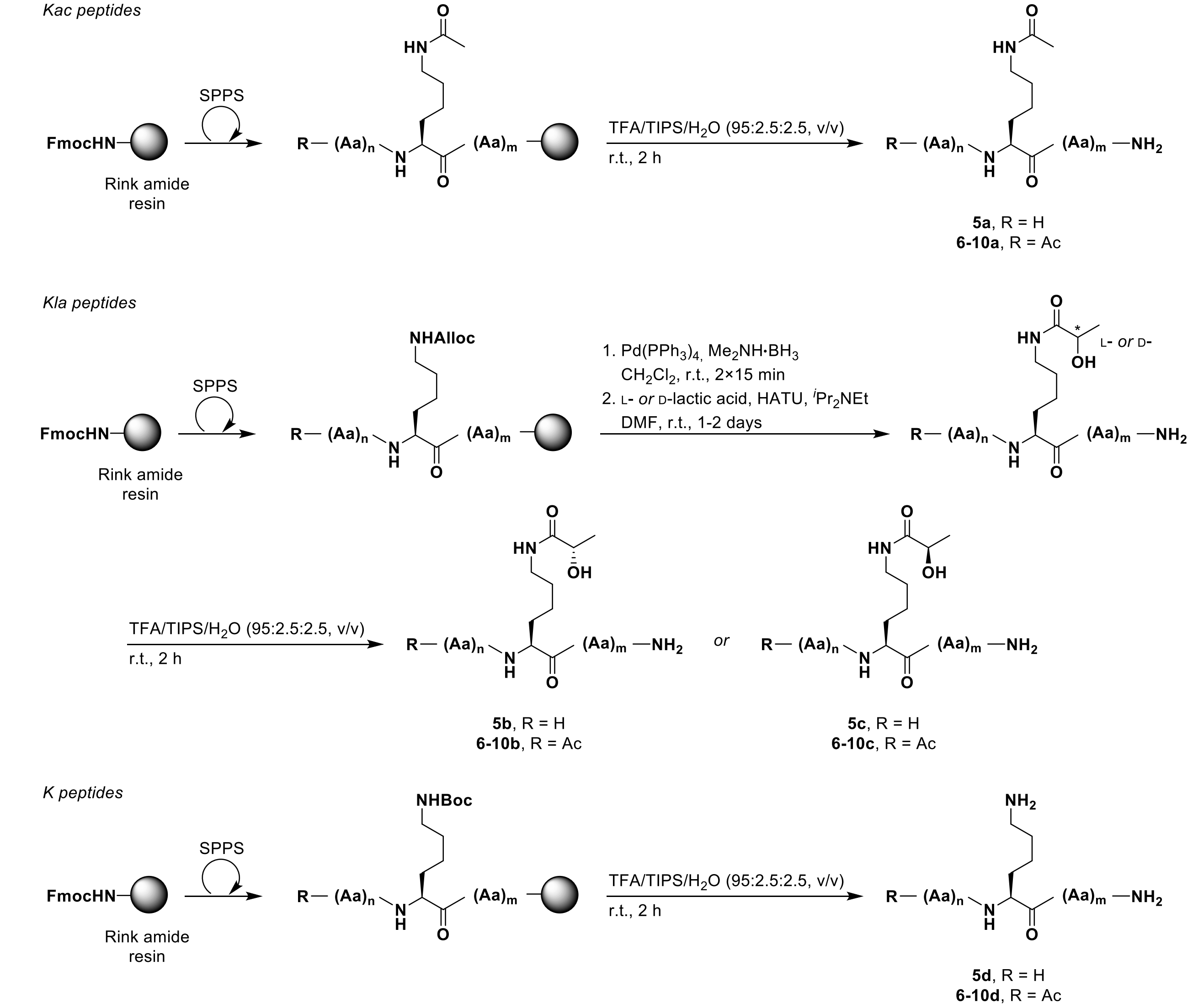


**Scheme S7. Synthesis of peptides 5–10 as Kac (a), K(L-la) (b), K(D-la) (c) or K (d) versions.**

Non-fluorogenic peptide substrates were synthesized by automated solid phase peptide synthesis (SPPS) using standard Fmoc/*t*Bu chemistry on a Biotage Syro Wave synthesizer (Scheme S7). The following commercially available protected amino acids were used: Fmoc-Ala-OH, Fmoc-Arg(Pbf)-OH, Fmoc-Gln(Trt)-OH, Fmoc-Glu(*t*Bu)-OH, Fmoc-Gly-OH, Fmoc-Leu-OH, Fmoc-Lys(Ac)-OH, Fmoc-Lys(Alloc)-OH, Fmoc-Lys(Boc)-OH, Fmoc-Phe-OH, Boc-Pro-OH (final amino acid for sequence **5**), Fmoc-Pro-OH, Fmoc-Ser(*t*Bu)-OH, Fmoc-Thr(*t*Bu)-OH and Fmoc-Trp(Boc)-OH. SPPS was performed on 0.04–0.08 mmol scale using preloaded TentaGel® S RAM resin (0.24 mmol/g, Rapp Polymere; #S30023). Fmoc deprotection was performed twice: (1) piperidine in DMF (2:3, v/v) for 3 min and (2) piperidine in DMF (1:4, v/v) for 2×8 min. Deprotection was followed by washing with DMF (2×45 s), $CH_2Cl_2$ (45 s), and DMF (2×45 s). Coupling reactions were performed as double couplings using

Fmoc-Xaa-OH (5.0 equiv to the resin loading), HBTU (5 equiv) and *i*$Pr_2NEt$ (10 equiv, 2.0 M in NMP) in DMF (final concentration = 0.2 M) for 2×40 min.

On-resin Alloc deprotection was performed by addition of borane dimethylamine complex (5 equiv.) and $Pd(PPh_3)_4$ (10 mol%) in anh. $CH_2Cl_2$ (2.5 mL) to the resin for 2×15 min, followed by washing with $CH_2Cl_2$ (3×4.0 mL), DMF (3×4.0 mL) and $CH_2Cl_2$ (3×4.0 mL). Then, on-resin lysine lactylation was performed using L- *or* D-lactic acid (5 equiv.), HATU (4.5 equiv.), and and *i*$Pr_2NEt$ (10 equiv) in DMF (2.5 mL) for 1–2 days, followed by washing with DMF (3×4.0 mL) and $CH_2Cl_2$ (3×4.0 mL). Alloc deprotection and on-resin lactylation were repeated when test cleavages showed remaining starting material.

Cleavage and global deprotection of the peptides was performed with a mixture of TFA/TIPS/$H_2O$ (95:2.5:2.5, v/v, 4 mL) for 2 h. TFA was removed under a stream of nitrogen, and the crude peptides were triturated in ice-cold diethylether and purified by preparative reversed-phase HPLC to afford the desired products as white fluffy powder after lyophilization (Scheme S8).

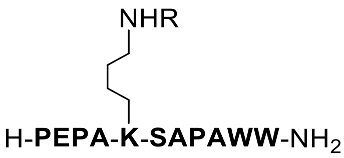


H2BK5
**5a**, R = Ac
**5b**, R = (L-La)
**5c**, R = (D-La)
**5d**, R = H
**CMY233_5,6,7,8**

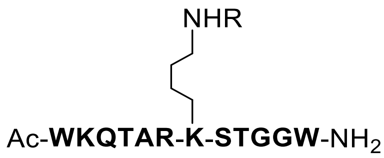


H3K9
**6a**, R = Ac
**6b**, R = (L-La)
**6c**, R = (D-La)
**6d**, R = H
**ALN-255-1,2, ALN-281a-b**

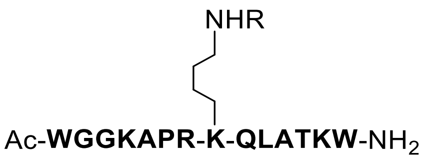


H3K18
**7a**, R = Ac
**7b**, R = (L-La)
**7c**, R = (D-La)
**7d**, R = H
**ALN-274, SAM-64,69,76**

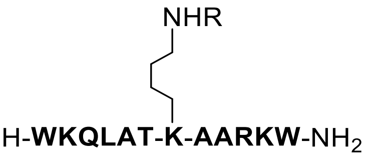


H3K23
**8a**, R = Ac
**8b**, R = (L-La)
**8c**, R = (D-La)
**8d**, R = H
**CMY233_1,2,3,4**

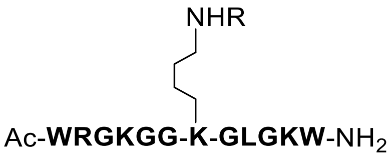


H4K8
**9a**, R = Ac
**9b**, R = (L-La)
**9c**, R = (D-La)
**9d**, R = H
**CMY231_5,6,7,8**

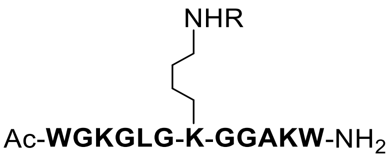


H4K12
**10a**, R = Ac
**10b**, R = (L-La)
**10c**, R = (D-La)
**10d**, R = H
**CMY231_1,2,3,4**

**Scheme S8. Structure of non-fluorogenic peptide substrates and lysine controls.**

*Purity and high-resolution mass spectrometry (HRMS) of non-fluorogenic peptides*

| | R = | Purity* | $t_R$ (min) | Formula | HRMS ($m/z$) | found | (calcd) |
|---|---|---|---|---|---|---|---|
| **5a** | Ac | 99% | 4.68 | $C_{62}H_{85}N_{15}O_{15}$ | $[M+H]^+$ | 1280.64216 | (1280.64223) |
| **5b** | L-La | 98% | 4.68 | $C_{63}H_{87}N_{15}O_{16}$ | $[M+H]^+$ | 1310.65211 | (1310.65280) |
| **5c** | D-La | 98% | 4.68 | $C_{63}H_{87}N_{15}O_{16}$ | $[M+H]^+$ | 1310.65224 | (1310.65280) |
| **5d** | H | 99% | 4.49 | $C_{60}H_{83}N_{15}O_{14}$ | $[M+H]^+$ | 1238.63224 | (1238.63167) |
| **6a** | Ac | 95% | 4.10 | $C_{67}H_{101}N_{21}O_{18}$ | $[M+H]^+$ | 1488.7724 | (1488.7705) |
| **6b** | L-La | 95% | 4.13 | $C_{68}H_{103}N_{21}O_{19}$ | $[M+H]^+$ | 1518.7804 | (1518.7810) |
| **6c** | D-La | 95% | 4.13 | $C_{68}H_{103}N_{21}O_{19}$ | $[M+H]^+$ | 1518.7847 | (1518.7810) |
| **6d** | H | 94% | 3.94 | $C_{65}H_{99}N_{21}O_{17}$ | $[M+H]^+$ | 1446.7613 | (1446.7599) |
| **7a** | Ac | 96% | 4.19 | $C_{80}H_{124}N_{24}O_{18}$ | $[M+H]^+$ | 1709.9610 | (1709.9596) |
| **7b** | L-La | 97% | 4.40 | $C_{81}H_{126}N_{24}O_{19}$ | $[M+H]^+$ | 1739.9722 | (1739.9702) |
| **7c** | D-La | 99% | 4.40 | $C_{81}H_{126}N_{24}O_{19}$ | $[M+H]^+$ | 1739.9720 | (1739.9702) |
| **7d** | H | 98% | 4.18 | $C_{78}H_{122}N_{24}O_{17}$ | $[M+H]^+$ | 1667.9512 | (1667.9491) |
| **8a** | Ac | 98% | 5.00 | $C_{74}H_{116}N_{22}O_{16}$ | $[M+2H]^{2+}$ | 785.45399 | (785.45426) |
| **8b** | L-La | 94% | 5.02 | $C_{75}H_{118}N_{22}O_{17}$ | $[M+2H]^{2+}$ | 800.45976 | (800.45954) |
| **8c** | D-La | 96% | 5.01 | $C_{75}H_{118}N_{22}O_{17}$ | $[M+2H]^{2+}$ | 800.45980 | (800.45954) |
| **8d** | H | 97% | 4.65 | $C_{72}H_{114}N_{22}O_{15}$ | $[M+2H]^{2+}$ | 764.44908 | (764.44898) |
| **9a** | Ac | 97% | 4.31 | $C_{66}H_{101}N_{21}O_{14}$ | $[M+2H]^{2+}$ | 706.89903 | (706.89912) |
| **9b** | L-La | 96% | 4.31 | $C_{67}H_{103}N_{21}O_{15}$ | $[M+2H]^{2+}$ | 721.90423 | (721.90440) |
| **9c** | D-La | 97% | 4.30 | $C_{67}H_{103}N_{21}O_{15}$ | $[M+H]^+$ | 1442.80226 | (1442.69300) |
| **9d** | H | 98% | 4.11 | $C_{64}H_{99}N_{21}O_{13}$ | $[M+2H]^{2+}$ | 685.89396 | (685.89384) |
| **10a** | Ac | 98% | 4.52 | $C_{63}H_{94}N_{18}O_{14}$ | $[M+H]^+$ | 1327.72851 | (1327.72697) |
| **10b** | L-La | 98% | 4.53 | $C_{64}H_{96}N_{18}O_{15}$ | $[M+H]^+$ | 1357.73773 | (1357.73753) |
| **10c** | D-La | 97% | 4.53 | $C_{64}H_{96}N_{18}O_{15}$ | $[M+H]^+$ | 1357.73711 | (1357.73753) |
| **10d** | H | 99% | 4.31 | $C_{61}H_{92}N_{18}O_{13}$ | $[M+H]^+$ | 1285.71807 | (1285.71640) |

*Peptide purity measured by integration of HPLC chromatograms at 215 nm or 230 nm. Please find HPLC traces below. Retention times correspond to linear gradients of eluent III and eluent IV, rising linearly from] 0% to 95% of IV during $t$ = 1−11.

*NMR spectra*

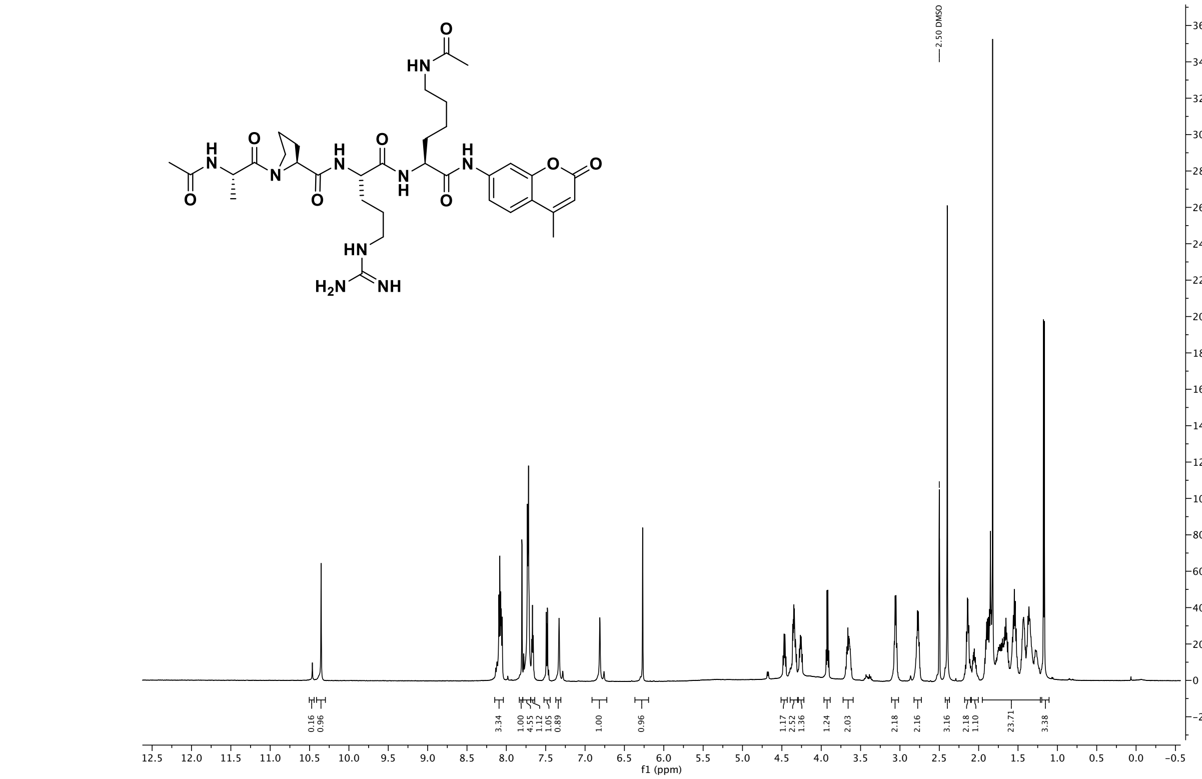


$^{1}$H NMR spectra (600 MHz, DMSO-$d_6$) of compound **1a**

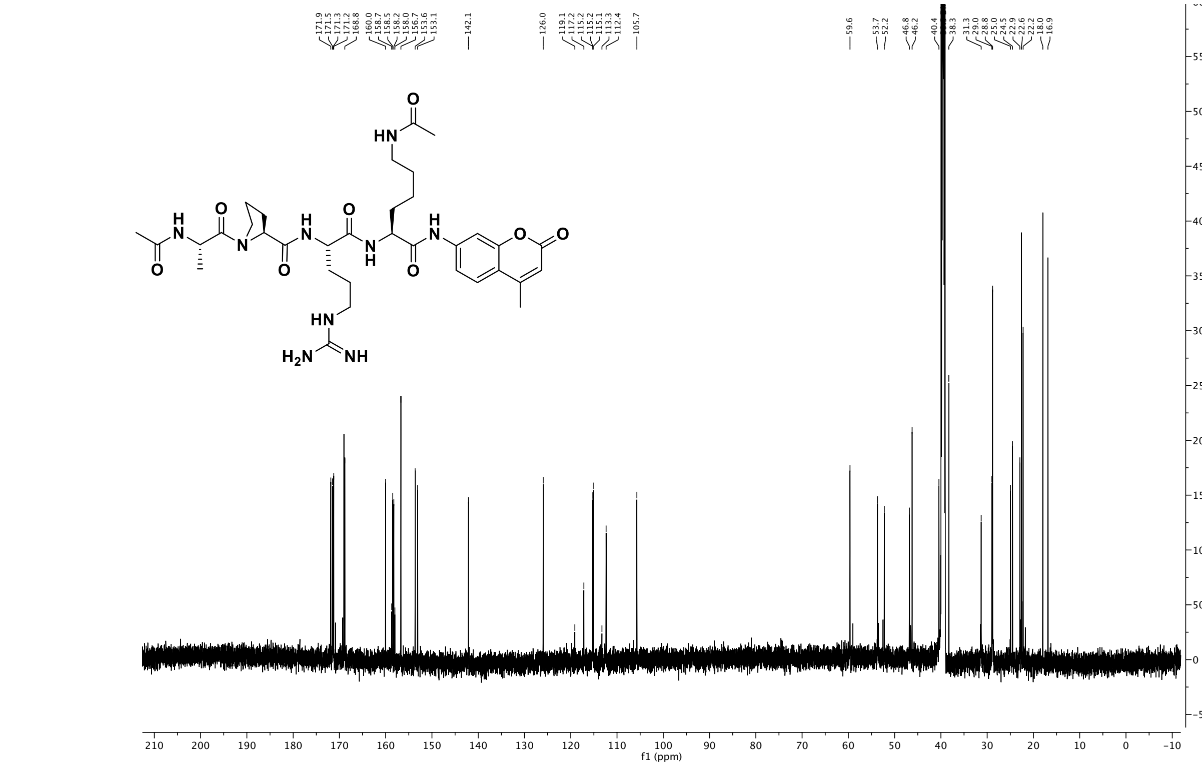


$^{13}$C NMR spectra (151 MHz, DMSO) of compound **1a**

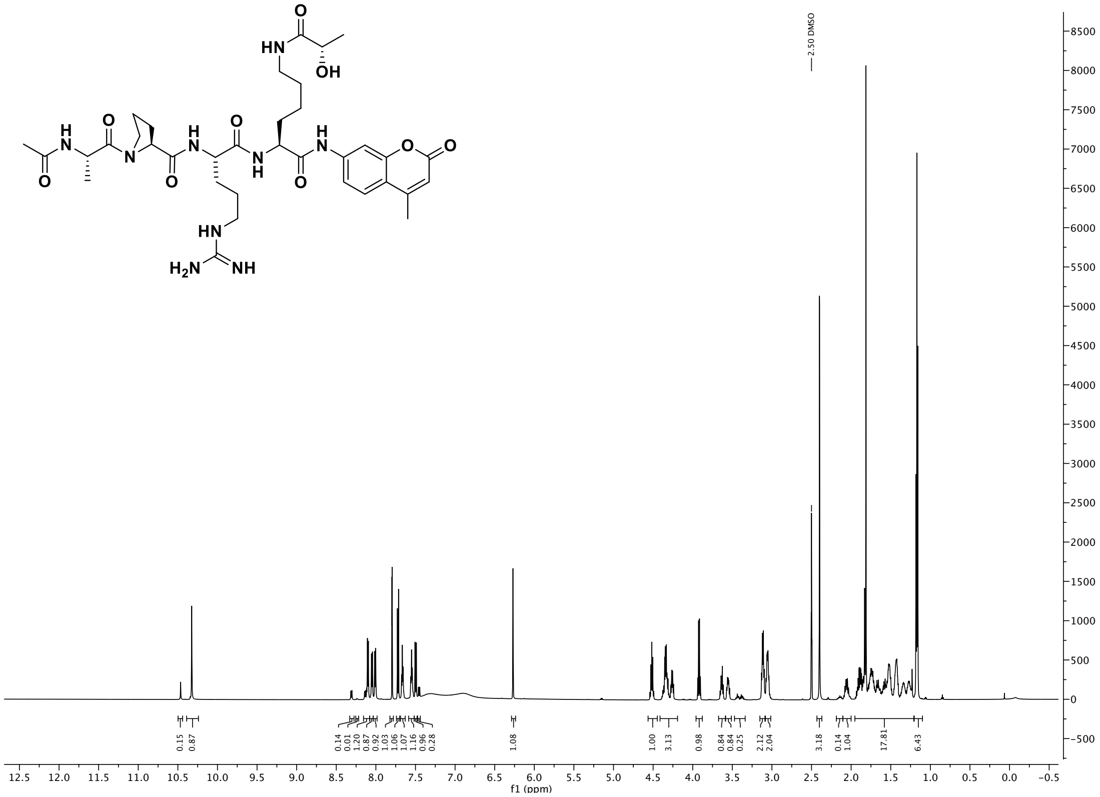

$^{1}H$ NMR spectra (600 MHz, DMSO-$d_6$) of compound **1b**

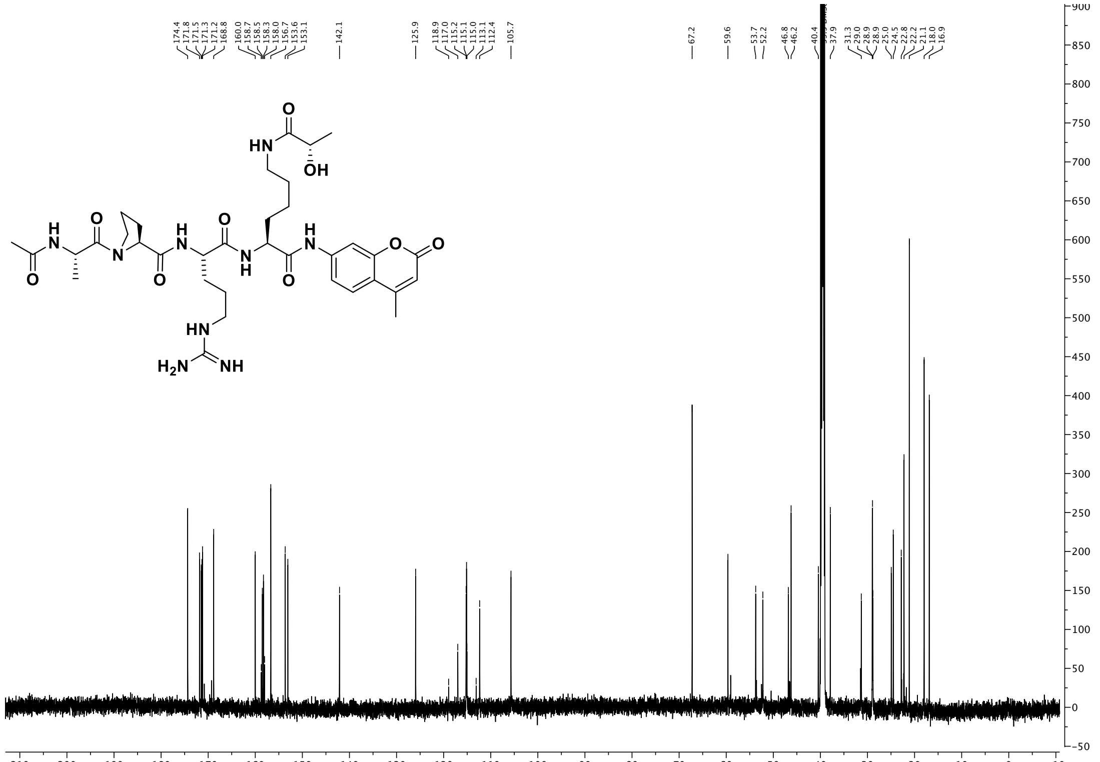

$^{13}C$ NMR spectra (151 MHz, DMSO) of compound **1b**

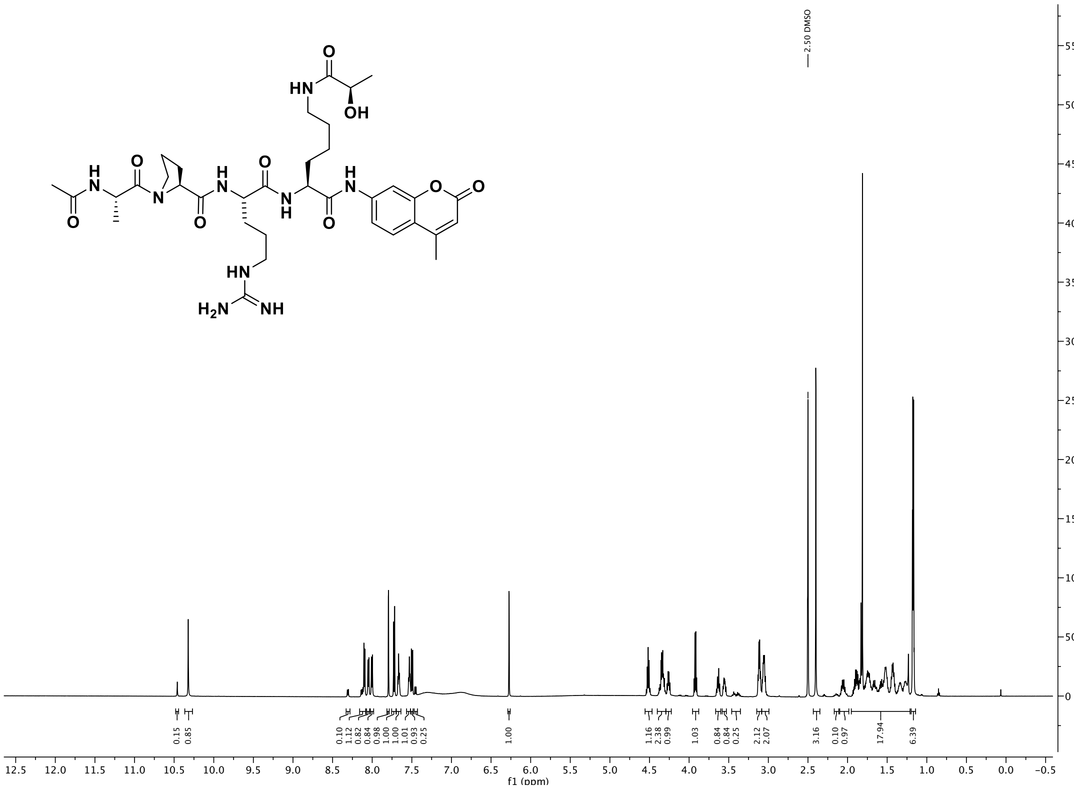


[1]H NMR spectra (600 MHz, DMSO-$d_6$) of compound **1c**

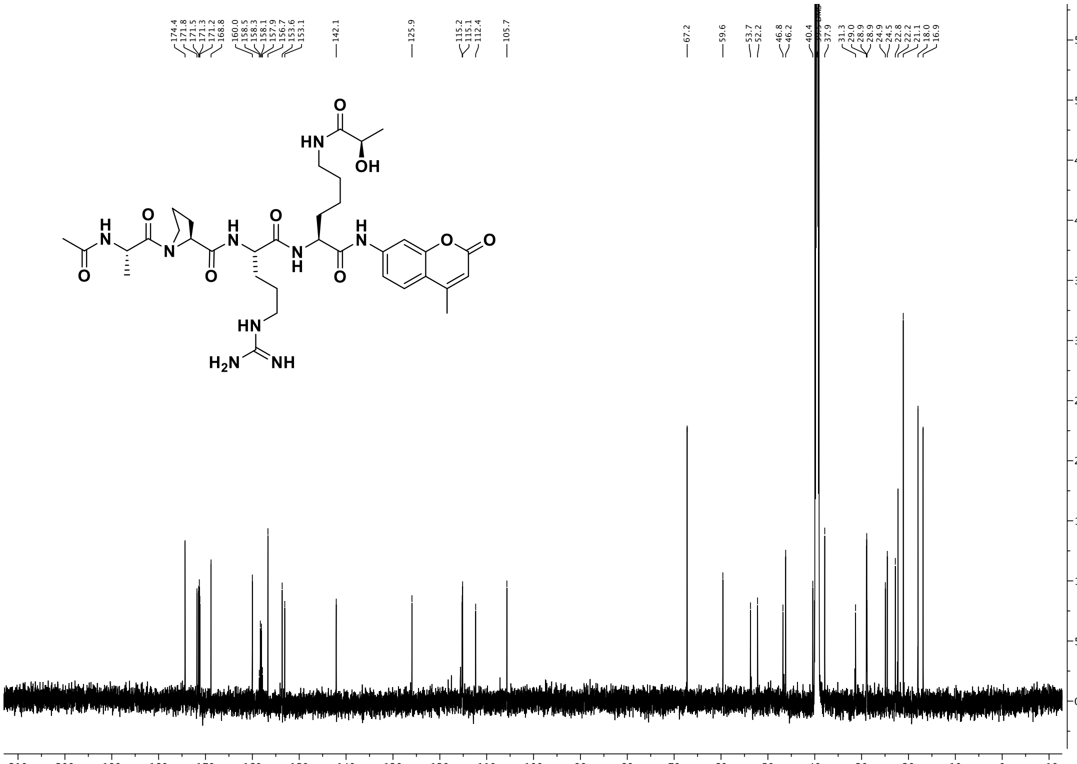


[13]C NMR spectra (151 MHz, DMSO) of compound **1c**

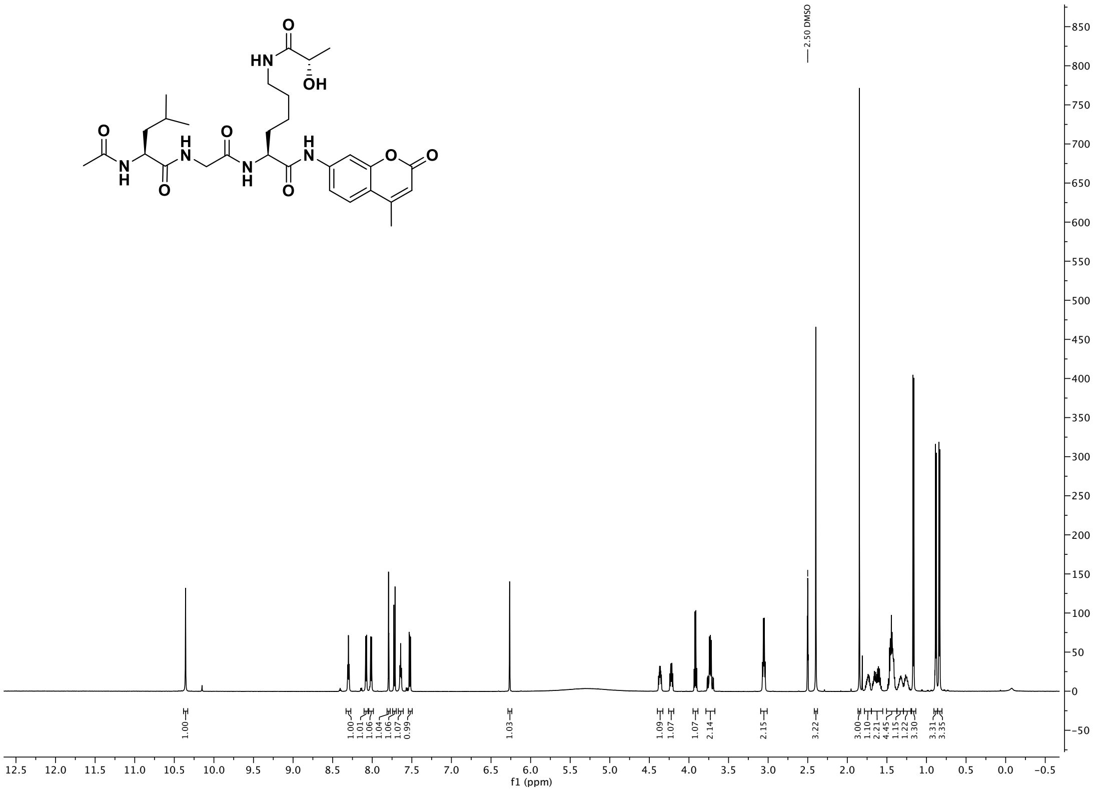

[1]H NMR spectra (600 MHz, DMSO-$d_6$) of compound **2b**

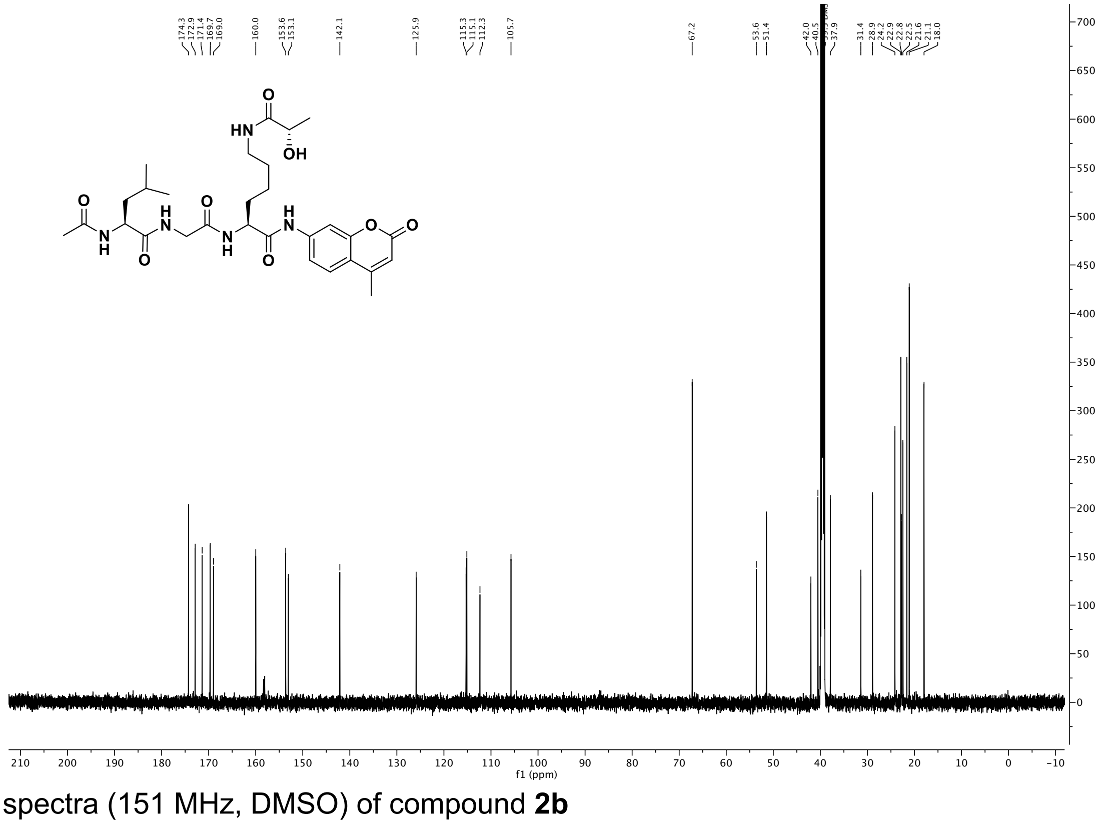

[13]C NMR spectra (151 MHz, DMSO) of compound **2b**

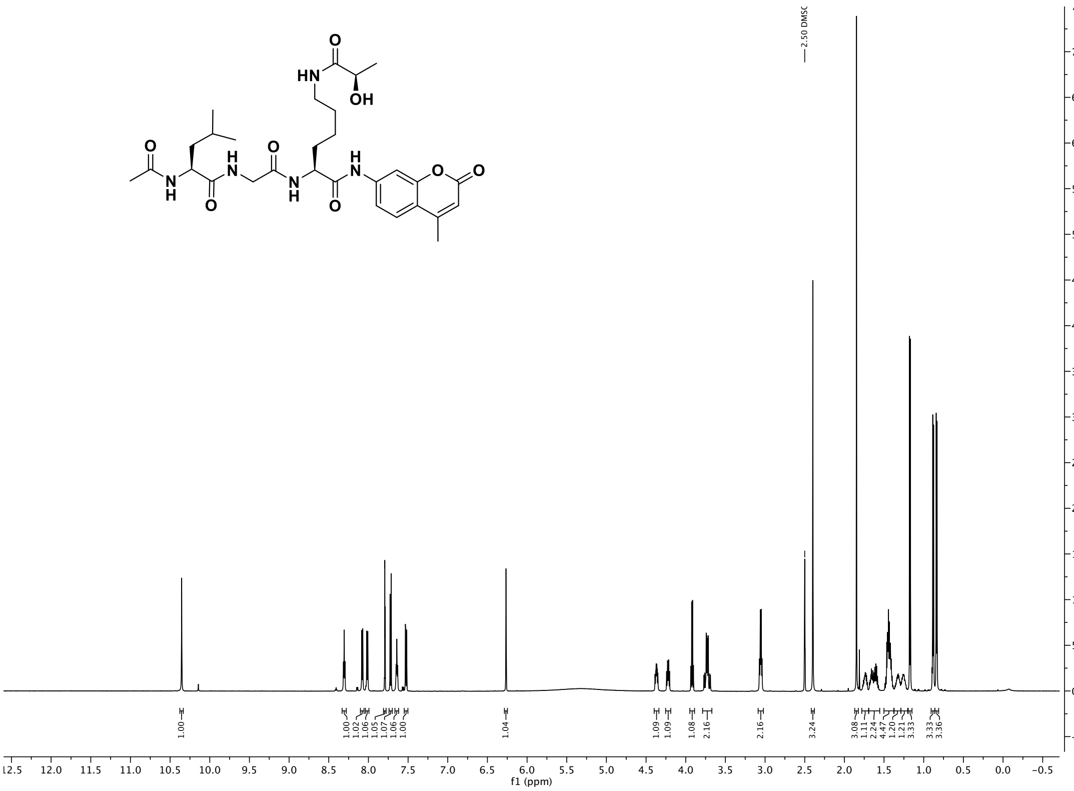

$^{1}H$ NMR spectra (600 MHz, DMSO-$d_6$) of compound **2c**

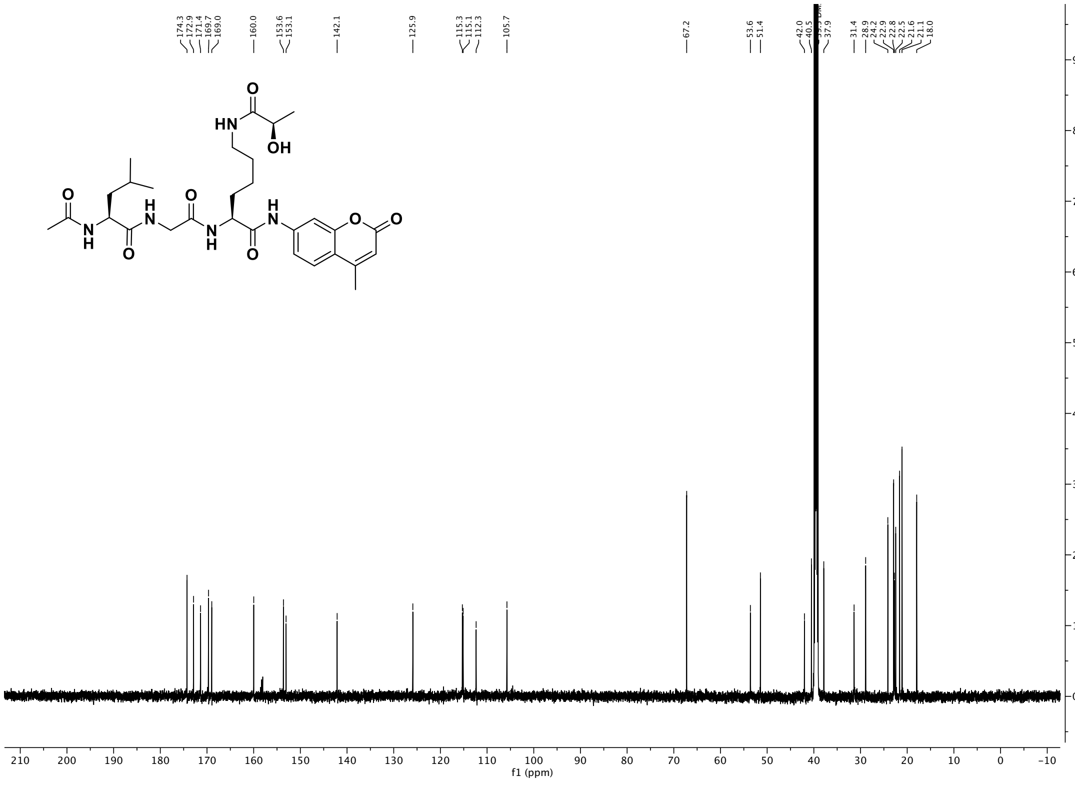

$^{13}C$ NMR spectra (151 MHz, DMSO) of compound **2c**

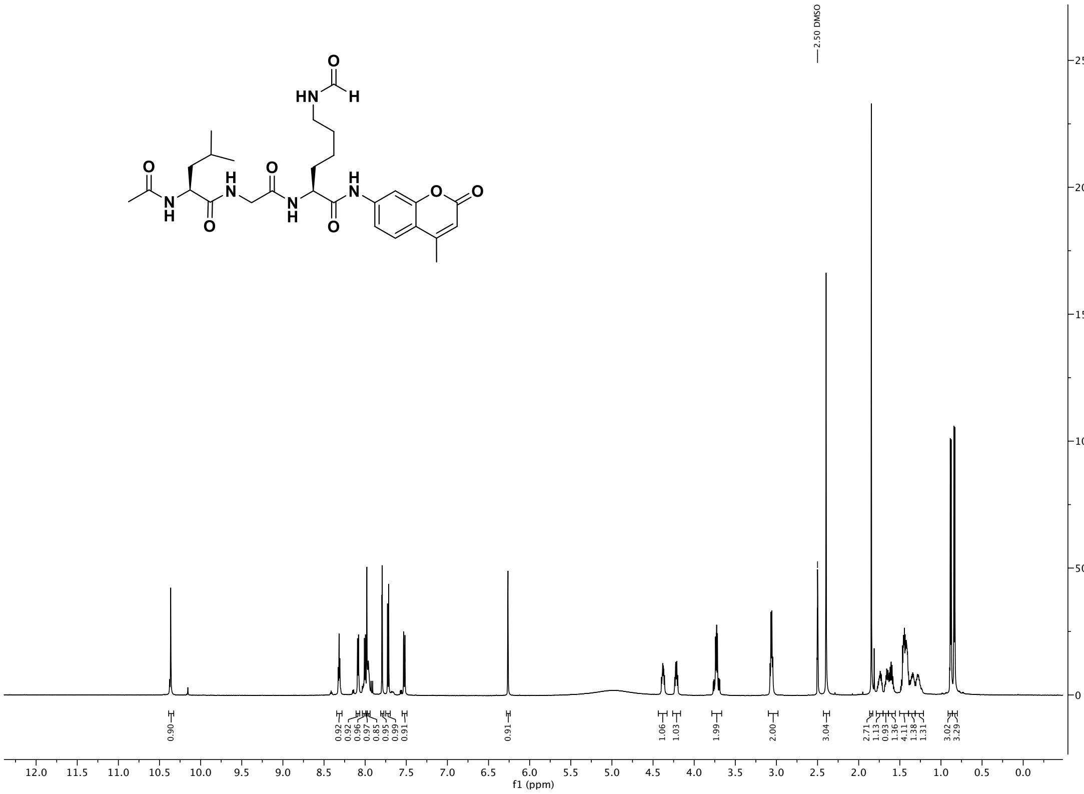

[1]H NMR spectra (600 MHz, DMSO-$d_6$) of compound **2l**

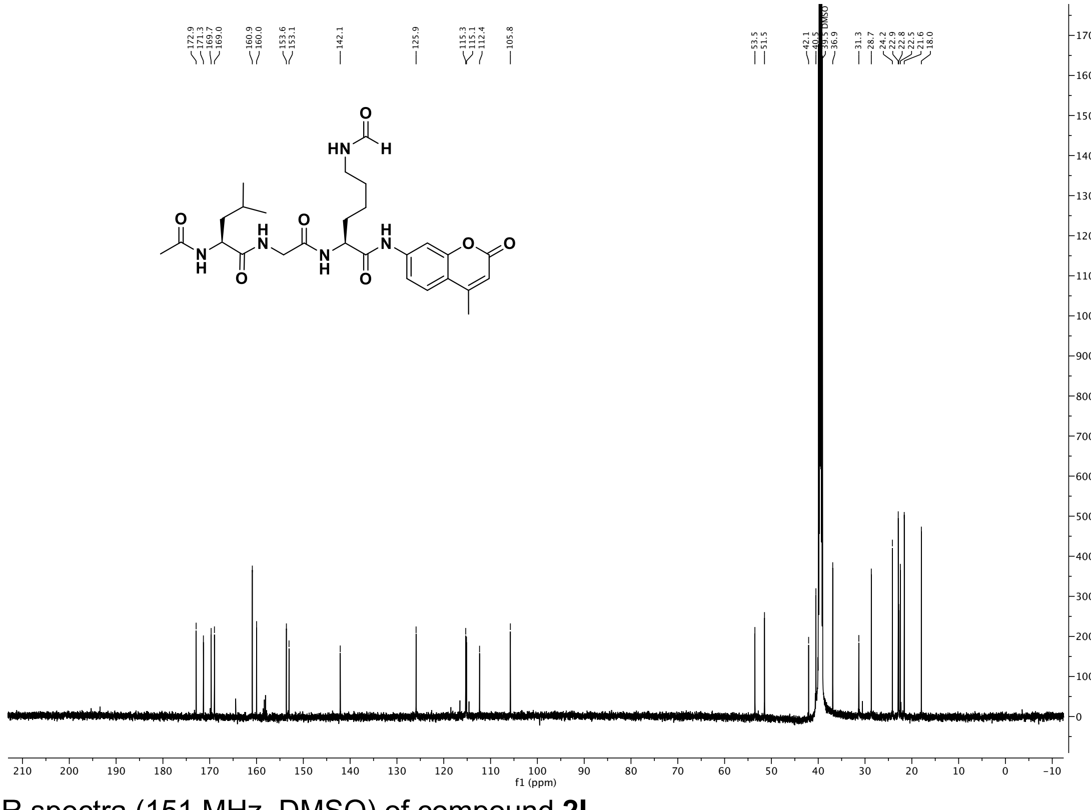

[13]C NMR spectra (151 MHz, DMSO) of compound **2l**

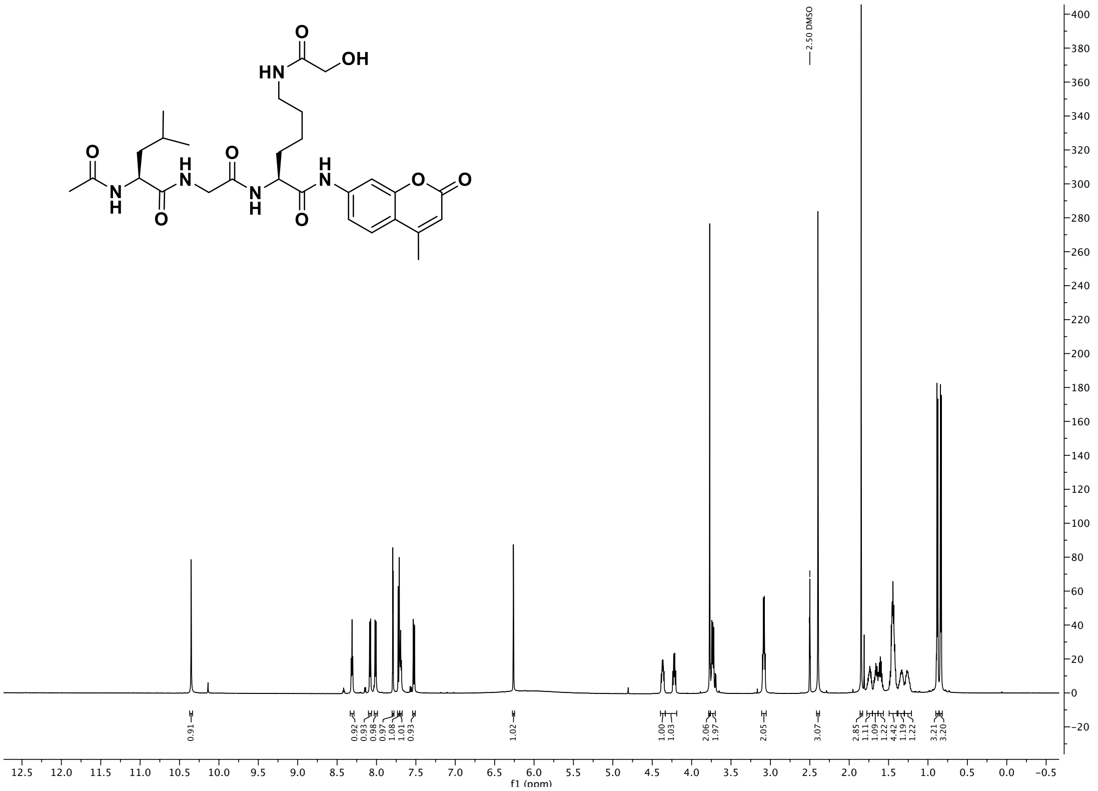

$^{1}H$ NMR spectra (600 MHz, DMSO-$d_6$) of compound **2n**

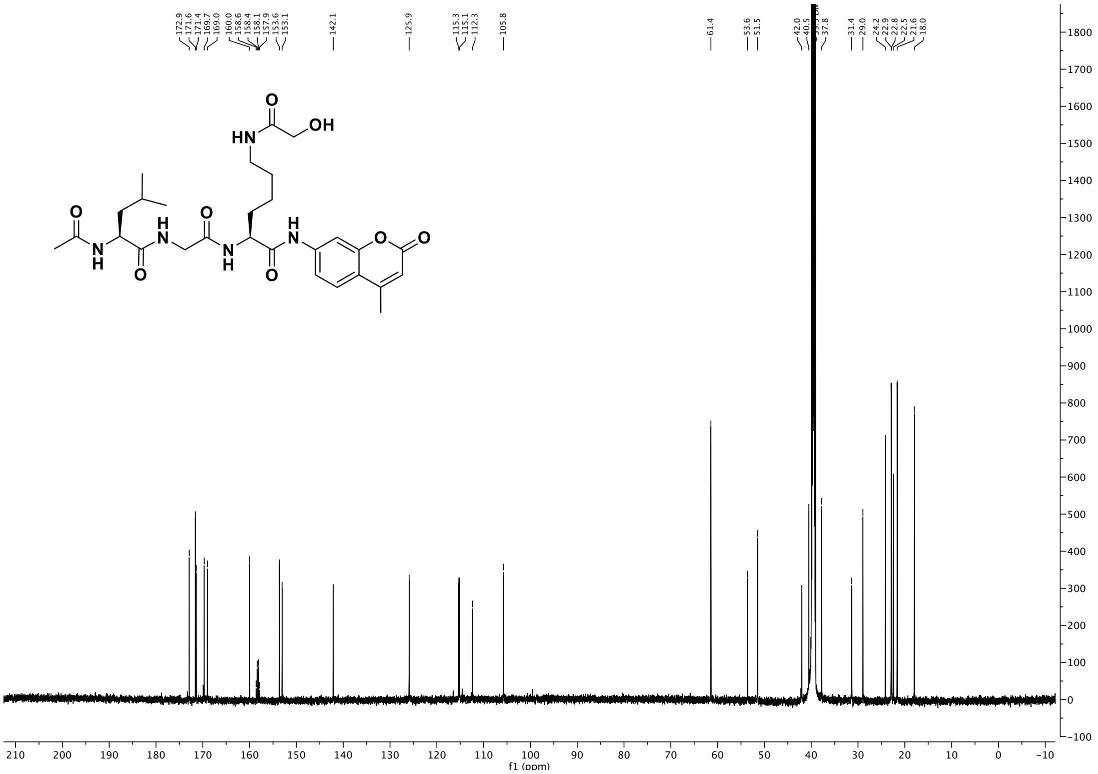

$^{13}C$ NMR spectra (151 MHz, DMSO) of compound **2n**

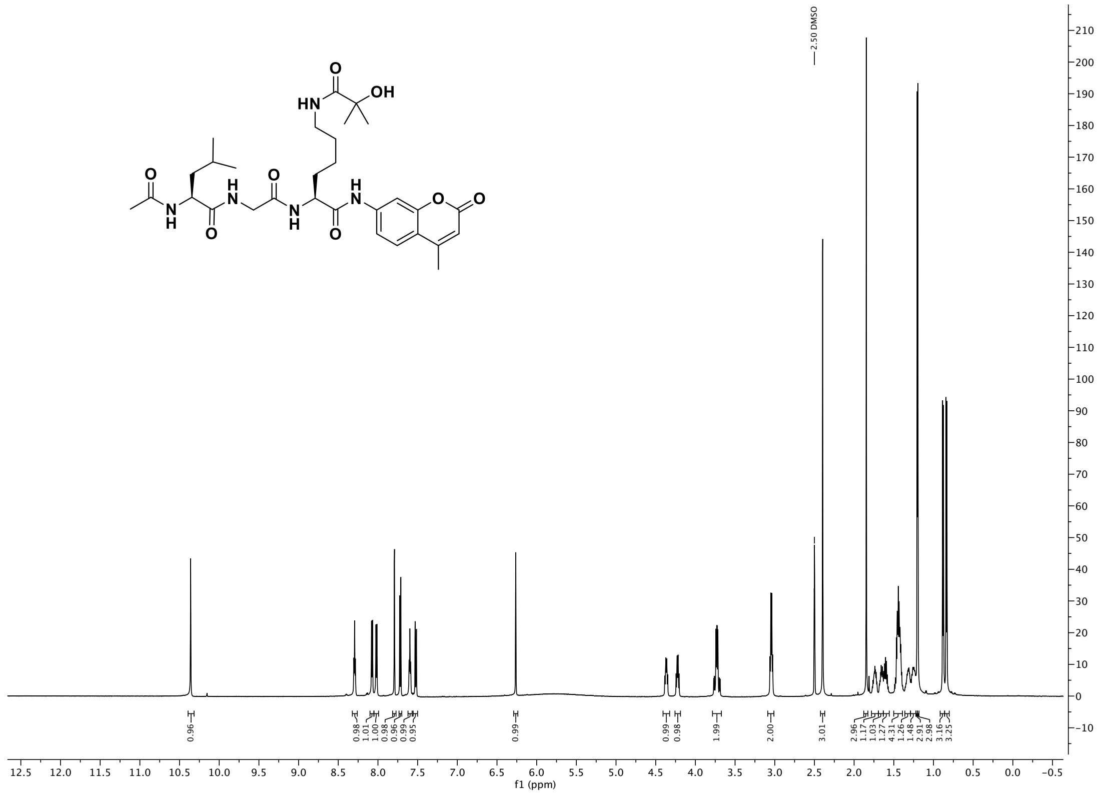

[1]H NMR spectra (600 MHz, DMSO-$d_6$) of compound **2o**

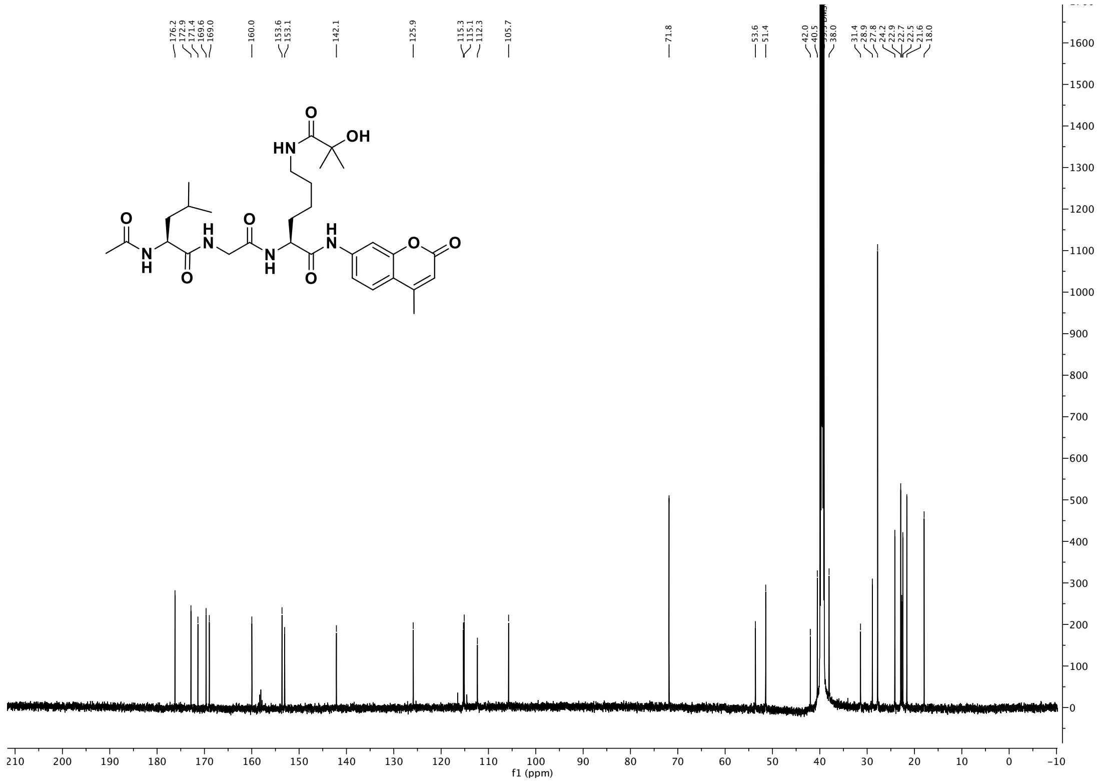

[13]C NMR spectra (151 MHz, DMSO) of compound **2o**

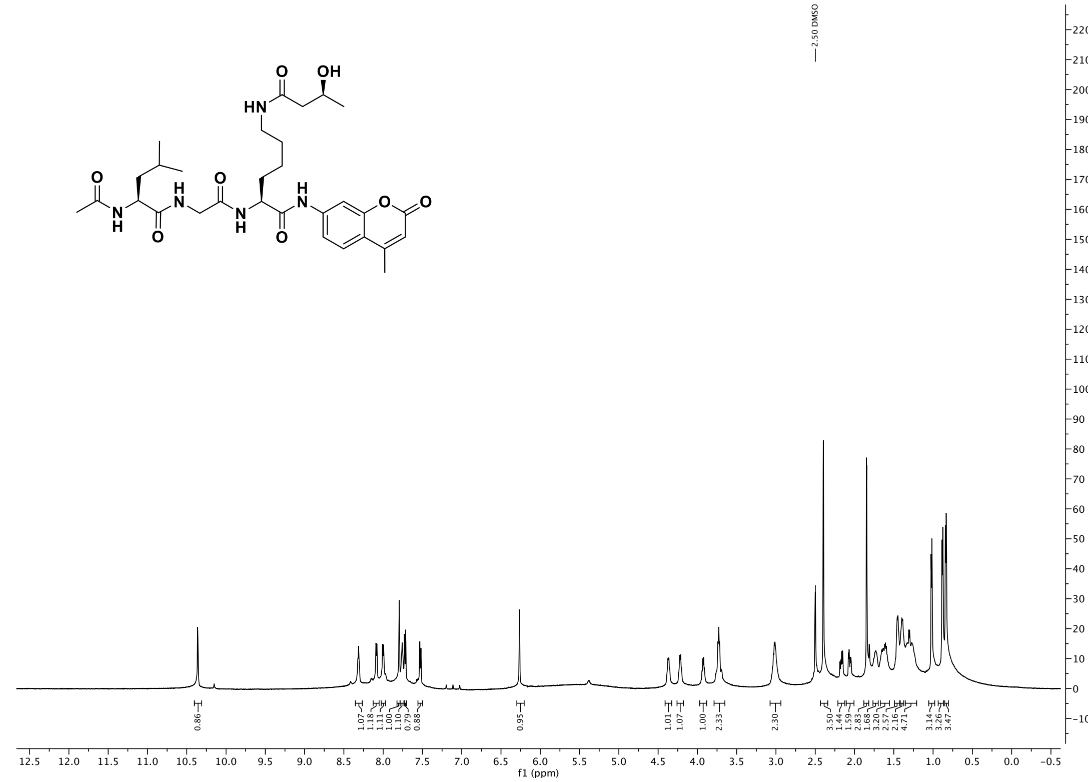

$^{1}H$ NMR spectra (600 MHz, DMSO-$d_6$) of compound **2p**

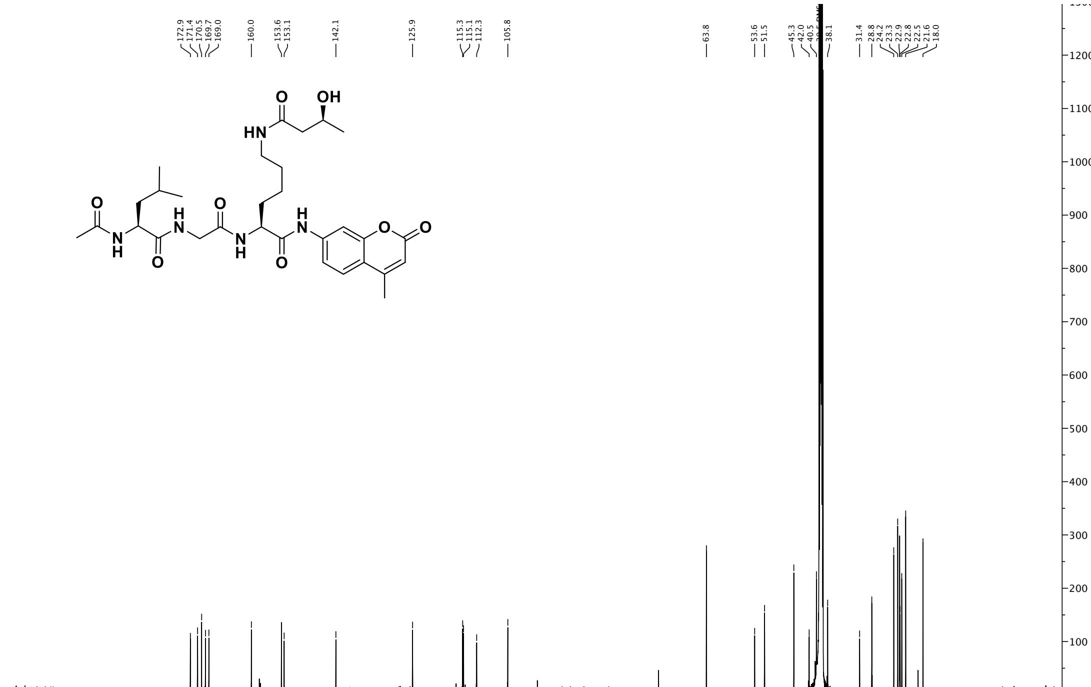

$^{13}C$ NMR spectra (151 MHz, DMSO) of compound **2p**

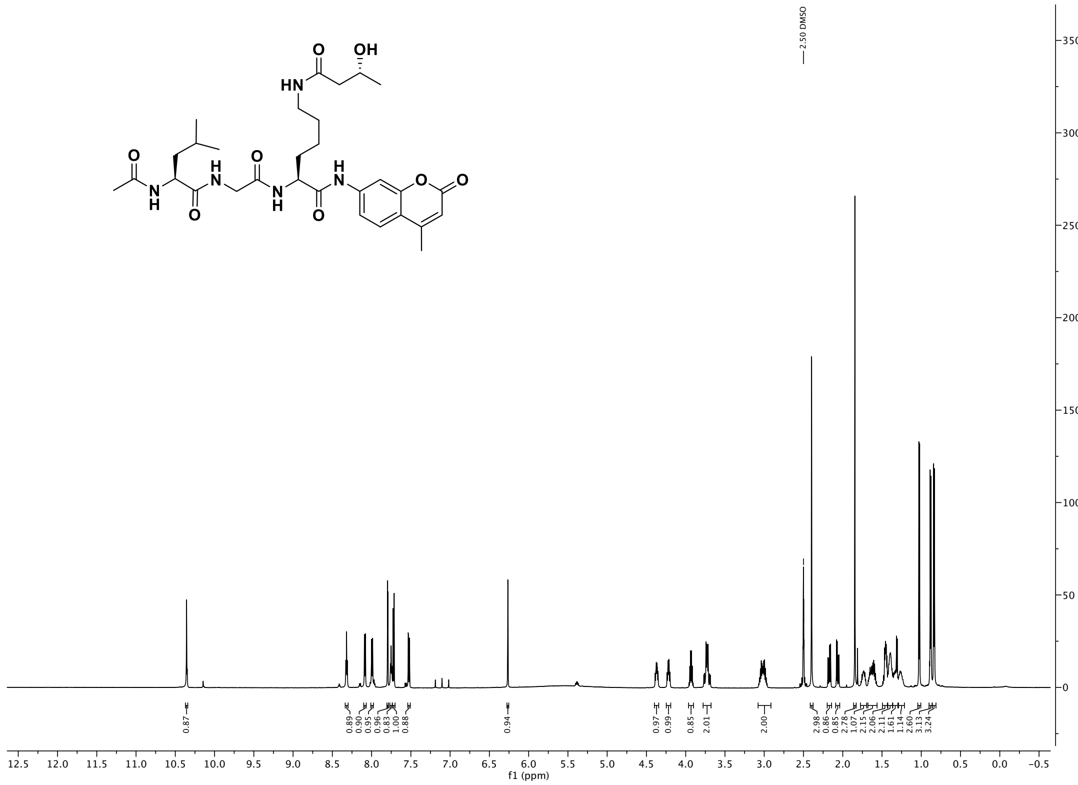

$^1$H NMR spectra (600 MHz, DMSO-$d_6$) of compound **2q**

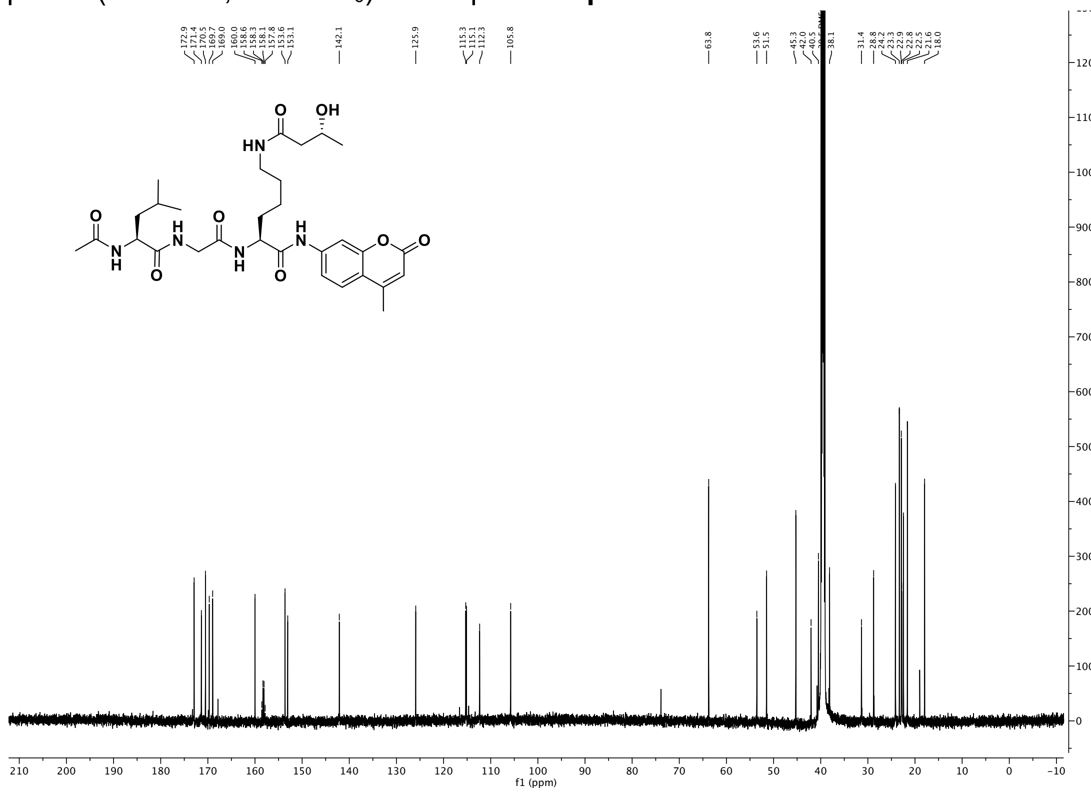

$^{13}$C NMR spectra (151 MHz, DMSO) of compound **2q**

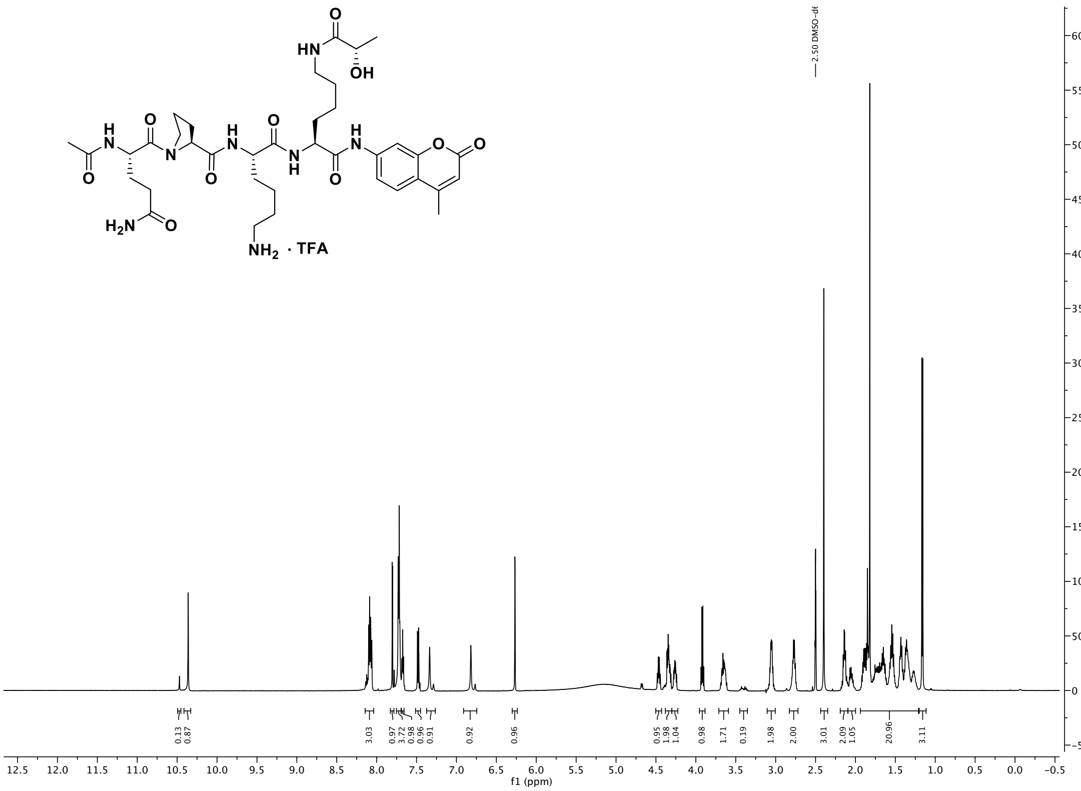

$^{1}$H NMR spectra (600 MHz, DMSO-$d_6$) of compound **4b**

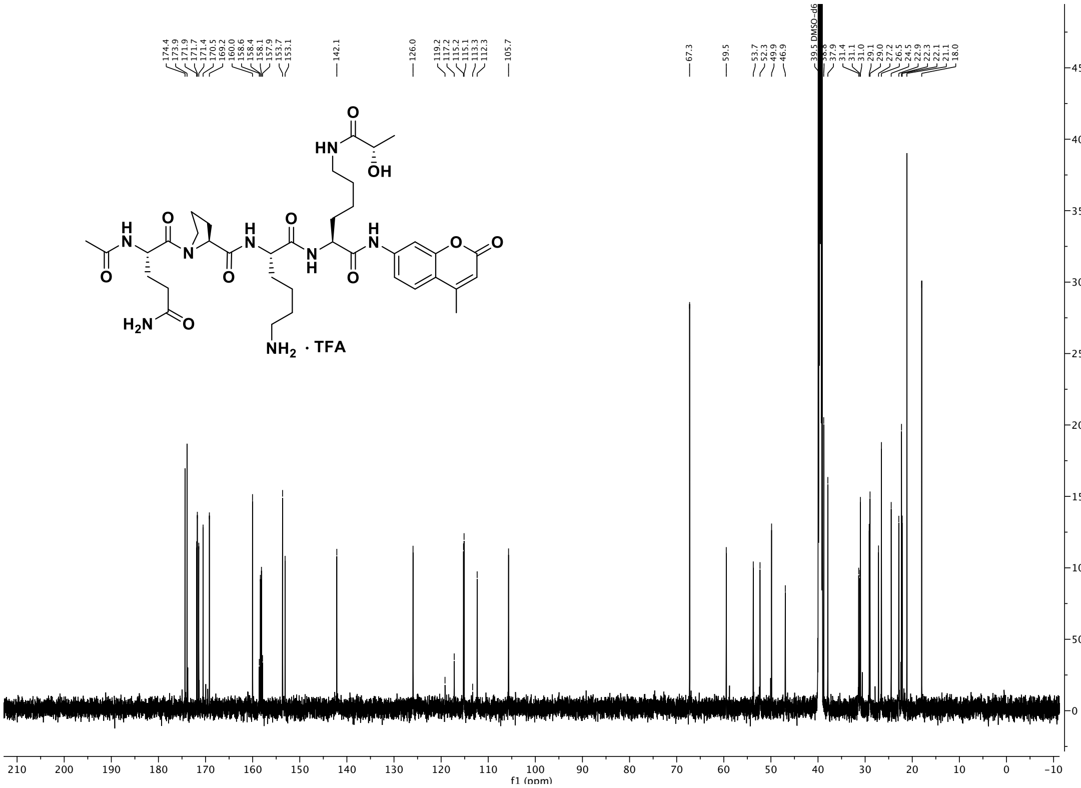

$^{13}$C NMR spectra (151 MHz, DMSO) of compound **4b**

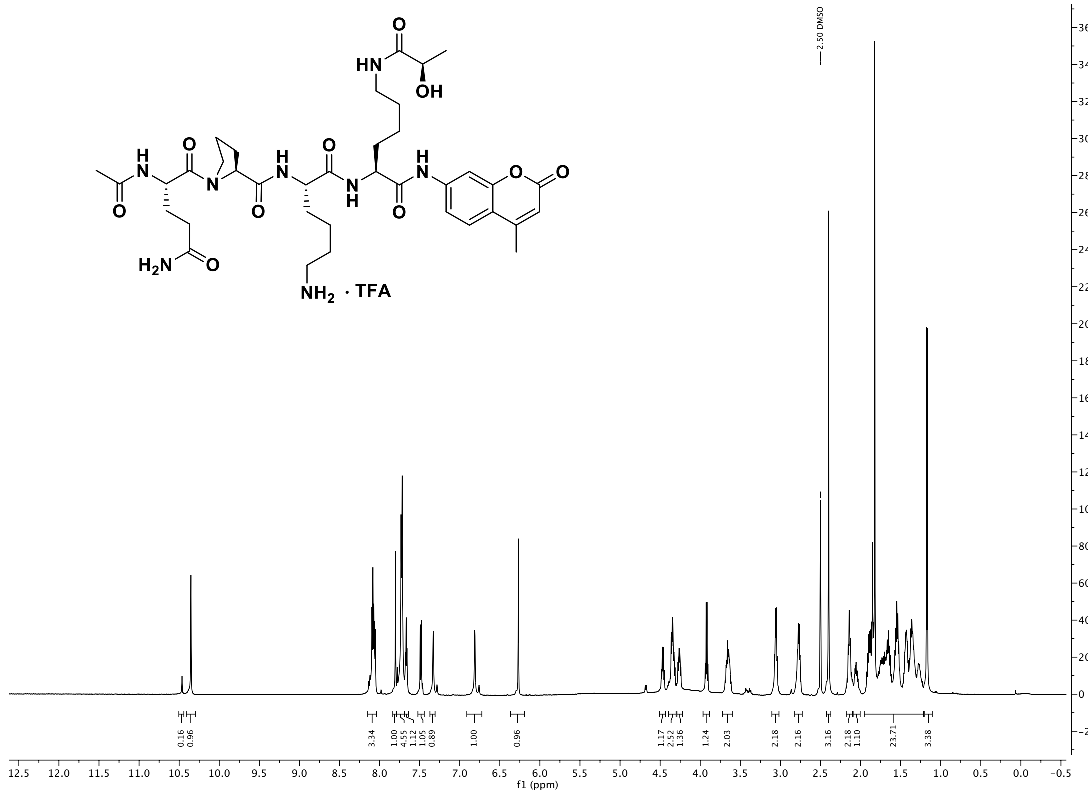

$^{1}$H NMR spectra (600 MHz, DMSO-$d_6$) of compound **4c**

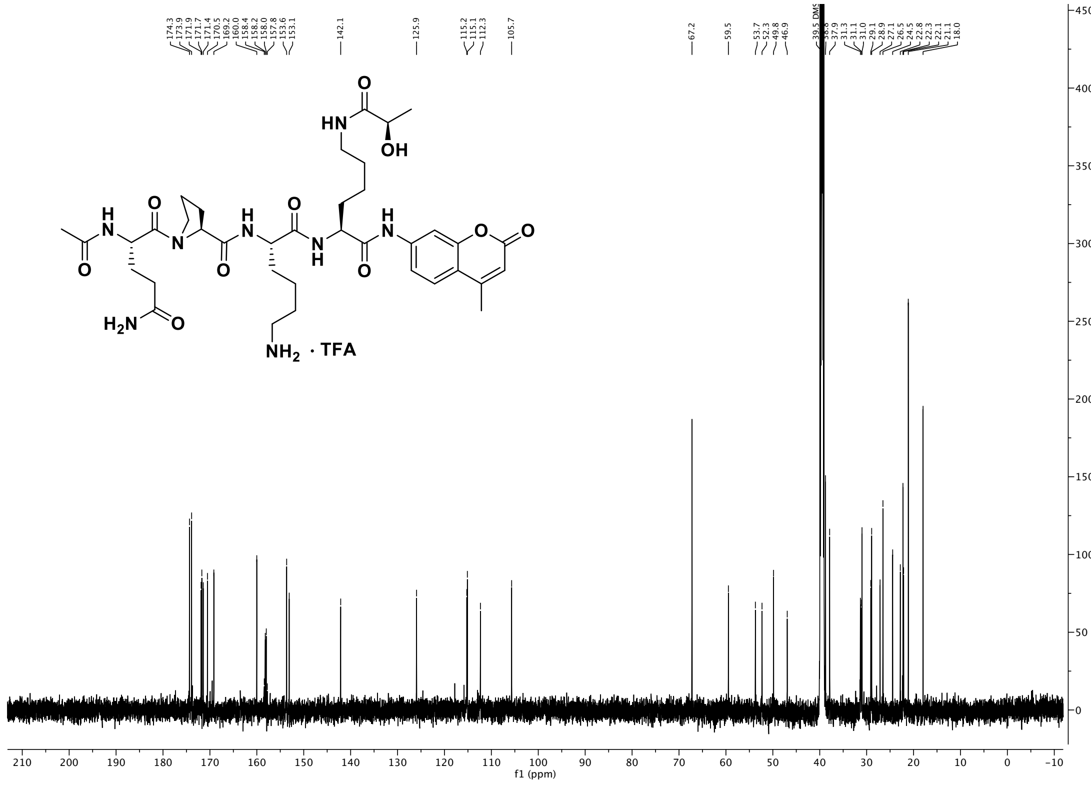

$^{13}$C NMR spectra (151 MHz, DMSO) of compound **4c**

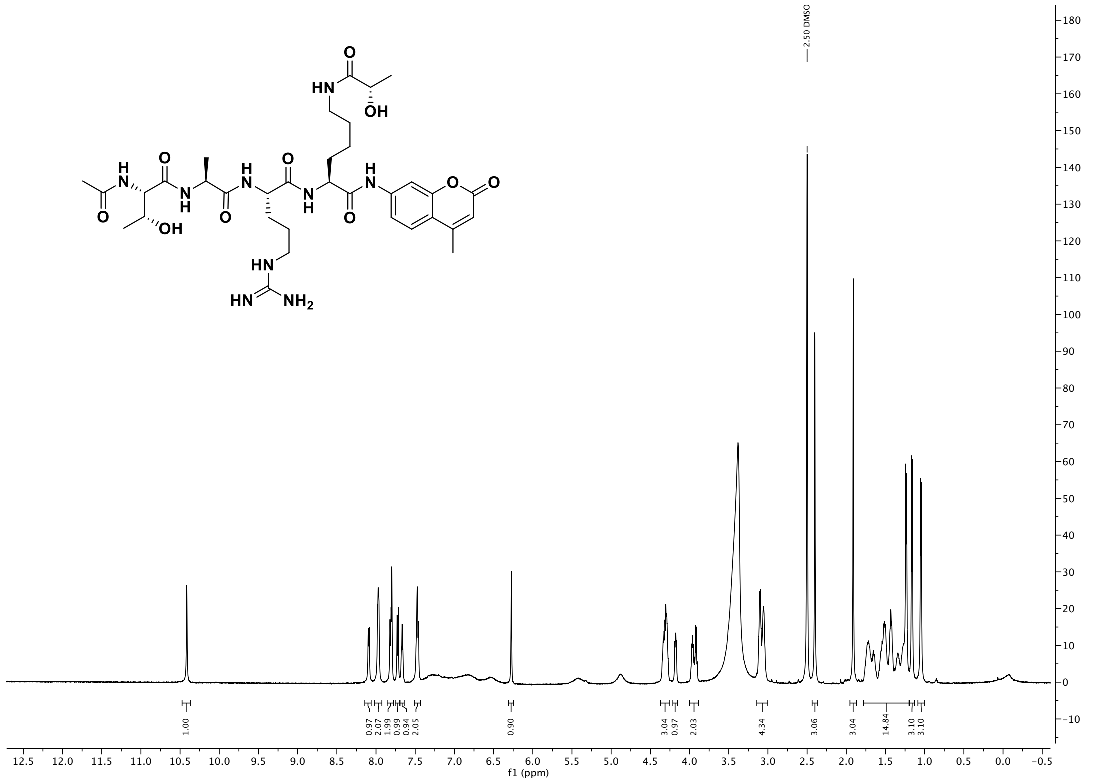

$^{1}$H NMR spectra (600 MHz, DMSO-$d_6$) of compound **5b**

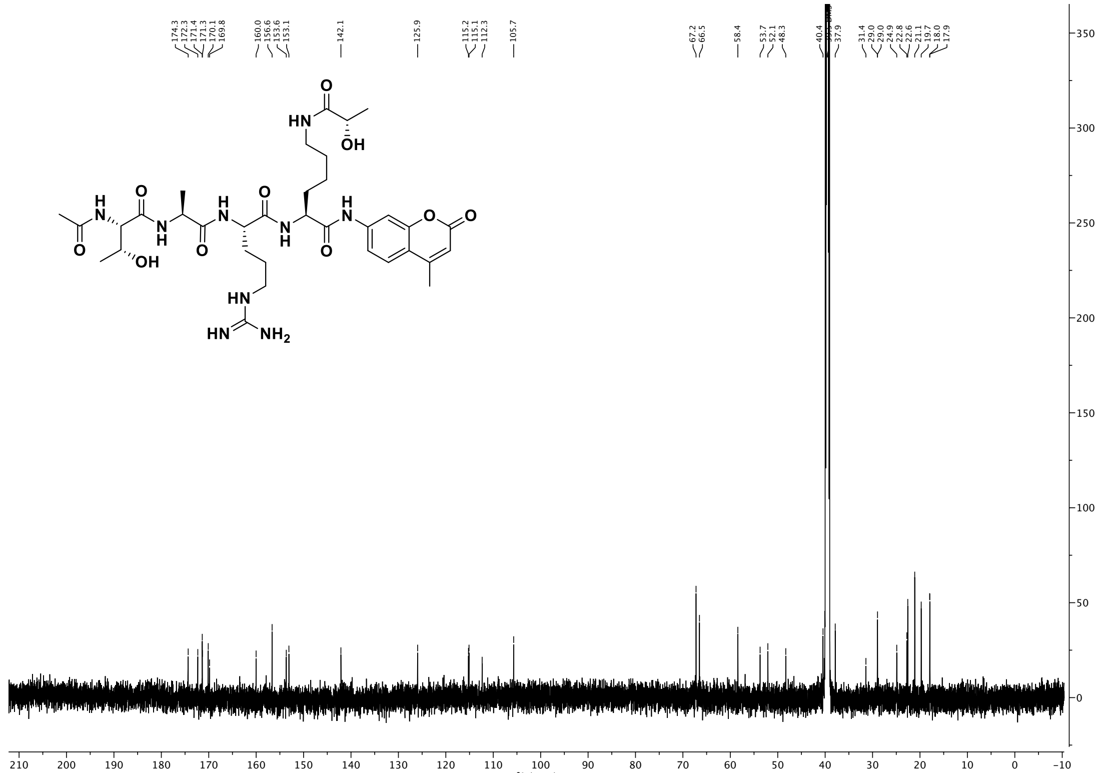

$^{13}$C NMR spectra (151 MHz, DMSO) of compound **5b**

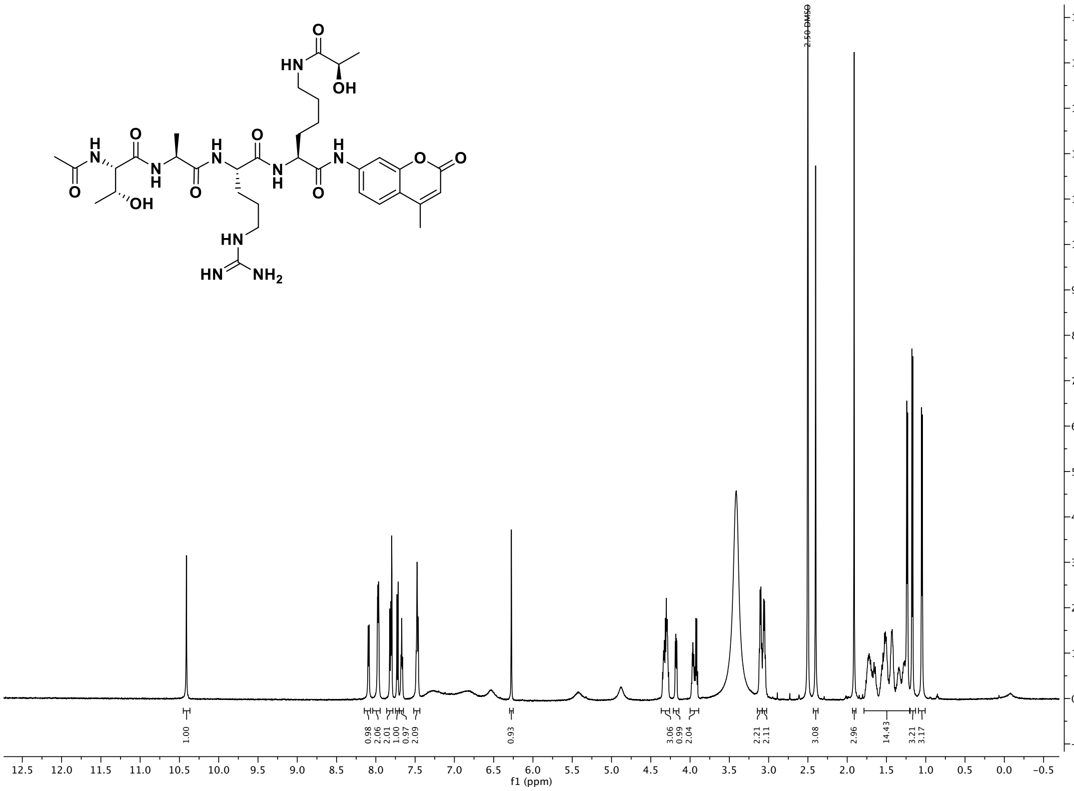

$^{1}$H NMR spectra (600 MHz, DMSO-$d_6$) of compound **5c**

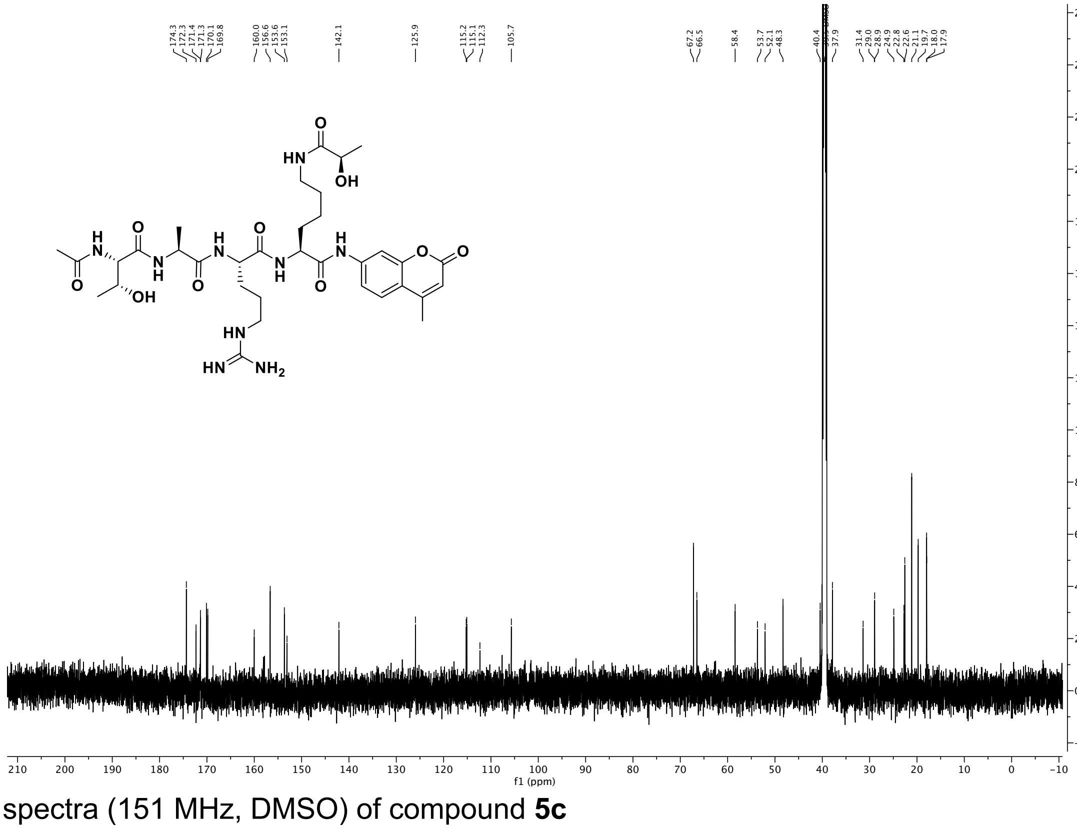

$^{13}$C NMR spectra (151 MHz, DMSO) of compound **5c**

*HPLC purity traces*

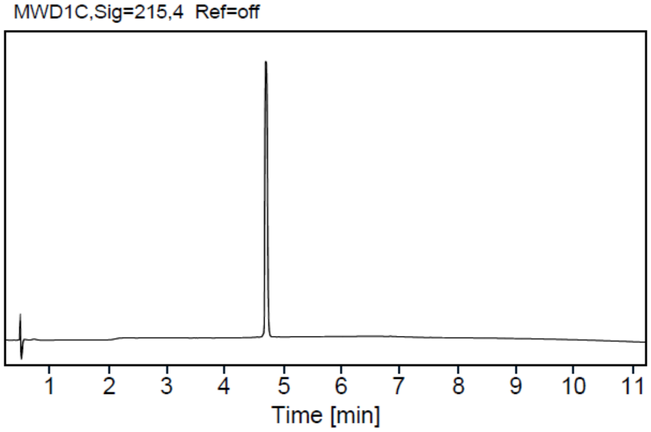


HPLC trace of peptide **5a**

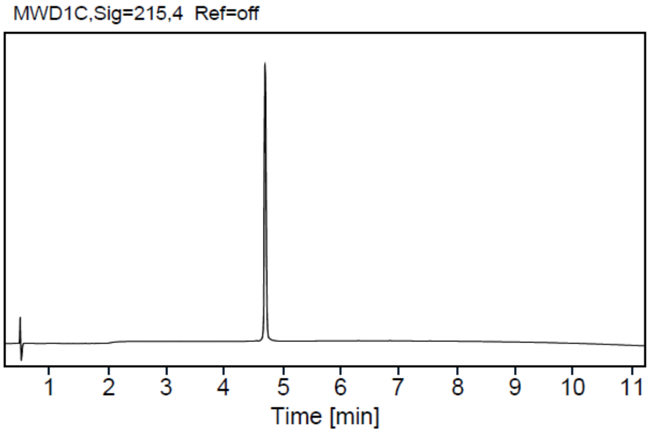


HPLC trace of peptide **5b**

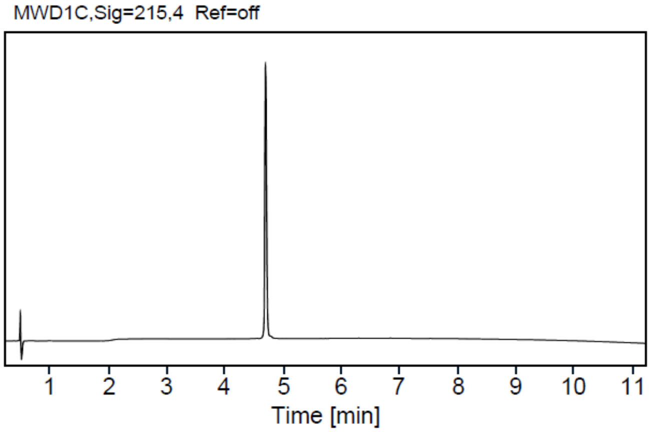


HPLC trace of peptide **5c**

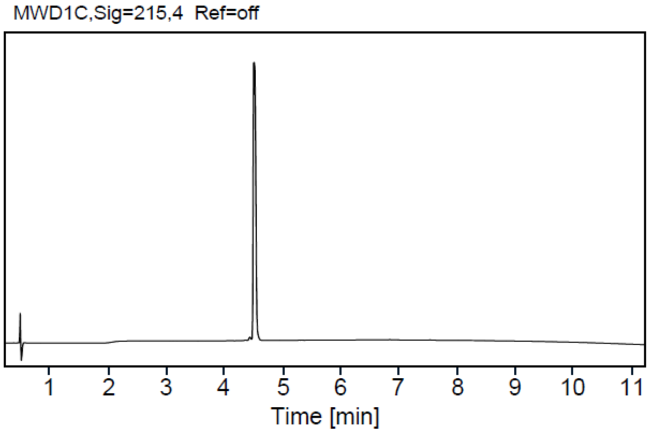


HPLC trace of peptide **5d**

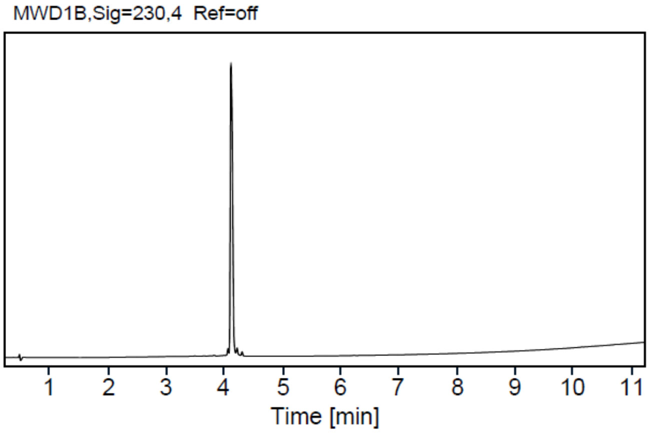


HPLC trace of peptide **6a**

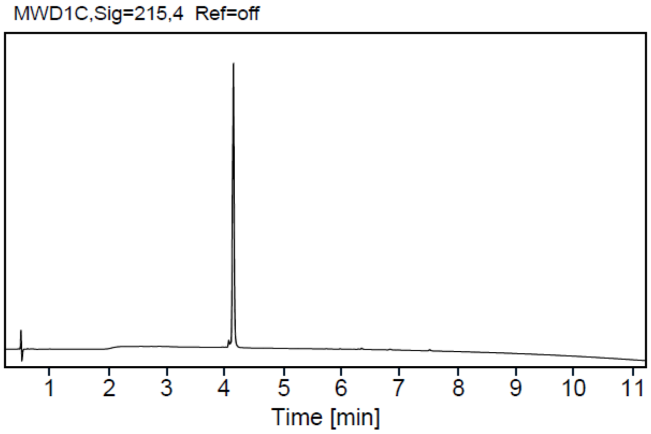


HPLC trace of peptide **6b**

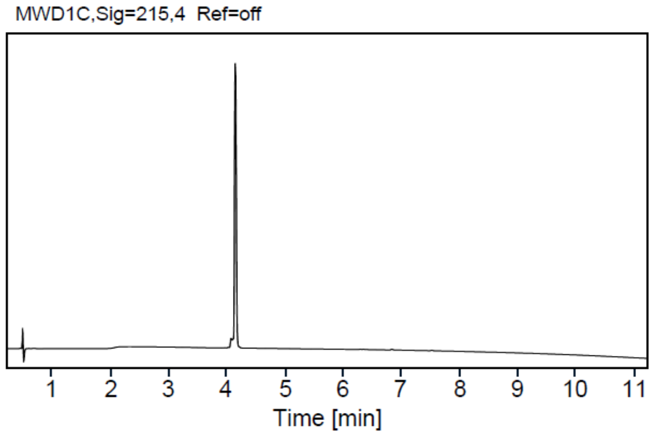

HPLC trace of peptide **6c**

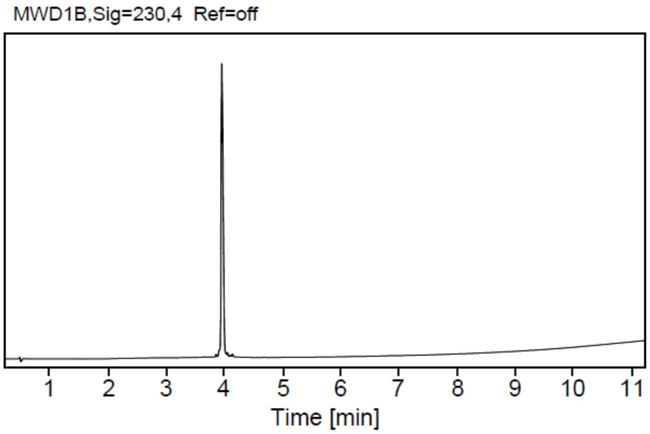

HPLC trace of peptide **6d**

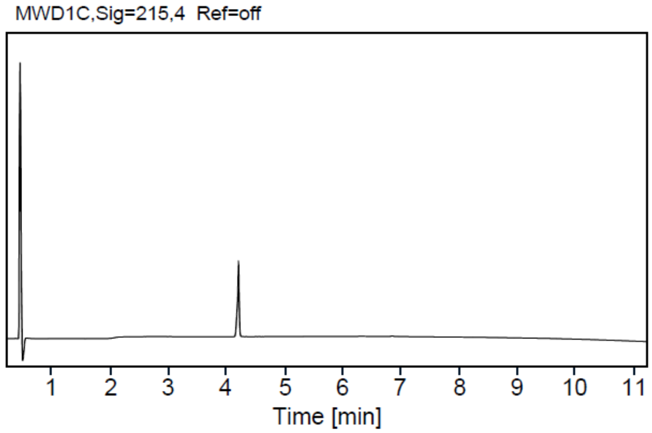

HPLC trace of peptide **7a**

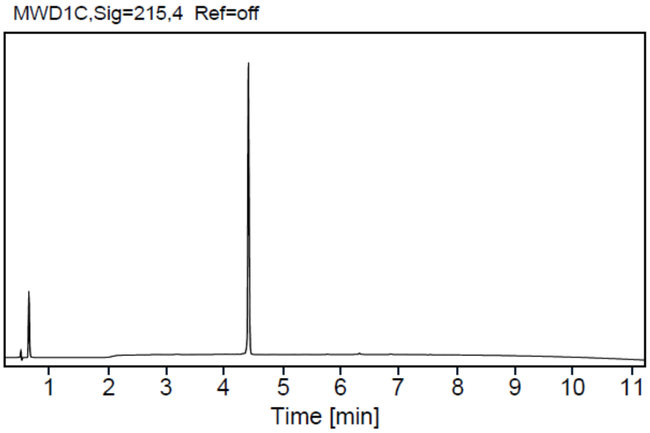

HPLC trace of peptide **7b**

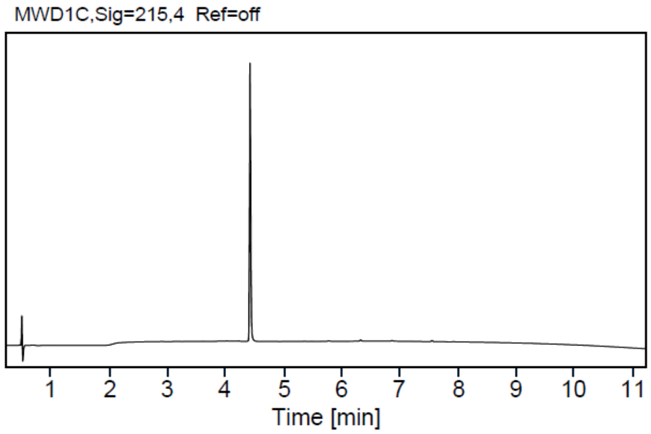

HPLC trace of peptide **7c**

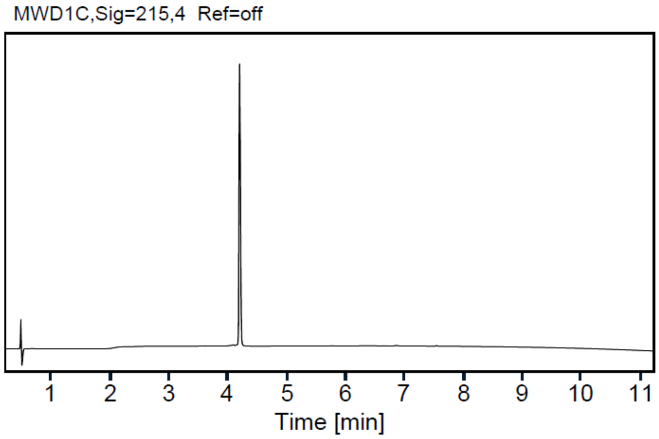

HPLC trace of peptide **7d**

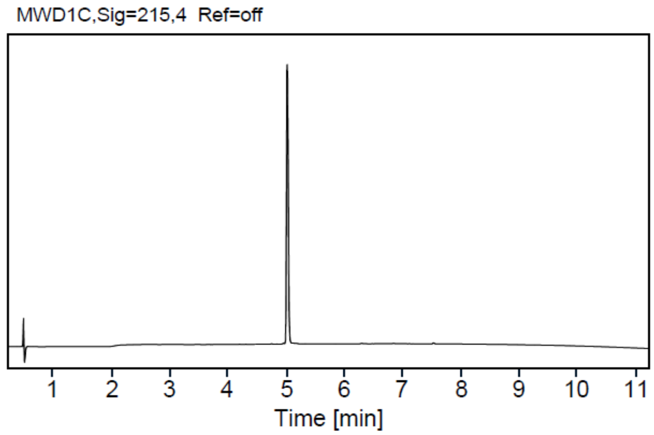

HPLC trace of peptide **8a**

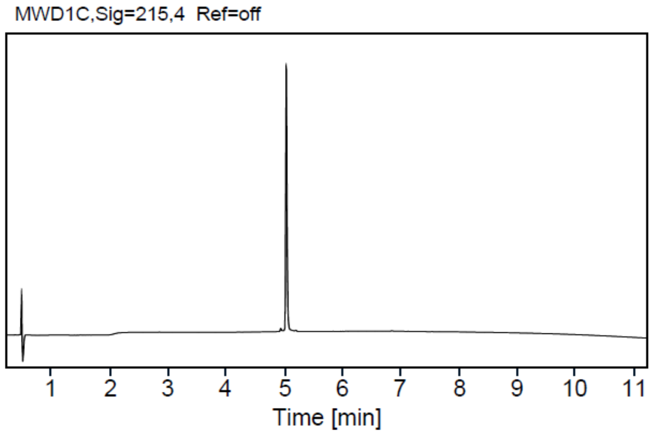

HPLC trace of peptide **8b**

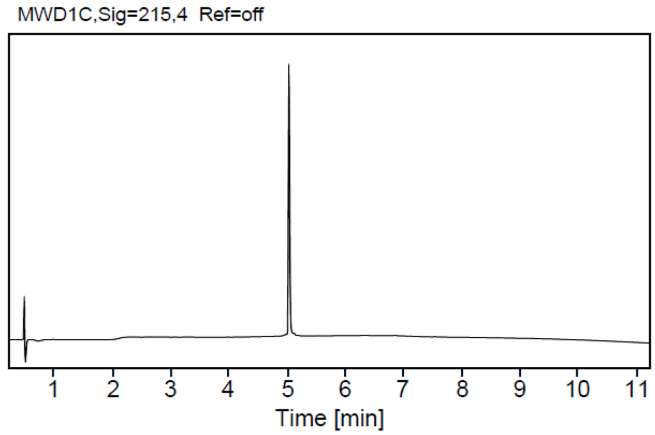

HPLC trace of peptide **8c**

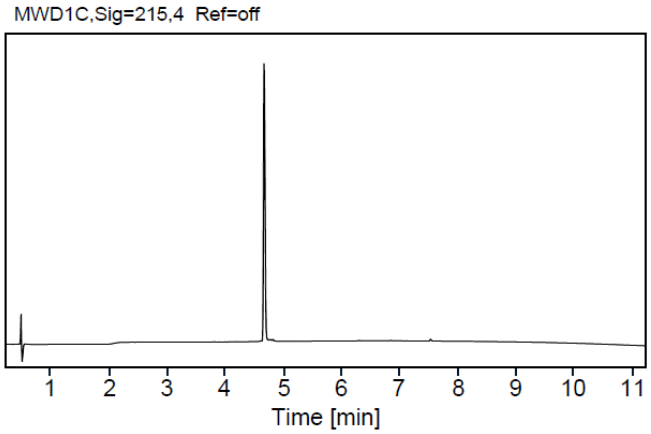

HPLC trace of peptide **8d**

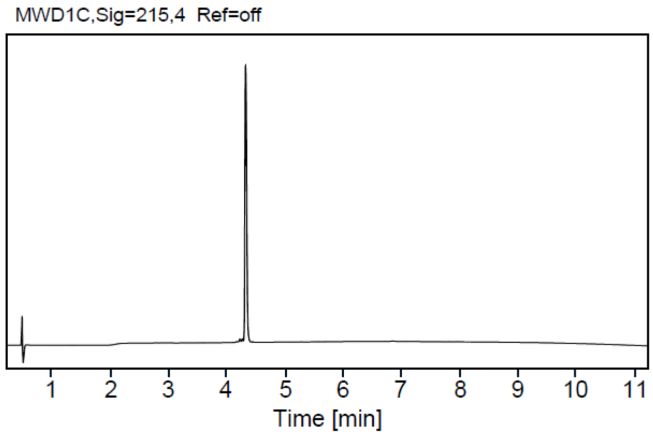

HPLC trace of peptide **9a**

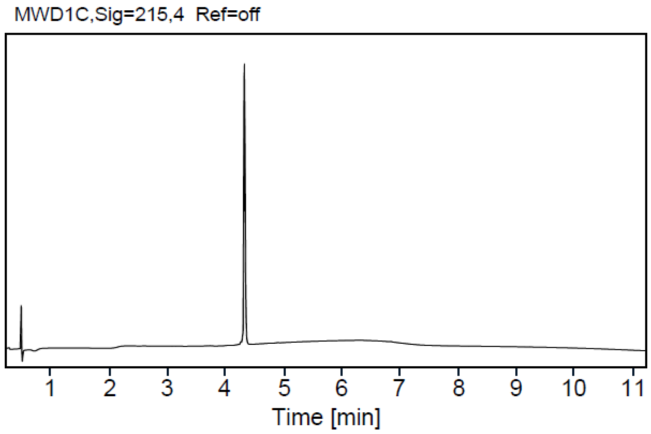

HPLC trace of peptide **9b**

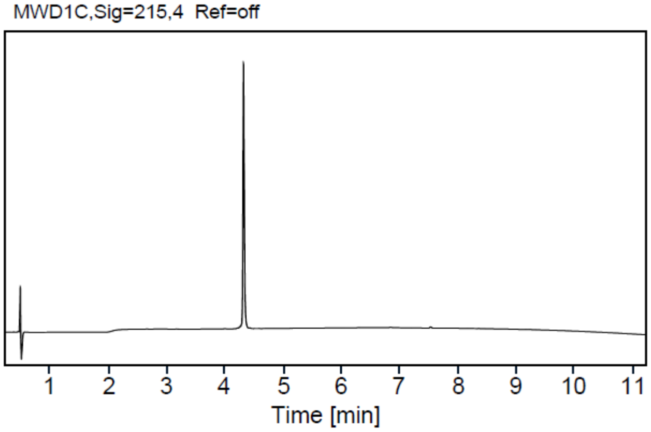


HPLC trace of peptide **9c**

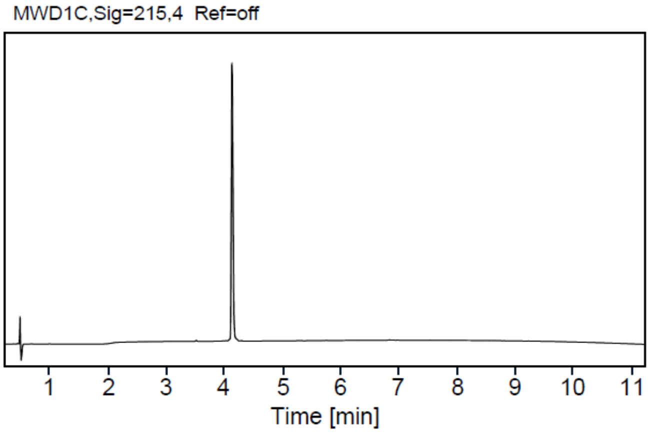


HPLC trace of peptide **9d**

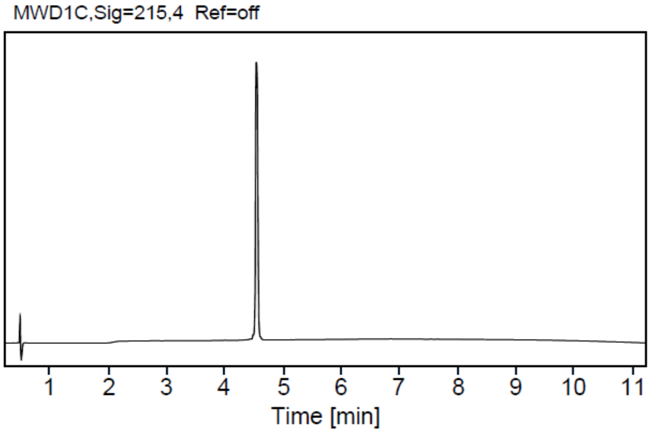


HPLC trace of peptide **10a**

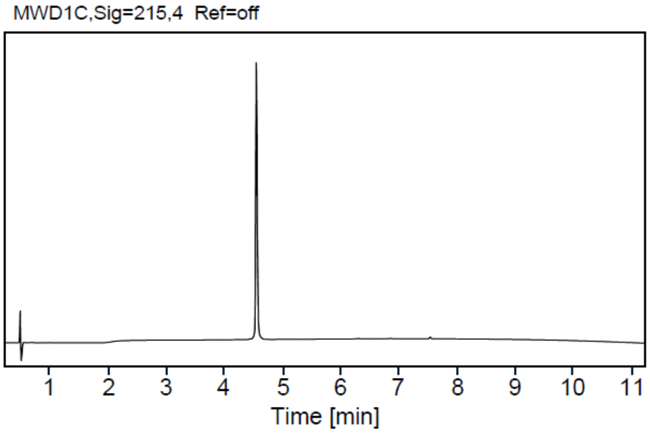


HPLC trace of peptide **10b**

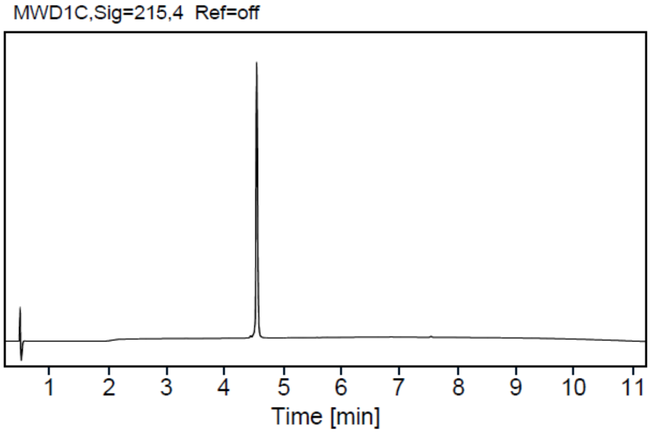


HPLC trace of peptide **10c**

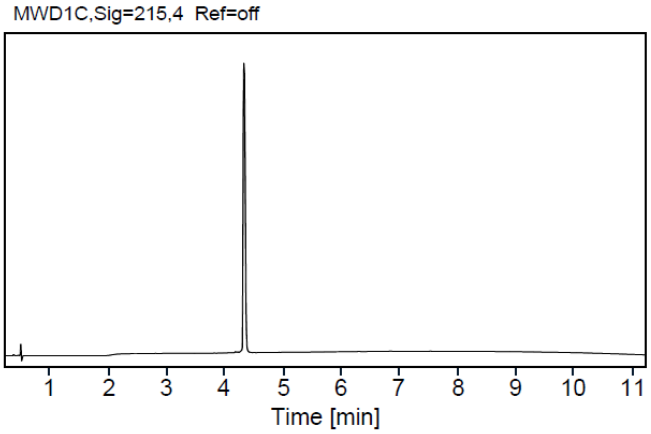


HPLC trace of peptide **10d**
